# Temporal dynamics of the multi-omic response to endurance exercise training across tissues

**DOI:** 10.1101/2022.09.21.508770

**Authors:** MoTrPAC Study Group, David Amar, Nicole R. Gay, Pierre M. Jean Beltran, Joshua N. Adkins, Jose J. Almagro Armenteros, Euan Ashley, Julian Avila-Pacheco, Dam Bae, Nasim Bararpour, Charles Burant, Clary Clish, Gary Cutter, Surendra Dasari, Courtney Dennis, Charles R. Evans, Facundo M. Fernández, David Gaul, Yongchao Ge, Robert Gerszten, Laurie J. Goodyear, Zhenxin Hou, Olga Ilkayeva, Anna A. Ivanova, David Jimenez-Morales, Maureen T. Kachman, Hasmik Keshishian, William E. Kraus, Ian R. Lanza, Jun Li, Malene E. Lindholm, Ana C. Lira, Gina M. Many, Shruti Marwaha, Michael E. Miller, Michael J. Muehlbauer, K. Sreekumaran Nair, Venugopalan D. Nair, Archana Natarajan Raja, Christopher Newgard, Eric A. Ortlund, Paul D. Piehowski, David M. Presby, Wei-Jun Qian, Jessica L. Rooney, James A. Sanford, Evan Savage, Stuart C. Sealfon, Gregory R. Smith, Kevin S. Smith, Alec Steep, Cynthia L. Stowe, Yifei Sun, Russell Tracy, Nikolai G. Vetr, Martin J. Walsh, Si Wu, Tiantian Zhang, Bingqing Zhao, Jimmy Zhen, Brent G. Albertson, Mary Anne S. Amper, Ali Tugrul Balci, Marcas Bamman, Elisabeth R. Barton, Bryan Bergman, Daniel Bessesen, Frank Booth, Brian Bouverat, Thomas W. Buford, Tiziana Caputo, Toby L. Chambers, Clarisa Chavez, Maria Chikina, Roxanne Chiu, Michael Cicha, Paul M. Coen, Dan Cooper, Elaine Cornell, Karen P. Dalton, Luis Oliveria De Sousa, Roger Farrar, Kishore Gadde, Nicole Gagne, Bret H. Goodpaster, Marina A. Gritsenko, Kristy Guevara, Fadia Haddad, Joshua R. Hansen, Melissa Harris, Trevor Hastie, Krista M. Hennig, Steven G. Hershman, Andrea Hevener, Michael F. Hirshman, Fang-Chi Hsu, Kim M. Huffman, Chia-Jui Hung, Chelsea Hutchinson-Bunch, Bailey E. Jackson, Catherine Jankowski, Christopher A. Jin, Neil M. Johannsen, Benjamin G. Ke, Wendy M. Kohrt, Kyle S. Kramer, Christiaan Leeuwenburgh, Sarah J. Lessard, Bridget Lester, Xueyun Liu, Ching-ju Lu, Nathan S. Makarewicz, Kristal M. Maner-Smith, DR Mani, Nada Marjanovic, Andrea Marshall, Sandy May, Edward Melanson, Matthew E. Monroe, Ronald J. Moore, Samuel Moore, Kerrie L. Moreau, Charles C. Mundorff, Nicolas Musi, Daniel Nachun, Michael D. Nestor, Robert L. Newton, Barbara Nicklas, Pasquale Nigro, German Nudelman, Marco Pahor, Cadence Pearce, Vladislav A. Petyuk, Hanna Pincas, Scott Powers, Shlomit Radom-Aizik, Krithika Ramachandran, Megan E. Ramaker, Irene Ramos, Tuomo Rankinen, Alexander (Sasha) Raskind, Blake B. Rasmussen, Eric Ravussin, R. Scott Rector, W. Jack Rejeski, Collyn Richards, Stas Rirak, Jeremy M. Robbins, Aliza B. Rubenstein, Frederique Ruf-Zamojski, Scott Rushing, Tyler J. Sagendorf, Mihir Samdarshi, Irene E. Schauer, Robert Schwartz, Nitish Seenarine, Tanu Soni, Lauren M. Sparks, Christopher Teng, Anna Thalacker-Mercer, John Thyfault, Rob Tibshirani, Scott Trappe, Todd A. Trappe, Karan Uppal, Sindhu Vangeti, Mital Vasoya, Elena Volpi, Alexandria Vornholt, Michael P. Walkup, John Williams, Ashley Xia, Zhen Yan, Xuechen Yu, Chongzhi Zang, Elena Zaslavsky, Navid Zebarjadi, Sue C. Bodine, Steven Carr, Karyn Esser, Stephen B. Montgomery, Simon Schenk, Michael P. Snyder, Matthew T. Wheeler

## Abstract

Regular exercise promotes whole-body health and prevents disease, yet the underlying molecular mechanisms throughout a whole organism are incompletely understood. Here, the Molecular Transducers of Physical Activity Consortium (MoTrPAC) profiled the temporal transcriptome, proteome, metabolome, lipidome, phosphoproteome, acetylproteome, ubiquitylproteome, epigenome, and immunome in whole blood, plasma, and 18 solid tissues in *Rattus norvegicus* over 8 weeks of endurance exercise training. The resulting data compendium encompasses 9466 assays across 19 tissues, 25 molecular platforms, and 4 training time points in young adult male and female rats. We identified thousands of shared and tissue- and sex-specific molecular alterations. Temporal multi-omic and multi-tissue analyses demonstrated distinct patterns of tissue remodeling, with widespread regulation of immune, metabolism, heat shock stress response, and mitochondrial pathways. These patterns provide biological insights into the adaptive responses to endurance training over time. For example, exercise training induced heart remodeling via altered activity of the *Mef2* family of transcription factors and tyrosine kinases. Translational analyses revealed changes that are consistent with human endurance training data and negatively correlated with disease, including increased phospholipids and decreased triacylglycerols in the liver. Sex differences in training adaptation were widespread, including those in the brain, adrenal gland, lung, and adipose tissue. Integrative analyses generated novel hypotheses of disease relevance, including candidate mechanisms that link training adaptation to non-alcoholic fatty liver disease, inflammatory bowel disease, cardiovascular health, and tissue injury and recovery. The data and analysis results presented in this study will serve as valuable resources for the broader community and are provided in an easily accessible public repository (https://motrpac-data.org/).

**Highlights:**

- Multi-tissue resource identifies 35,439 analytes regulated by endurance exercise training at 5% FDR across 211 combinations of tissues and molecular platforms.
- Interpretation of systemic and tissue-specific molecular adaptations produced hypotheses to help describe the health benefits induced by exercise.
- Robust sex-specific responses to endurance exercise training are observed across multiple organs at the molecular level.
- Deep multi-omic profiling of six tissues defines regulatory signals for tissue adaptation to endurance exercise training.
- All data are available in a public repository, and processed data, analysis results, and code to reproduce major analyses are additionally available in convenient R packages.

## Introduction

Regular exercise provides wide-ranging health benefits, including substantially reduced risk of all-cause mortality^1,2^ and protection against cardiometabolic and neurological diseases, cancer, and other pathologies^3–5^. Exercise affects nearly all organ systems in either improving health or in reducing disease risk^3–6^. The benefits of regular exercise result from cellular and molecular adaptations across various tissues and organ systems^6^.

Various “omic” platforms, including transcriptomics, epigenomics, proteomics, and metabolomics, have been used to study these events. These studies identified, for example, structural and biochemical alterations in skeletal muscle^7,8^, glycolytic-related remodeling in heart^9^, immune and cell cycle adaptation processes in blood^10–12^, and changes of gene regulation architecture in adipose tissue^13^. With technological advances, additional omic platforms have been applied to study exercise training adaptations. For example, phosphoproteomic studies have identified critical kinases in both human and rodent muscle^14–16^, and combined with acetylomic and metabolomic analysis, have identified a link between protein acetylation and fatty acid oxidation^17^. However, extant work typically covers one or two omes at a single time point, is biased towards one sex, and often focuses on a single tissue, mostly skeletal muscle, heart, or blood, e.g., ^7,9,10,17,18^, with few studies considering other tissues, e.g., ^19^. Accordingly, a comprehensive, organism-wide, multi-omic map of the effects of exercise is needed to understand the molecular underpinnings of exercise training-induced adaptations.

To address this need, the Molecular Transducers of Physical Activity Consortium (MoTrPAC) was established with the goal of building a molecular map of the exercise responses across a broad range of tissues in animal models and humans^20^. Here, we present the MoTrPAC animal endurance exercise training study conducted in young adult male and female rats. The rat is preferred over the mouse as a model organism for endurance exercise^21,22^, as glucose metabolism and cardiac responses are more similar to humans^23,24^, and their large tissue masses enable extensive multi-omic interrogation. Altogether, this work presents the first whole-organism molecular map of the temporal effects of endurance exercise training and demonstrates multiple insights enabled by this multi-omic data resource.

## Results

### Multi-omic approach characterizes the endurance exercise training responses

Six-month-old male and female Fischer 344 rats were subjected to progressive endurance exercise training (hereon referred to as “endurance training”) on a motorized treadmill for 1, 2, 4, or 8 weeks, with tissues collected 48 hours after the last training bout (Figure 1A). Sex-matched sedentary, untrained rats were used as controls (Fig. 1A); all animals were reverse light/dark cycle adapted. Training resulted in robust phenotypic changes (Figure S1A-D), including significantly increased maximal aerobic capacity (VO_2_max) by 18% and 14% at 8 weeks in males and females, respectively (q-value < 0.01), with no change in controls (Fig. S1A). Nuclear magnetic resonance spectroscopy showed that the absolute percentage of body fat decreased by 5% in males at 8 weeks (Fig. S1B), without a significant change in lean mass (Fig. S1C). In females, body fat percentage did not change after 4 or 8 weeks of training, while sedentary control females increased body fat by 4% (Fig. S1B). Females showed increased body weight consistent with growth over all intervention groups while males showed no change in body weight (Fig. S1D).

**Figure 1.**
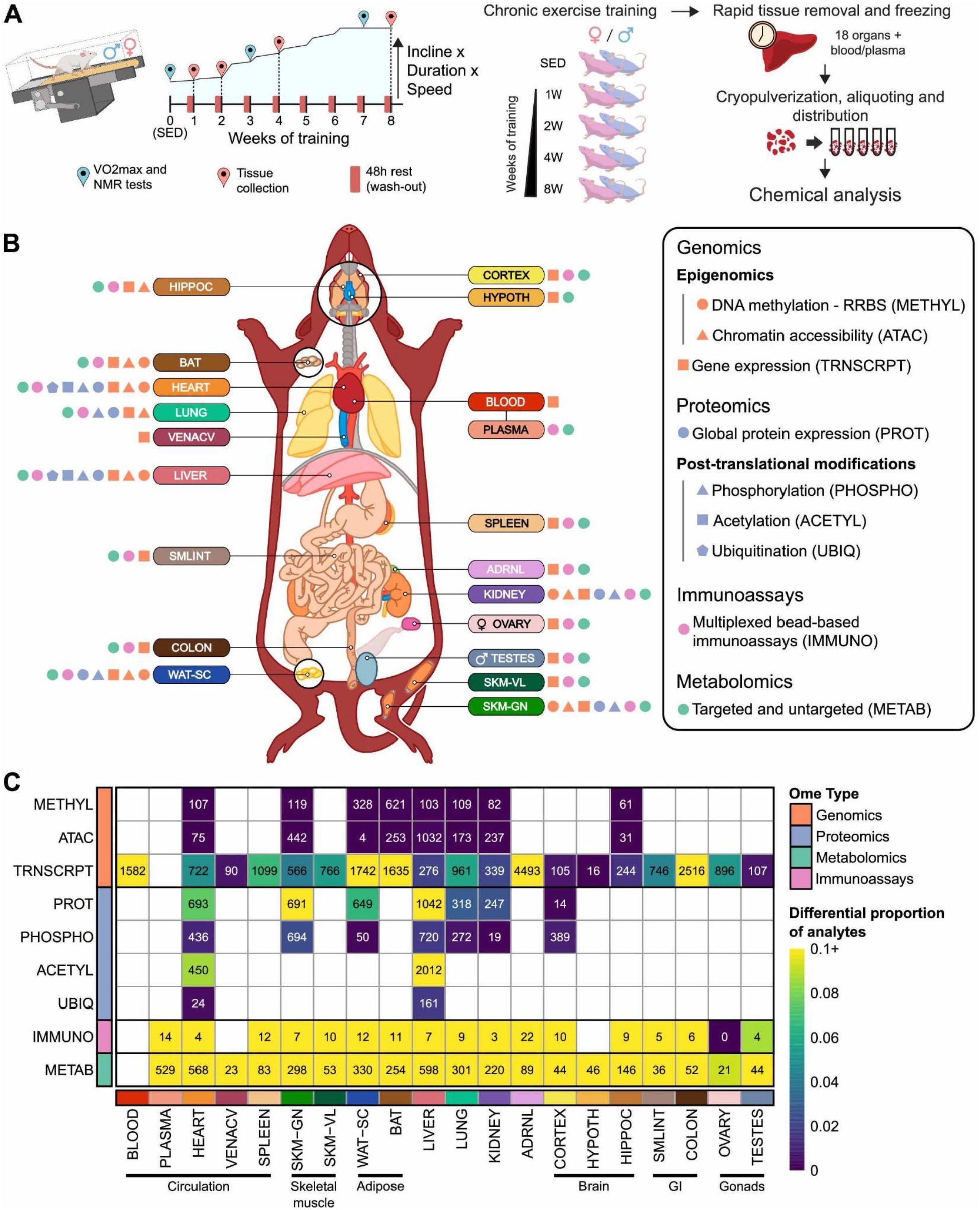
Study design and the dataset. **A)** Experimental design and tissue sample processing. Fischer 344 (F344) inbred rats were subjected to a progressive treadmill training protocol. Tissues were collected from male and female animals that remained sedentary (SED) or completed 1, 2, 4, or 8 weeks of endurance exercise training. For trained animals, samples were collected 48 hours after their last exercise bout (red pins). Maximal oxygen consumption (VO_2_max) and nuclear magnetic resonance (NMR) tests are indicated by the blue pins. SED = sedentary. **B)** Summary of molecular datasets included in this study. Up to nine data types were generated for blood, plasma, and 18 solid tissues, per animal. Tissue labels indicate the location, color code, and abbreviation for each tissue. Icons next to each tissue label indicate the molecular data types generated for that tissue. **C)** Number of differential molecular features at 5% false discovery rate (FDR). Each cell represents results for a single tissue and data type. Numbers indicate the number of training-differential features whose abundances significantly changed over the training time course in at least one sex; colors indicate the proportion of measured features that are differential, where the brightest color means that at least 10% of measured features are differential. Tissues are grouped by organ system. Data types are grouped by ome, as defined in **(B)**, and ordered to reflect the central dogma. Created in part with BioRender.com.

Whole blood, plasma, and 18 solid tissues were analyzed using genomics, proteomics, metabolomics, and protein immunoassay technologies, with most assays performed in a subset of these tissues (Figure 1B, Figure S1E). For all omic analyses, tissue samples were cryopulverized to provide homogenous samples (Fig. 1A). To reduce the effects of inter-animal variation on data integration, the omic assays were performed on tissues from the same individual animals whenever possible (Figure S1F). Molecular assays were prioritized based on available tissue quantity and biological relevance, with gastrocnemius skeletal muscle, heart, liver, and white adipose tissue having the most diverse set of molecular assays performed, followed by the kidney, lung, brown adipose tissue, and hippocampus (Fig. S1E). Transcriptomic data from RNA sequencing (RNA-Seq) were collected in all 18 solid tissues and whole blood (n=5 per sex and time point); epigenomic data from transposase-accessible chromatin using sequencing (ATAC-seq) and reduced representation bisulfite sequencing (RRBS) were collected in 8 tissues (n=5 per sex and time point). Proteomic data were generated by LC-MS/MS for 7 tissues (n=6 per sex and time point) using TMT-based quantification where each plex contained a common tissue reference to enable quantification across plexes (Figure S2A). Global proteome and phosphoproteome data were acquired in all 7 tissues, whereas acetylome (acetylated lysines; acetylsites) and ubiquitylome (K-ε-GG ubiquitin remnants; ubiquitylsites) were acquired from the liver and heart (Fig. 1B, Fig. S1E).

Metabolomic data were generated for plasma and all 18 solid tissues using up to seven targeted platforms, four untargeted metabolomic platforms, and two untargeted lipidomic platforms (n=5 per sex and time point). Multiplexed immunoassay panels were applied to 17 tissues to accurately quantify low-abundance proteins of biological relevance, including cytokines. These datasets were generated using five different rat panels (54 analytes total; n=3 per sex and time point). For each omic analysis, QC metrics were used to ensure the data were of high quality (Figure S2B-K; Methods). Altogether, datasets were generated from 9466 assays across 211 combinations of tissues and molecular platforms, quantifying a total of 213,689 and 2,799,307 unique non-epigenetic and epigenetic (RRBS/ATAC-seq) features, respectively, out of a total of 681,256 and 14,334,496 distinct non-epigenetic and epigenetic measurements, respectively.

Differential analysis was used to characterize the molecular responses to endurance training (e.g., negative binomial regression via DESeq2 for RNA-Seq count data; see Methods). For each molecular feature, a regression model was fit for each sex separately with the training group as an explaining variable. Meta-regression models were used for metabolites measured on more than one platform (see Methods). Using these models, we quantified the overall significance of the training response, denoted as the *training p-value*. We also computed eight sex- and time-specific pairwise contrasts between each group of trained animals and its sex-matched sedentary controls, denoted as the *timewise summary statistics*. To select molecular features with any training response, all training p-values within each omics data type (or “ome”) were adjusted for multiple hypothesis testing using Independent Hypothesis Weighting (IHW) with the corresponding tissue as a covariate^25^. The 35,439 features at 5% FDR comprise the *training-regulated differential features* (Figure 1C; Table S1).

Training-regulated molecules were observed in the vast majority of tissues for all omes, including a relatively large proportion of transcriptomics, proteomics, metabolomics, and immunoassay features (Fig. 1C). The observed fold-changes were modest, where 56% of the differential features had a maximum fold-change greater than 0.67 and less than 1.5. Among the tissues measured by transcriptomics, the hypothalamus, cortex, testes, and vena cava had the smallest proportion of training-regulated genes, whereas the blood, brown and white adipose tissues, adrenal gland, and colon showed broader transcriptional effects. For proteomics, the gastrocnemius, heart, and liver showed substantial differential regulation in both protein abundance and post-translational modifications, with more restricted differential regulation of white adipose tissue, lung, and kidney protein abundance. For metabolomics, a large proportion of differential metabolites were consistently observed across all tissues, although the absolute numbers were related to the number of metabolomic platforms employed (Fig. S1E). The number of differential features over the training time course across tissues and data types highlights the multi-layered organism-wide molecular adaptations to endurance training.

### Multi-tissue integration reveals system-wide molecular responses to endurance training

To identify genes that change across tissues during endurance training, we considered the lung, gastrocnemius, white adipose tissue, kidney, liver, and heart. These six tissues had data from the following assays: DNA methylation, chromatin accessibility, transcriptomics, global proteomics, phosphoproteomics, and multiplexed immunoassays. The 11,407 differential features from these datasets mapped to 7,115 unique genes across the six tissues (Figure 2A-D, Figure S3A; Table S2). These genes were tissue-specific (67%), with the greatest number appearing in white adipose tissue. However, 2,359 genes had differential features in at least two tissues. For these genes, we characterized the most common tissue combinations (Fig. 2A). The largest number of shared genes (m=249) corresponded to the lung and white adipose tissue, with predominantly immune-related pathway enrichments (q-value < 1e-05, Fig. 2B). The second-largest group of shared genes corresponded to the heart and gastrocnemius with enrichment of mitochondrial metabolism pathways (q-value < 1e-03, Fig. 2C).

**Figure 2.**
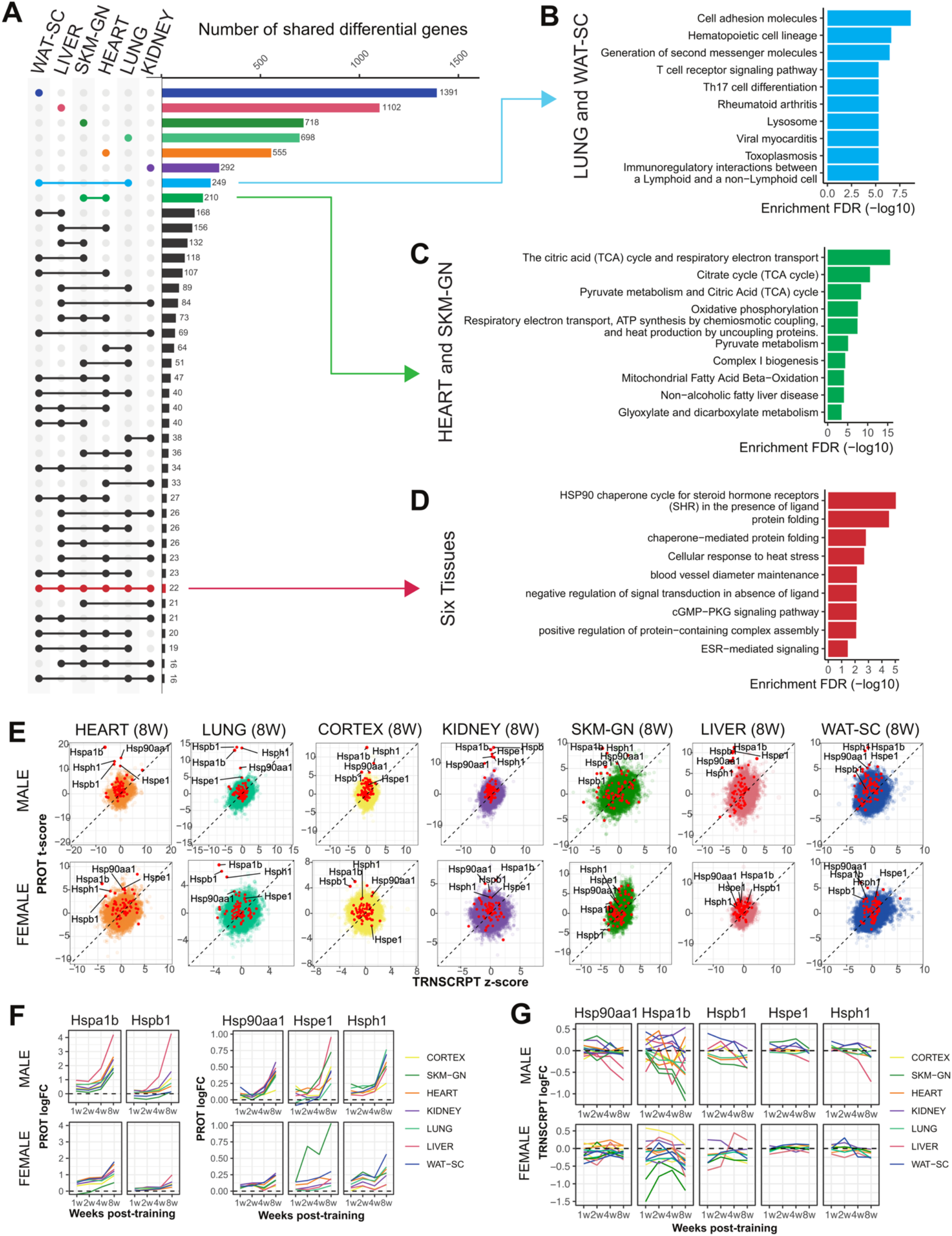
Multi-tissue molecular endurance training responses. **A)** Upset plot of the gene sets associated with the training response of each tissue. Molecular features were mapped to gene symbols. Numbers indicate the number of genes regulated by training in the tissues indicated by the connected points on the left. Bars and points indicating tissue-specific differential genes are colored by tissue. Pathway enrichment analysis is shown for selected sets of genes as indicated by the arrows. **B-D)** Significantly enriched pathways (10% FDR) corresponding to **(B)** genes differential in both LUNG and subcutaneous white adipose tissue (WAT-SC), **(C)** genes differential in both HEART and SKM-GN, and **(D)** the 22 genes that are training-regulated in all six tissues considered in **(A)**. Redundant pathways (i.e., those with an overlap of 80% or greater) were removed. **E)** Scatter plots of the protein t-scores (PROT) vs. the transcript z-scores (TRNSCRPT), per gene at 8 weeks of training (8W) relative to sedentary controls. Data are shown for the seven tissues for which both proteomics and transcriptomics was acquired. Red points indicate genes associated with the heat shock response, and the labeled points indicate those with a large differential response at the protein level. **F-G)** Line plots showing protein **(F)** and transcript **(G)** log_2_ fold-changes relative to the untrained controls for a subset of heat shock proteins with increased abundance during exercise training. Each line represents a protein in a single tissue.

Next, we compared the 8-week timewise summary statistics of the transcriptomics and protein abundance data for each tissue and sex combination (Figure 2E). This analysis revealed low to moderate correlation (Figure S3B). This is in agreement with numerous previous studies showing that exercise triggers a transient increase in transcript abundance, with a longer-lasting response at the protein level^26–28^. Nevertheless, gene set enrichment analysis (GSEA) revealed a greater concordance between these two omes (Figure S3C-D; Table S3-4; Methods).

The two analyses above suggested an up-regulated heat shock response across the organism. First, 22 genes were training-regulated in all six tissues, and this set was enriched in heat shock response pathways (q-value < 0.01, Fig. 2D). Second, multiple heat shock proteins (HSPs) appeared as outliers in our transcriptomics-proteomics comparison (Fig. 2E). The HSP family has important cytoprotective functions^29^, is known to be induced by exercise in various tissues, and is dysregulated in metabolic diseases like diabetes^30^. In humans, higher levels of the major HSPs HSPA1A/HSPA1B and HSP90A1AA (a.k.a HSP70 and HSP90, respectively) were previously observed in skeletal muscle of lifelong footballers when compared to untrained controls^31^ and were induced following states of increased protein turnover due to either muscle damage or states of increased myofiber turnover^32^. We found a robust increase in HSP abundance and virtually no change at the transcript level of the cognate genes, including subunits of the major HSPs *Hspa1b* and *Hsp90aa1*, as well as *Hspb1*, *Hspe1*, and *Hsph1* (Fig. 2E). The transcript and protein discordance suggests that a transient transcript induction or post-transcriptional regulation in response to each bout of exercise results in an accumulation of HSPs in response to long-term training. Our results also show a similar pattern of simultaneous cross-tissue inductions of HSPs in an intensity-or time-dependent manner only at the protein level, with a stronger response observed in males (Figure 2F-G).

Altogether, we find that endurance training induces a heat shock response and HSP accumulation across tissues, which can explain some of the cytoprotective effects associated with exercise^33,34^. The alteration of HSPs primarily at the protein level illustrates the importance of measuring multiple omes to fully understand the endurance training response. Moreover, our organism-wide analysis reveals overwhelmingly tissue-specific responses to endurance training.

### Heart and liver regulatory cascades are modulated by endurance training

To better understand the regulatory behavior behind the responses to endurance training, we inferred changes in activity of transcription factors (TFs) and phosphosignaling activities from the transcriptomic and proteomic data. TF activities were inferred by TF motif enrichment analysis via HOMER^35^, applied to the set of all differential transcripts in each tissue (see Methods). We isolated the most significantly enriched TFs and compared their enrichment levels as a proxy for TF activity regulation across tissues (Figure 3A, Figure S4A; Table S5). This revealed a cluster of enriched *Mef2* family TF motifs in the heart and skeletal muscle (Fig. 3A). MEF2C is a well-known muscle-associated TF and is involved in skeletal, cardiac, and smooth muscle cell differentiation^36^. Moreover, it has been implicated in vascular development, formation of the cardiac loop, and even neuron differentiation^37^.

**Figure 3.**
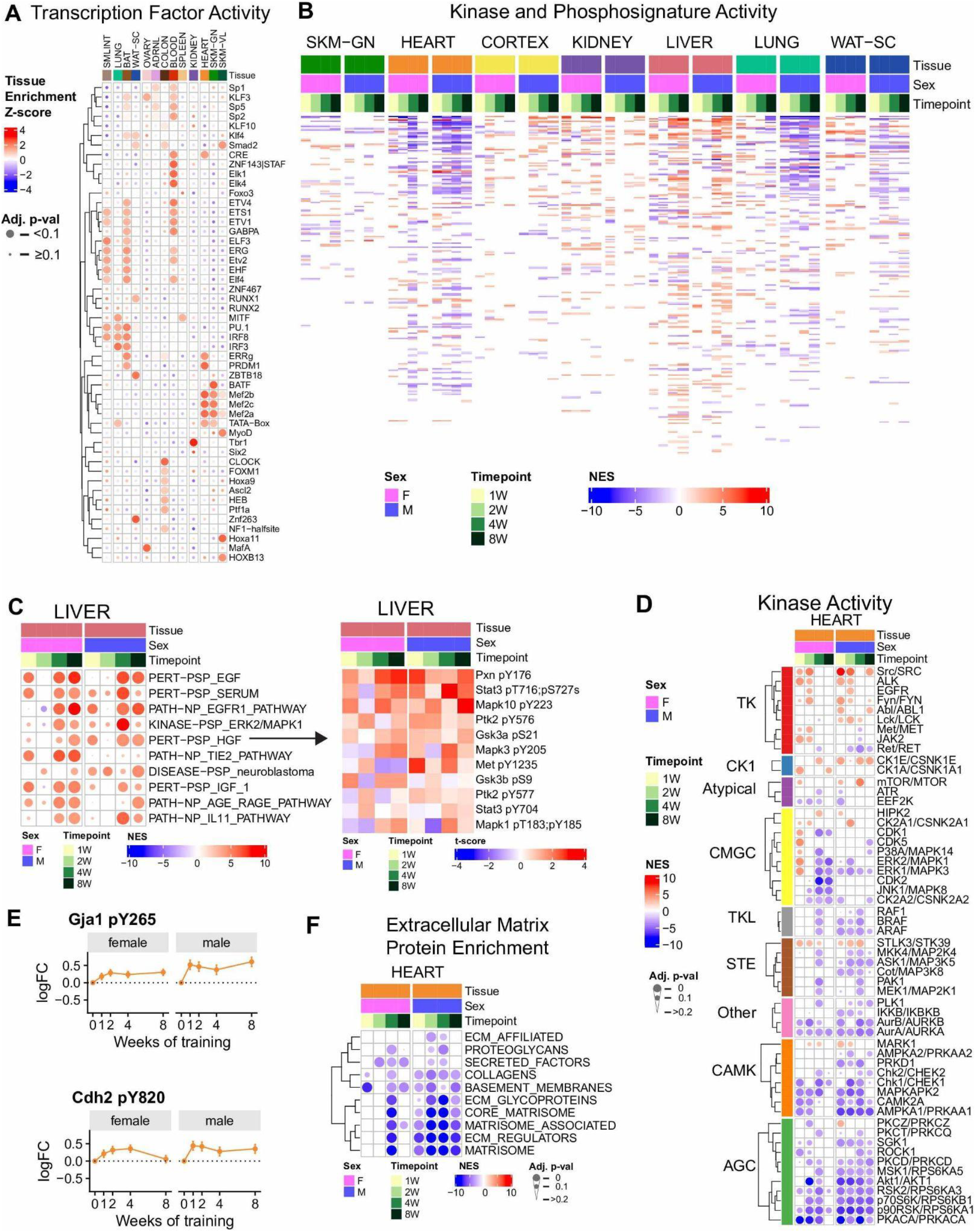
Regulatory signaling pathways modulated by endurance training. **A)** Transcription factor (TF) motif enrichment analysis of the training-differential transcripts in each tissue. Heatmap shows the motif enrichment z-score across the differential genes for the 13 tissues that had at least 300 genes after mapping transcript ids to gene symbols. For each tissue, the plot depicts the TFs that were both among the top ten significantly enriched and were expressed according to the RNA-seq data. Tissues were ordered based upon their clustering in Figure S4A, and TFs were hierarchically clustered by their enrichment across tissues. **B)** Estimate of activity changes in kinases and signaling pathways using PTM-SEA through enrichment of phosphoproteomics data. Heatmap shows normalized enrichment score (NES) for tissue/sex/time point groups as columns and kinases or pathways as rows. Non-significant enrichments are colored white (q-value > 0.1). Rows were sorted using the row sums of the absolute NES. **C)** Filtered PTM-SEA results for the liver showing kinases and signaling pathways with increased activity (left). Heatmap showing t-scores for phosphosites within the HGF signaling pathway (right). **D)** Filtered PTM-SEA results for the heart showing selected kinases with significant enrichments in at least one time point. Heatmap shows the NES as color and enrichment p-value as dot size. Kinases are grouped by kinase family and sorted by hierarchical clustering. **E)** Log_2_ fold-changes for selected Src kinase phosphosite targets, GJA1 pY265 and CDH2 pY820, in the heart. These phosphosites show a significant response to exercise training (5% FDR). **F)** Gene Set Enrichment Analysis (GSEA) results from the heart global proteome dataset using the matrisome gene set database. Heatmap shows NES as color and enrichment p-value as dot size. Rows are clustered using hierarchical clustering.

Next, we performed a comprehensive analysis of phosphosignature changes in response to training using PTM-SEA^38^ (Figure 3B; Table S6). Heart and liver tissues showed sex-consistent and robust decreased and increased phosphosignaling activity, respectively. In the liver, the top regulated phosphosignatures indicated increased growth factor response, including EGF, IGF, and the hepatic growth factor, HGF (q-value < 0.01; Figure 3C). The HGF phosphosignature could be explained by an increase in either HGF levels or HGF sensitivity in the liver. Exercise enhances liver health and metabolism^39,40^, yet the mechanism for improved liver health is incompletely understood. In rats, HGF treatment improves conditions of alcohol-induced fatty liver through reduced fibrosis^41–43^. STAT3 and PXN, HGF downstream targets involved in cell proliferation and migration, were differentially phosphorylated in response to training, suggesting a mechanism for activation of cellular maintenance in the liver (Figure S4B). Altogether we identify a potential mechanism of exercise-mediated improvement in liver health through HGF signaling.

In the heart, kinases showed bidirectional changes in their predicted basal activity in response to endurance training (Figure 3D). Some AGC protein kinases showed a concerted decrease in predicted activity, including AKT1 kinase, which is known to be involved in cardiac hypertrophy^44^. In contrast, some tyrosine kinases were predicted to have increased activity primarily at earlier time points, including SRC. SRC signaling has been implicated in the regulation of heart hypertrophy and structural remodeling^45–47^. Heart hypertrophy can occur as a beneficial physiological adaptation to exercise^48,49^. The known SRC target phosphosites GJA1 pY265 (q-value = 0.028) and CDH2 pY820 (q-value = 0.049) showed significantly increased phosphorylation in response to training (Figure 3E) with no significant change in the concomitant protein (q-value > 0.05; Figure S4C). Both *Gja1* (also known as *Cx43*) and *Cdh2* are known to regulate cell adhesion and communication and have been implicated in heart diseases^50,51^. Importantly, phosphorylation of GJA1 Y265 has previously been shown to disrupt gap junctions and regulate interactions with the tight junction protein ZO-1^52–54^. This suggests that SRC signaling may in part regulate reorganization of the heart extracellular structure. In agreement with this hypothesis, GSEA of extracellular matrix proteins revealed a negative enrichment in response to endurance training (Figure 3F; Table S7), showing decreased abundance of these proteins, such as basement membrane proteins (Figure S4D-E). Therefore, remodeling of the heart structure and ECM in response to endurance training could be explained in part through the activation of SRC via GJA1 phosphorylation.

### Temporal dynamics across tissues and omes identify molecular hubs for exercise adaptation

To compare the dynamic multi-omic responses to endurance training across tissues, we clustered the 34,244 differential features with complete timewise summary statistics using their timewise z-scores. We used the repfdr algorithm^55^ to assign features to one of nine possible combined states per time point, comprising one of three states per z-score (up, unchanged, down) for each sex (see Methods). The dynamics of the molecular training response can be visualized by constructing a summary graph in which rows represent these nine combined states and columns represent the four training time points. Nodes correspond to a combination of time, sex, and state. An edge connects two nodes from adjacent time points, representing a local temporal pattern. The differential abundance trajectory of any given training-regulated feature can then be represented by drawing a path through the nodes in this graph (Figure S5A; Table S8; Methods). When analyzing multiple analytes, this graphical analysis can be used to query the set of analytes that are associated with a specific node, edge, or a full path.

We summarized the node set sizes by tissue and time to identify the main temporal or sex-associated responses (Figure 4A). We observed that the small intestine, colon, and plasma had more changes at the early time points. Conversely, many up-regulated features in brown adipose tissue and down-regulated features in subcutaneous white adipose tissue showed a delayed response, observed only at week 8. The largest proportion of opposite effects between males and females were observed at week 1 in the adrenal gland. Other tissues, including the blood, heart, lung, kidney, and skeletal muscles (gastrocnemius and vastus lateralis), had relatively consistent numbers of up- and down-regulated features at all training time points.

**Figure 4.**
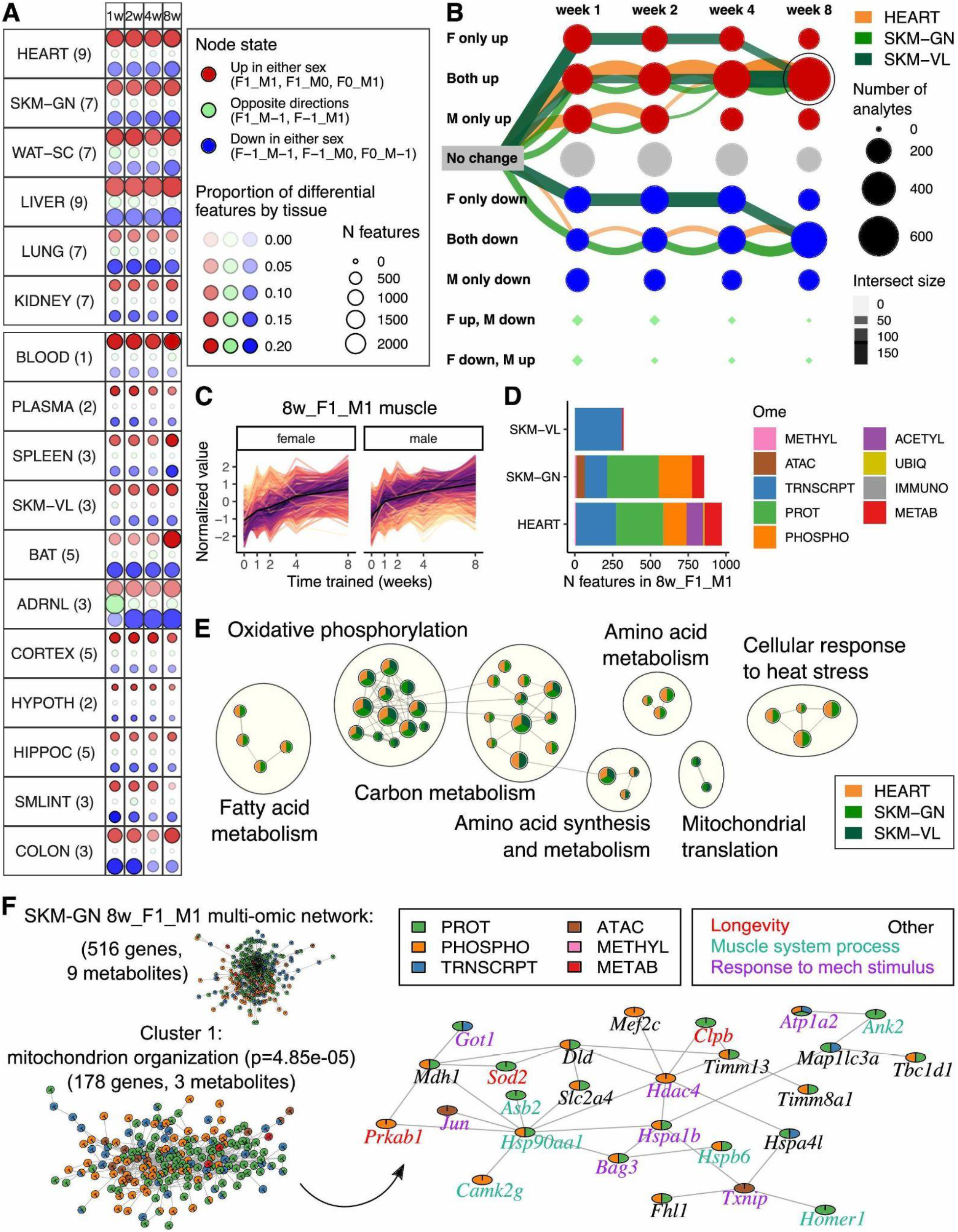
Temporal patterns of the molecular training response. **A)** Number of training-regulated features assigned to groups of graphical states across tissues and time. Red points include features that are up-regulated in at least one sex. That is, only in males (denoted as F0_M1), only in females (F1_M0), or in both sexes (F1_M1). Blue points include features that are down-regulated in at least one sex (states labeled F-1_M0, F0_M-1, and F-1_M-1, similarly to the notation for the up-regulated nodes); green points include features that are up-regulated in males and down-regulated in females or vice versa (states F-1_M1, F1_M-1). Point size is proportional to the number of differential features, scaled across all tissues. Point opacity is proportional to the fraction of total differential features represented by that point, calculated separately within each tissue. Features can be represented in multiple points. The number of omes profiled in each tissue is provided in parentheses next to the tissue abbreviation. The six most deeply profiled tissues are shown at the top and are separated from all other tissues. **B)** Graphical representation of the training-differential transcripts in the three muscle tissues: heart (HEART), gastrocnemius (SKM-GN), and vastus lateralis (SKM-VL). Each node represents one of nine possible states (row labels) at each of the four training time points (column labels). Edges are drawn through these nodes to represent the path of differential features over the training time course. This graph includes the three largest paths of differential transcripts in each of the muscle tissues, with edges colored by tissue. Both node and edge size are proportional to the number of features represented by the node or edge. The 8w_F1_M1 node, i.e., the group of features that are up-regulated in both females and males at 8 weeks, is circled. **C)** Line plots of standardized abundances of training-differential muscle features across all omes in the 8w_F1_M1 node. The black line in the center represents the average value across all features. **D)** Number of 8w_F1_M1 training-regulated muscle features shown in **(C)** corresponding to different omes and muscle tissues. **E)** Network view of pathway enrichment results corresponding to the features in **(C)**. Nodes indicate significantly enriched pathways (10% FDR); an edge represents a pair of nodes with a similarity score of at least 0.3 between the gene sets driving each pathway enrichment. Nodes (pathways) are only included if they are significantly enriched in at least two of the muscle tissues, as indicated by node color. Node size is proportional to the number of differential feature sets (e.g., gastrocnemius transcripts) for which the pathway is significantly enriched. Clusters of enriched pathways were defined using Louvain community detection, and are annotated high-level biological themes. **F)** Clustering analysis reveals a connected multi-omic network of stress response and muscle system processes. Top left: the input gene and metabolite protein-protein and gene-metabolite interactions network corresponding to features from the 8w_F1_M1 node of gastrocnemius (516 genes, 9 metabolites). Bottom left: the largest cluster identified using the leading eigenvector clustering algorithm. Right: a subnetwork of significant enrichments from the largest cluster contains genes functionally related to longevity, muscle system processes, and response to mechanical stimulus. Node colors indicate the ome through which the gene was identified.

We next focused on characterizing the shared molecular responses to endurance training in the three striated muscle tissues (heart, gastrocnemius, and vastus lateralis). First, to accommodate limited molecular profiling in the vastus lateralis, we visualized the three largest graphical trajectories, or paths, of training-regulated transcripts in each tissue (Figure 4B). For all three muscle tissues, both the up- and down-regulated paths concordantly converged into a sex-consistent response. Two of the three vastus lateralis paths represented transcripts that were either up-or down-regulated at all time points in females but were unchanged in males until week 8, at which time both sexes exhibited the same direction of effect. We next examined the full multi-omic set of analytes that were up-regulated in week 8 in these tissues (Figure 4C). In the gastrocnemius and heart, where global proteomics and phosphoproteomics data were available, a large proportion of the features corresponded to differential proteins and phosphosites (Figure 4D). A smaller proportion of the heart features corresponded to differential acetylsites. Pathway enrichment analysis of the genes associated with these differential features, which was performed separately for features in each tissue and ome (Table S9), demonstrated a shared endurance training response that reflected mitochondrial metabolism, biogenesis, and translation, and cellular response to heat stress (Table S9; Figure 4E), which is consistent with existing literature for skeletal^6,56^ and cardiac^57^ muscles.

We utilized a network analysis framework to further characterize the multi-omic, sex-consistent, up-regulated features in the gastrocnemius. After mapping these features to genes, we observed modest yet significant overlaps between the transcriptomic, chromatin accessibility, and proteomic assays but not with the methylation assays (Figure S5B). We then compared three different resources of biological interaction networks: BioGRID^58^, STRING^59^, and biological pathways, including interactions from Reactome, PathBank, and KEGG^60–62^. Network connectivity analysis showed that all three networks had significant enrichments of interactions between genes identified by a single ome and multi-omic genes (i.e., genes identified by two or more omes), as well as among these multi-omic genes (Figure S5C; Methods). Network eigenvector clustering^63^ of the BioGRID network, which outperformed the other networks in our tests above (Fig. S5C), identified three clusters of nodes (genes or metabolites, Table S10). The largest cluster had 181 nodes, was more balanced in its omic representation (Figure S5D), and was significantly enriched (at 5% FDR) for multiple muscle adaptation processes (Table S11), including: mitochondrion organization, longevity, muscle system processes, and response to mechanical stimulus. A subset of the cluster that focuses on these functions is shown in Figure 4F. This network contains many of the heat shock proteins identified in Fig. 2D-G, including HSP90AA1 and HSPA1B as major hubs, as well as the MEF2C transcription factor suggested to increase activity in our TF enrichment analysis (Fig. 3A). HDAC4, showing increased phosphorylation with training, is a central network hub. Training-induced phosphorylation of HDAC4 by calcium/calmodulin-dependent protein kinase (CaMK) leads to its nuclear export and subsequent regulation of target genes^64,65^, including GLUT4 for glucose uptake in muscle^66^. In the cytosol, non-histone substrates of HDAC4 include the master transcription cofactor for mitochondrial and oxidative metabolism, PPARGC1A, and several of the myosin heavy chain isoforms^67^, linking the central network hub to both mitochondrial and structural remodeling in skeletal muscle. Thus, this network analysis links the HSPs and MEF2C identified in the integrative multi-tissue and regulatory analyses to other major biological processes, revealing key multi-omic regulatory hubs.

### Training-regulated features in rats are associated with human exercise responses, disease signatures, and complex traits

To systematically evaluate the translational value of our rat-based data, we integrated our results with human exercise studies and disease ontology annotations. First, we compared the transcriptomics results from the vastus lateralis to a previously published meta-analysis of long-term training gene expression studies from similar human skeletal muscle tissue^10^. A significant correlation was observed between the meta-analysis-inferred fold-changes and our results for both sexes at 8 weeks post training (p < 1e-20, Figure S6A). Focusing on our training-differential, sex-consistent genes, we observed a significant and direction-consistent enrichment of our gene sets in a GSEA analysis in which genes were ranked by their human meta-analysis results (p < 1e-5, Figure S6B). Human gene expression data have been shown to exhibit excessive heterogeneity of fold-changes across studies due to observed and unobserved effect modifiers, including sex, age, training duration, and transcriptomics platform^10^. These effects substantially limit the power of the meta-analysis, suggesting a high false negative rate^10^. In our analysis we observed a significant overlap between the training-regulated rat genes and human genes with high fold-change heterogeneity, suggesting that the rat data may help in identifying exercise-responsive genes that were not detected by the human meta-analysis (p < 1e-4, Figure S6C).

We next performed disease ontology enrichment analysis using the DOSE R package^68^ (Table S12; Methods). Down-regulated genes from white adipose tissue, kidney, and liver were enriched with several disease terms suggesting a link between the exercise response and type 2 diabetes (T2D), heart disease, obesity, and kidney disease (5% FDR; Figure S6D), which are all epidemiologically related co-occurring diseases^69^. Leptin (*Lep*) and methallopeptitase 2 (*Mmp2*), which were down-regulated in white adipose tissue, are both associated with adipogenesis, T2D, and heart disease^70,71^. The proinflammatory agent Apolipoprotein C-III (*Apoc3*), down-regulated in white adipose tissue and the kidney, is associated with T2D, heart, and kidney diseases. *Apoc3* reduction in mice was recently demonstrated to protect white adipose tissues against obesity-induced inflammation and insulin resistance^72^. Overall, these results support a high concordance of our rat-based data with human studies and their relevance to human disease.

### Endurance training causes sex-specific responses in multiple tissues

Most tissues showed sex differences in their training responses. To further investigate these sex-biased effects, we performed two analyses across all tissues. First, we calculated the distributions of the correlations between the male and female timewise effects for all differential analytes, where a distribution with a dense non-positive correlation indicates uncorrelated or opposite regulation (Figure 5A, Figure S7). Prominent opposite responses were observed in training-regulated adrenal gland transcripts, lung phosphosites and chromatin accessibility peaks, white adipose tissue transcripts, and liver acetylsites. In addition, proinflammatory cytokines exhibited substantial sex-associated changes across tissues (Figure 5B, Figure S8A; Table S13). Most female-specific cytokines were differentially regulated between 1 and 2 weeks, while most male-specific cytokines were differentially regulated between 4 and 8 weeks (Figure 5C).

**Figure 5.**
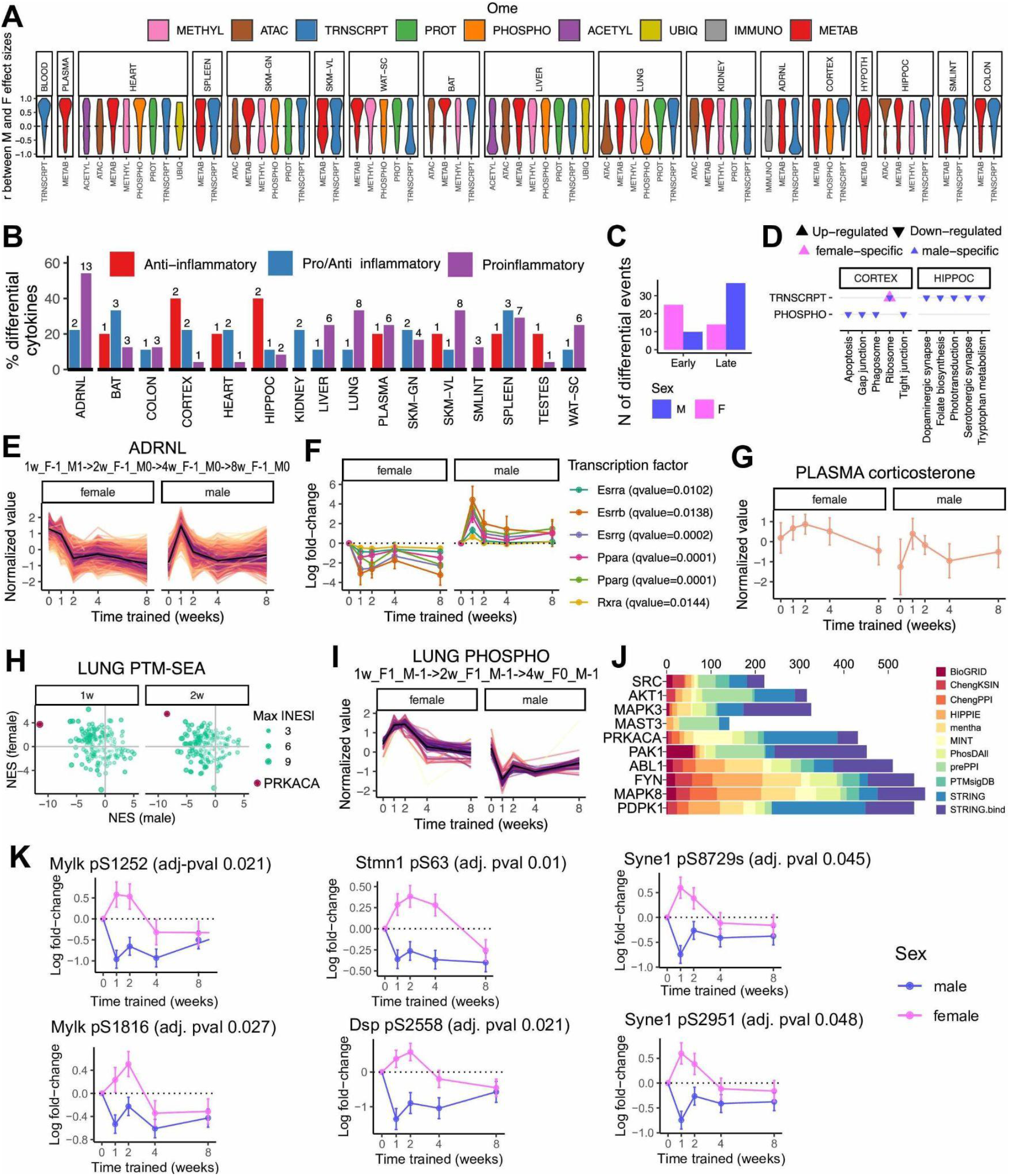
Sex differences in the endurance training response. **A)** Violin plots of Pearson correlations between male and female log_2_ fold-changes for each training-differential feature within each tissue and ome (5% FDR), colored by ome. **B)** Training-differential cytokines across tissues. 5, 24, and 9 cytokines were annotated as anti-, pro-, and pro/anti-inflammatory, respectively. Bars indicate the proportion and number (on top of each bar) of annotated cytokines in each category that are differential (5% FDR). **C)** Counts of early vs. (1-or 2-week) late (4-or 8-week) differential cytokines, according to states assigned by the graphical analysis, including all tissues. Cytokines with both early and late responses in the same tissue were excluded. **D)** Top five KEGG pathway enrichments (by sum-of-log combined p-value per tissue and pathway) in 8-week sex-stratified nodes (i.e., nodes representing training-differential features that do not have the same direction of effect in both sexes) in cortex and hippocampus. Each point represents a significant pathway enrichment in a given node, where the direction of the triangle points in the direction of the training effect (up or down) and the color indicates the corresponding sex. **E-F)** Sex difference in the transcriptional training response in the adrenal gland. **E)** Line plots of standardized abundances of training-differential features that follow the largest graphical path in the adrenal gland (i.e., 1w_F-1_M1->2w_F-1_M0->4w_F-1_M0->8w_F-1_M0 according to our graphical analysis notation). The black line in the center represents the average value across all features. **F)** Line plots of transcript-level log_2_ fold-changes corresponding to six transcription factors (TFs) whose motifs are significantly enriched by transcripts in **(E)**. TF motif enrichment q-values are provided in the legend. **G)** Line plot of normalized plasma corticosterone levels. **H-K)** Sex difference in the lung phosphosignature. **H)** Male versus female NES from PTM-SEA in the lung. Anticorrelated points corresponding to PRKACA NES are in dark red. **I)** Line plots of standardized abundances of training-differential phosphosites that follow the largest graphical edges of phosphosites in the lung (1w_F1_M-1->2w_F1_M-1->4w_F0_M-1). **J)** Top ten kinases with the greatest over-representation of substrates (proteins) corresponding to training-differential phosphosites in **(I)**. MeanRank scores by library are shown, as reported by KEA3. **(K)** Line plots showing relative abundance of PRKACA phosphosite substrates identified in lung as differential with disparate sex responses (Mylk pS1252, Mylk pS1816, Stmn1 pS63, Dsp pS2558, Syne1 pS8729, and Syne1 pS2951).

Next, we examined the the sex-specific or sex-stratified nodes identified in the graphical analysis results. Most (58%) of the 8-week training-regulated features demonstrated sex-associated responses, corresponding to multiple enriched pathways (Table S9). For example, several sex-specific changes were observed in the brain (Figure 5D), which may result in variable functional changes across the two sexes. In the brain, the cortex showed decreased phosphorylation of junction proteins, which may be related to changes in brain-blood-barrier functions^73^. The hippocampus showed decreased serotonin expression which, in agreement with previous studies^74^, may be associated with the antidepressant effects of exercise. These high-level analyses identified numerous sex-dependent differences in endurance training adaptations across many tissues, emphasizing the importance of studying exercise effects in both sexes in humans and model systems. We proceeded to analyze the sex-dependent responses in several tissues in detail.

#### The adrenal gland undergoes sex-specific adaptations through the regulation of steroid and hormone metabolism

We observed extensive transcriptional remodeling of the adrenal gland, with more than 4000 differentially regulated genes. Notably, the largest graphical path of training-regulated features were negatively correlated between males and females, with sustained downregulation in females and transient upregulation at 1 week in males (Figure 5E). Endurance training increases adrenal gland mass in both sexes, with more pronounced hypertrophy in male rats^75^. While we did not directly quantify changes in tissue mass, the 1-week induction in males could be explained by transient training-induced hypertrophy of the adrenal gland. The genes in this path were also associated with metabolism and steroid hormone synthesis pathways, particularly those pertaining to mitochondrial function (e.g., oxidative phosphorylation, lipid metabolism, and amino acid metabolism; Table S9). This result suggests that the adrenal gland additionally undergoes metabolic modifications related to its major function in production and release of different steroid hormones and catecholamines in response to endurance training. Further, TF motif enrichment analysis of the transcripts in this path showed enrichment of 14 TFs (5% FDR, Table S14), including the metabolism-regulating factors PPARg, PPARa, estrogen-related receptor gamma (ERRg), and the vitamin D receptor. The mRNA expression levels of several significantly enriched TFs themselves followed the same trajectory as this path (Figure 5F). PPARs are known regulators of lipid metabolism in response to endurance training^26^, and ERRg regulates mitochondrial metabolism in both cardiac^76^ and skeletal muscles^77^. Additionally, plasma corticosterone levels mirrored the male response of this adrenal path (Figure 5G). Altogether, the adrenal gland shows substantial sex differences in endurance training adaptation at the molecular level, likely leading to altered metabolic or hormonal functions relevant for the physiology of each of the two sexes.

#### Female-specific training adaptations through phosphorylation are associated with lung mechanical functions and disease

In healthy humans, anatomical differences between the sexes cause female-specific physiological lung responses to exercise^78^. In the rat lung, we observed decreased phosphosignaling activity with training in males, with opposite effects in females (Fig. 3B). Among these, PRKACA had the strongest sex difference in the PTM-SEA analysis at 1 and 2 weeks, with one of the greatest upregulation patterns in females (Figure 5H; Table S6). Consistently, the graphical analysis identified a training phosphosignature in the lung that was negatively correlated between the sexes and showed enrichment for PRKACA substrates (Figure 5I-J). PRKACA is a kinase involved in signaling within multiple cellular pathways. However, the four PRKACA substrates that followed this pattern were in proteins associated with cellular structures (e.g., cytoskeleton and cell-cell junctions): DSP, MYLK, STMN1, and SYNE1 (Figure 5K).

The phosphorylation of these proteins support a sex-dependent role of PRKACA in exercise-mediated regulation of lung structure and mechanical function. For example, both DSP and MYLK are associated with physiological modifications in the lung. DSP, a major component of desmosomes, gives mechanical strength to the alveolar epithelium in the lung^79,80^, and DSP phosphorylation has previously been associated with desmosome disassembly^81^. Genetic variants in *Dsp* are associated with various lung diseases^82–90^. MYLK plays an essential role in endothelial cell cytoskeleton rearrangement^91–93^, and *Mylk* is a candidate gene for inflammatory lung diseases^92^. Considering these associations and the known physiological sex differences in the lung, we hypothesize that increased phosphorylation of DSP and MYLK in the female lung during the early weeks of training plays a role in adaptation to increased mechanical stress.

#### Immune cells are recruited to male white and brown adipose tissues

Immune pathway enrichment analysis of the training-regulated transcripts at 8 weeks showed limited enrichment in the muscle (heart, gastrocnemius, and vastus lateralis) and brain (cortex, hippocampus, hypothalamus) tissues, downregulation in the male lung and female small intestine, and strong upregulation in both adipose tissues in males only (Figure 6A, Figure S8B; Table S9). While infiltration and activation of immune cells in adipose tissues are typically discussed in the context of obesity and insulin resistance^94–98^, immune cell induction also plays a critical role in tissue remodeling^99^. Since body fat percentage decreased after 8 weeks of endurance training in males only (Fig. S1B), we further investigated the male-specific up-regulated features at 8 weeks in brown and white adipose.

**Figure 6.**
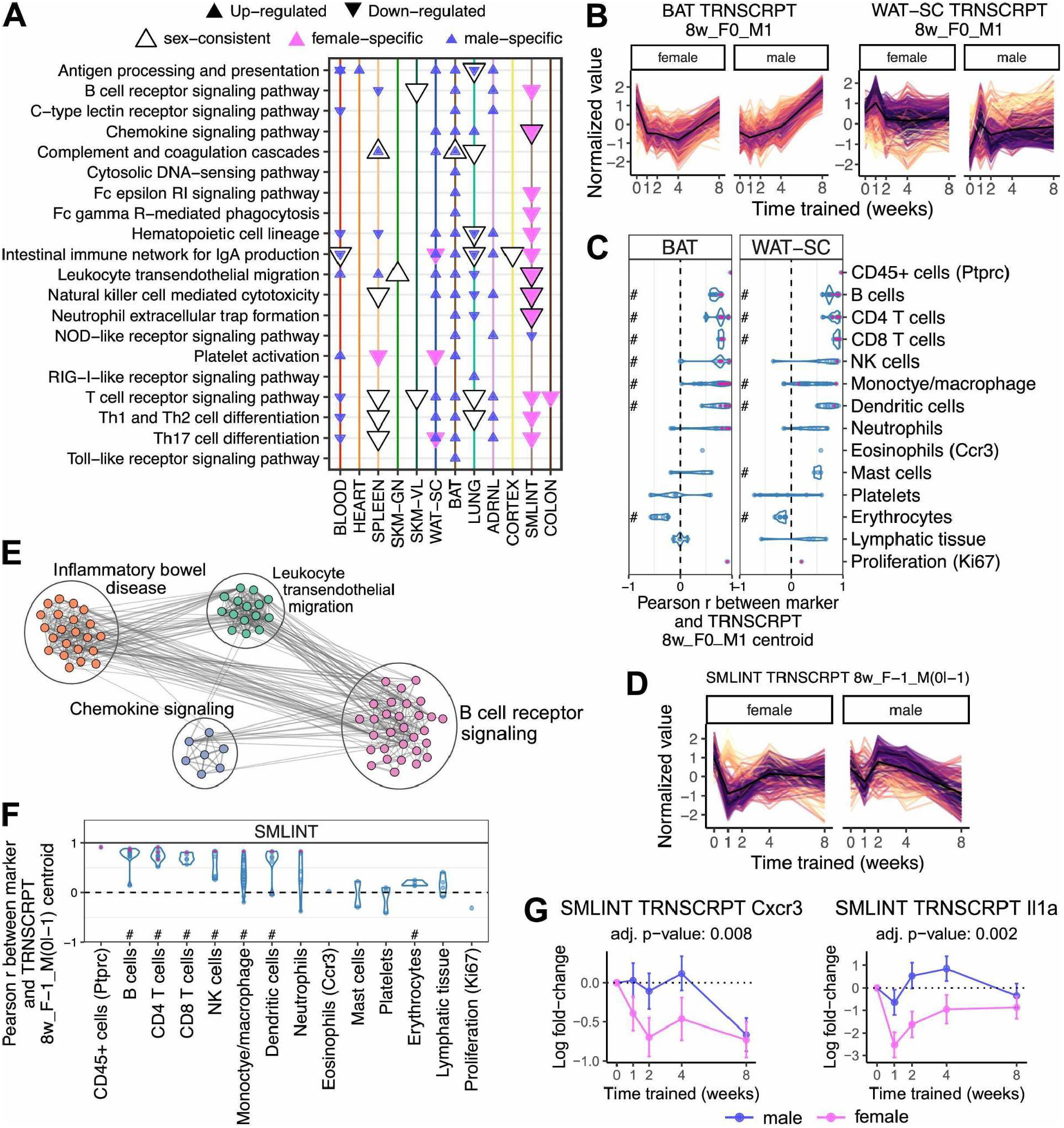
Training-induced immune responses. **A)** Enrichment analysis results of the training-differential transcripts at 8 weeks in KEGG immune system pathways (10% FDR). Each point represents a pathway enrichment in the set of transcripts contained in a particular 8-week node of the graphical clustering results, e.g., a blue triangle pointing up indicates an enrichment in the 8w_F0_M1 node. **B)** Line plots of standardized abundances of training-differential transcripts that are up-regulated only in males at the 8-week time point (8w_F0_M1) in brown adipose tissue (BAT) and WAT-SC. The black line in the center represents the average value across all features. **C)** Violin plots of the sample-level Pearson correlation between markers of immune cell types, lymphatic tissue, or cell proliferation and the average value of features in **(B)** at the transcript level. A red point indicates that the marker is also one of the differential features plotted in **(B)**. # indicates that the distribution of Pearson correlations for a set of at least two markers is significantly different from 0 (one-sample t-test, 5% BY FDR). When only one marker is used to define a category on the y-axis, the gene name is provided in parentheses. **D)** Line plots of standardized abundances of training-differential transcripts in the small intestine (SMLINT) that are down-regulated in females and either null or down-regulated in males at the 8-week time point (8w_F-1_M0 or 8w_F-1_M-1). **E)** Network view of pathway enrichment results corresponding to features in **(D)**. Nodes indicate significantly enriched pathways (10% FDR); edges connect nodes if there is a similarity score of at least 0.375 between the gene sets driving each pathway enrichment. Clusters of enriched pathways were defined using the Louvain algorithm for community detection and annotated with high level biological themes. **F)** Same as **(C)** but instead for features in **(D)**. **G)** Line plots showing the log_2_ fold-changes for *Cxcr3* and *Il1a* transcripts in the small intestine.

First, we observed that many of the same immune pathways were enriched in both the white and brown adipose tissues (Table S15). Second, this concordance was also reflected at the level of TF activity, where we observed similar enrichments of immune-related TFs in both types of male adipose tissue, including PU.1, IRF8, and IRF3 (q-value < 0.001, Table S16). Third, we used immune cell markers from external cell typing assays (Table S17; Methods) as proxies for immune cell type abundance and correlated their expression profiles with the profiles of our male-specific up-regulated transcripts and proteins (Figure 6B, Figure S8C). We observed a strong positive correlation for numerous immune cells, including B, T, and natural killer (NK) cells, and low correlation with platelets, erythrocytes, and lymphatic tissue at the transcript level (Figure 6C). These patterns suggest recruitment of peripheral immune cells or proliferation of tissue-resident immune cells as opposed to non-biological variation in blood or lymph content. The correlations at the protein level were not as striking (Figure S8D), likely because many of the markers were not robustly detected. In complementary analyses, cell type deconvolution and enrichment of immune cell type gene expression signatures in the bulk RNA-Seq data with CIBERSORTx^100^ recapitulated the upregulation of multiple immune cell types in males after 8 weeks of training (Figure S8E-F).

Endurance training has previously been shown to increase abundance of CD8+ T cells and NK cells in rodent adipose tissue^101^, and adipose tissue may serve as a reservoir for T cells^102^. NK cells are the most exercise-responsive immune cells^103^ and may play a role in limiting adipocyte hypertrophy^104^. Furthermore, previous studies have identified an association between immune cell recruitment and lipolysis^105–108^. While adipose immune responses are most often associated with pathogenic inflammation induced by obesity^109,110^, our data suggest an important role of increased immune cell activity in male adipose adaptation to endurance training.

### Intestine genes associated with inflammatory bowel disease are down-regulated in the response to endurance training

The small intestine was among the tissues with the highest enrichment in immune-related pathways (Fig. S8B). The main pattern of differential expression in our graphical analysis indicated downregulation of transcripts in females at week 8 (Figure 6D). This transcript set was significantly enriched with pathways related to inflammatory bowel disease (IBD), chemokine signaling, leukocyte transendothelial migration, and B cell receptor signaling (Figure 6E; Table S9). Using the same immune cell type marker correlation analysis described above, we observed positive associations between these transcripts and markers of several immune cell types, including B cells, T cells, NK cells, monocytes/macrophages, and dendritic cells, suggesting decreased abundance of these immune cell populations (Figure 6F). Specifically, our gene set included genes involved in B cell receptor signaling activation (*Btk, Cd72, Cd79a*) and B and T cell recruitment and activation (*Cd3e, Gata3, Il2rg, Ptpn22, Zap70, Cd5, Card11*). Endruance training also decreased expression of transcripts with genetic risk loci for IBD (*Pntp22*, MHC class II [*RT1-Dob, RT1-Db2*]), *Itgal*)^111^. The pathogenesis of IBD and other autoimmune disorders is associated with reduced or altered microbial diversity, inflammation, and reduced gut barrier integrity, which lead to systemic inflammation and reduced immune tolerance^112–115^. Endurance training is suggested to reduce systemic inflammation and improve health, in part by increasing gut microbial diversity and gut barrier integrity^116^. In accordance, we observed decreases in *Cxcr3* and *Il1a* with training (Figure 6G), both implicated in the pathogenesis of IBD^117–120^. CXCR3 increases small intestinal permeability^121^. Reduced expression of *Il1a* is indicative of improvements in gut tissue homeostasis^122^ and promotes localized immune cell recruitment^117,123^. Consistent with reduced *Il1a* expression, the expression of transcripts involved in superoxide production in the small intestine (*Rac2, Ncf1/Ncf4, Cybb*)^124,125^ were decreased with endurance training (Table S1). Reactive oxygen species increase intestinal permeability and propagate gut-mediated systemic inflammation^126,127^. Together, these down-regulated immune networks suggest that endurance training has the potential to reduce oxidative stress and inflammation in the small intestine, which may decrease intestinal permeability and thereby reduce systemic inflammation.

### Endurance training induces changes in mitochondria and lipid metabolism

To investigate organism-wide metabolic changes regulated by endurance training, we summarized the sex-consistent significant enrichments at 8 weeks using metabolic subcategories of the KEGG pathways (10% FDR, Figure 7A; Table S9). Down-regulated transcripts of brown adipose tissue were enriched with many metabolic pathways, suggesting an overall decrease in this tissue’s metabolic profile in agreement with a previous mouse study^128^. The heart showed enrichment of various carbohydrate metabolism subcategories across many omes, and remarkably, all enzymes within the glycolysis/gluconeogenesis pathway showed a consistent increase in abundance, except for GPI, FBP2, and DLAT (Figure S9A). The enrichment of oxidative phosphorylation was identified in the greatest number of tissues and is consistent with the joint analyses of the muscle tissues (Fig. 4E). We suspected that regulation of these mitochondrial pathways could be associated with changes in mitochondrial biogenesis due to endurance training. Therefore, we estimated proportional mitochondrial changes to endurance training using mitochondria RNA-seq reads (Figure 7B-C, Figure S9B) and changes of mitochondrial functions through GSEA using gene expression, protein abundance, and protein PTMs (Figure 7D, Figure S9C; Tables S18-S21; Methods). Increased mitochondrial biogenesis was consistently observed in the skeletal muscles, heart, and liver across these analyses. Moreover, sex-specific mitochondrial changes were observed in the adrenal gland, as described above, as well as in the colon, lung, and kidney. These results highlight the role of mitochondria in exercise adaptation across many tissues captured in this study.

**Figure 7.**
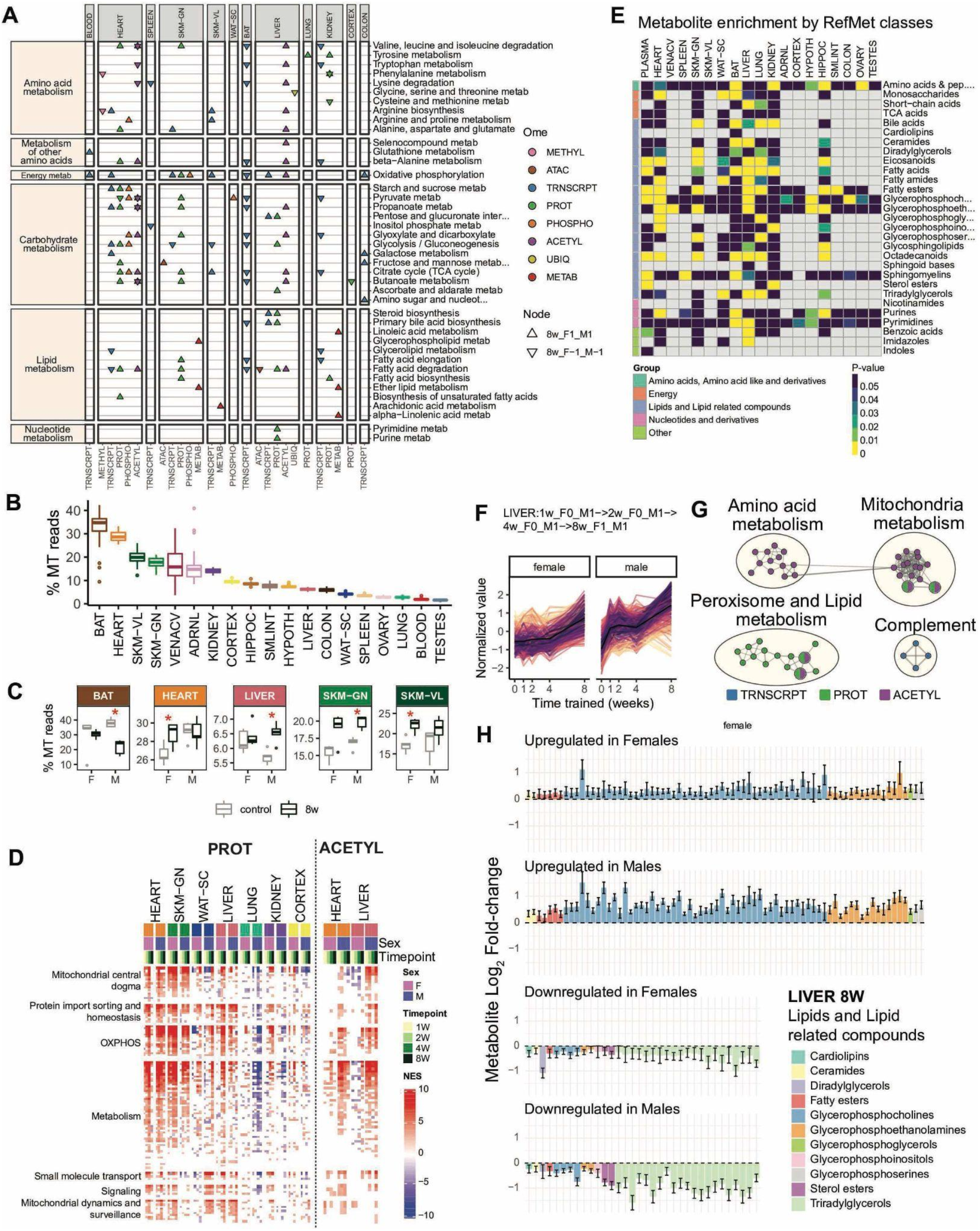
Training-induced changes in metabolism. **A)** Significant enrichments for relevant categories of KEGG metabolism pathways from features that are up-or down-regulated in both sexes at 8 weeks (8w_F1_M1 and 8w_F-1_M-1 nodes, respectively). Triangles point in the direction of the response (up or down). Points are colored by ome. **B)** Boxplots showing the percent of reads across samples in each tissue that map to the mitochondrial genome (% MT reads). **C)** Comparison of % MT reads between untrained controls and animals trained for 8 weeks. Plot shows tissues with a statistically significant change after 8 weeks in at least one sex (red asterisk, Dunnett’s test, 10% FDR). **D)** GSEA results using the MitoCarta MitoPathways gene set database and proteomics (PROT) or acetylome (ACETYL) timewise summary statistics for training. NES are shown for significant pathways (10% FDR). Mitochondrial pathways shown as rows are grouped using the parental group in the MitoPathways hierarchy. **E)** RefMet metabolite class enrichment calculated using GSEA with the -log_10_ training p-value. Colored cells are significant chemical class enrichments (5% FDR). **F)** Line plots of standardized abundances of liver training-differential features across all omes that are up-regulated in both sexes, with a later response in females (LIVER: 1w_F0_M1−> 2w_F0_M1−> 4w_F0_M1−> 8w_F1_M1). The black line in the center represents the average value across all features. **G)** Network view of pathway enrichment results corresponding to features in **(F)**. Nodes indicate significantly enriched pathways (10% FDR); edges connect nodes if there is a similarity score of at least 0.375 between the gene sets driving each pathway enrichment. Node colors indicate omes in which the enrichment was observed. Clusters of enriched pathways were defined using the Louvain algorithm for community detection and annotated with the high level biological themes. **H)** 8-week log_2_ fold-changes (relative to sedentary controls) for metabolites within the “Lipids and Lipid related compounds” category in the liver.

We summarized training-regulated metabolites across tissues by performing enrichment of RefMet metabolite classes (Figure 7E; Table S22). This analysis focused primarily on lipid classes, as 2105 out of the total 2430 detected metabolite features in our data were classified as lipids (Table S23). The liver showed the greatest number of significantly enriched classes, followed by the heart, lung, and hippocampus. The eicosanoid and octadecanoid lipid classes had significant enrichments in 8 and 7 tissues, respectively. Eicosanoids and octadecanoids are derived from 20- and 18-carbon polyunsaturated fatty acids, respectively, most frequently arachidonic acid (AA)^129^. The AA pathway plays a key role as an inflammatory mediator involved in many molecular and cellular functions under different physiological conditions^130^. Fatty acids and fatty esters, which serve as both energy sources and building blocks for cellular membranes, were enriched in 7 tissues. Within fatty esters, acylcarnitines showed unique regulation across the plasma, brown and white adipose tissues, heart, and liver (Figure S9D). An early increase in medium- and long-chain acylcarnitines occurred in the plasma, white adipose tissue, and liver, while a decrease in short-chain acylcarnitines occurred in brown adipose tissue and the heart. Together, these analyses reveal the dynamic mobilization of lipids across many tissues in response to endurance training.

In the liver, we observed substantial regulation of metabolic pathways across the proteome, acetylome, and lipidome (Fig. 7A, Fig. 7D, Fig. 7E). To investigate the temporal dynamics of metabolic regulation in the liver, we focused on the large group of features that increased in abundance over time for both sexes, with a larger effect in males (Figure 7F). Most of the liver features in this graphical path corresponded to protein abundance and protein acetylation changes in the mitochondrial, amino acid, and lipid metabolic pathways (Figure 7G; Table S24). Although changes in liver mitochondrial functions have been previously observed during exercise^131^, to our knowledge extensive changes in protein acetylation in response to endurance training have not been described. In addition to mitochondria, we observed increased abundance and acetylation of proteins from the peroxisome, an organelle with key functions in fatty acid beta-oxidation, sterol precursor synthesis, and etherphospholipid production (Figure S9E). Lipid metabolism is critical for liver health, and lipid dysregulation can lead to non-alcoholic fatty liver disease (NAFLD) and eventually to the advanced form of steatohepatitis (NASH). Exercise is one of the standard clinical interventions recommended against NAFLD^132^, though the molecular mechanisms are unclear. In our data, the liver showed significant enrichment in 12 metabolite classes belonging to “lipids and lipid-related compounds’’ (Fig. 7E), and we observed an increase in phosphatidylcholines (PCs) and a concomitant decrease in triacylglycerols (TAGs) (Figure 7H). Increased PC and decreased TAG are associated with healthy liver metabolism, and the opposite is associated with NAFLD^133^. Moreover, although unrelated to exercise, it has been proposed that mitochondrial dysfunction is a key component of NAFLD progression^134^. Therefore, our study indicates that endurance training induces profound molecular changes in mitochondrial protein abundance and acetylation, which may be associated with improved liver health through the regulation of lipids.

## Discussion

Mapping the molecular responses to endurance exercise training in a whole organism is critical for gaining a holistic understanding of the mechanisms that underlie the benefits of exercise. Previous studies have provided valuable findings but are restricted in scope because they used limited omics platforms, examined few tissues, interrogated a narrow temporal range, or were biased towards a single sex. As large-scale publicly available omic data have proven to be indispensable for rapidly advancing biomedical research, e.g, ^135–138^, we expect that our new resource will similarly accelerate the generation of novel hypotheses pertaining to the molecular basis of endurance training. This work illustrates how mining our data resource can both recapitulate expected mechanisms and provide novel biological insights. Together, our study provides a comprehensive multi-omic, multi-tissue map of the temporal endurance training responses in both male and female rats.

We identified thousands of training-associated molecular alterations within and across tissues. We observed genes, proteins, and other molecular analytes that responded to endurance training with both shared and specific cellular responses and early and late dynamics. While the fold-changes of many training-regulated features associated with tissue remodeling during endurance exercise training are modest, they represent coherent processes. We detected wide-spread effects that likely transduce remodeling of multiple tissues simultaneously, including alterations of mitochondrial functions, induction of a global heat shock response, activation of tissue-specific response programs, and regulation of immune-related processes.

The translational aspects of our findings illustrate how our rich dataset can be leveraged to deepen our understanding of exercise-related improvement of health and prevention and/or reversal of diseases. The global heat shock response to exercise may confer cytoprotective effects, including in pathologies related to tissue damage and injury recovery^34,139–141^. The acetylation of liver mitochondria and regulation of lipid metabolism provide a potential mechanism for protection against non-alcoholic fatty liver disease and steatohepatitis^132^. Training-modulated cytokines and receptors are linked to intestinal inflammation, including transcripts with genetic risk loci for inflammatory bowel disease^111^. Finally, the regulation of kinase activity in the lung may be associated with lung mechanical stress^142,143^.

The sex-biased responses to exercise training have not been well characterized. We observed sex differences in the training response across the majority of tissues and omes, highlighting the critical importance of including both sexes in exercise science research. Though we recognize that that sexual dimorphism is substantially greater in rats than in humans^144^, this work identified important sex-specific adaptive processes in the white adipose tissue, brain, adrenal gland, and lung, with implications for how endurance training may differentially improve health in males and females.

We note limitations in our experimental design, datasets, and analyses. Our assays were performed on bulk tissue and do not cover single-cell platforms. Our resource has limited omic characterization for certain tissues, and some omes with emerging biological relevance were not performed in this study, for example, microbiome profiling. We analyzed 3-6 animals per time point and sex combination, and this sample size may limit our ability to identify modest yet physiologically relevant molecular alterations. The findings and associations we describe are hypothesis-generating and require biological validation. Towards this end, we have generated a tissue bank from this study to facilitate detailed hypothesis-driven work by others.

This landscape resource provides future opportunities to enhance and refine the molecular map of the endurance training response. We expect that this dataset will remain an ongoing platform to translate tissue- and sex-specific molecular changes in rats into humans. MoTrPAC has placed extensive effort into facilitating access, exploration, and interpretation of this resource, as the program aims to provide key data resources to enable integration with public resources. Combined, this first MoTrPAC multi-omic resource sets the landscape for rapid advancement of our understanding of the milieu of molecular changes in endurance training adaptation and provides transformative opportunities to understand the impact of exercise on health and disease.

## Primary authors

### Lead Analysts†

David Amar**‡**, Nicole R. Gay**‡**, Pierre M. Jean-Beltran**‡**

### Lead Data Generators†

Dam Bae, Surendra Dasari, Courtney Dennis, Charles R. Evans, David A. Gaul, Olga Ilkayeva, Anna Ivanova, Maureen T. Kachman, Hasmik Keshishian, Ian R. Lanza, Ana C. Lira, Michael J. Muehlbauer, Venugopalan D. Nair, Paul D. Piehowski, Jessica S. Rooney, Kevin S. Smith, Cynthia L. Stowe, Bingqing Zhao

### Analysts†

David Jimenez-Morales*, Malene E. Lindholm*, James A. Sanford*, Gregory R. Smith*, Nikolai G. Vetr*, Tiantian Zhang*, Bingqing Zhao*, Jose J. Almagro Armenteros, Julian Avila-Pacheco, Nasim Bararpour, Yongchao Ge, Zhenxin Hou, Anna Ivanova, Gina M. Many, Shruti Marwaha, David M. Presby, Archana Natarajan Raja, Evan Savage, Alec Steep, Yifei Sun, Si Wu, Jimmy Zhen

### Animal Study Leadership†

Sue C. Bodine^, Karyn A. Esser^, Laurie J. Goodyear, Simon Schenk^

### Manuscript Writing Group Leads

Nicole R. Gay**‡**, David Amar**‡**, Pierre M. Jean-Beltran**‡**

### Manuscript Writing Group

Malene E. Lindholm, Gina M. Many, Simon Schenk^, Stephen B. Montgomery^, Jose J. Almagro Armenteros*, Julian Avila-Pacheco*, Nasim Bararpour*, Sue C. Bodine*^, Karyn A. Esser*^, Facundo M. Fernández*, Zhenxin Hou*, David Jimenez-Morales*, Archana Natarajan Raja*, James A. Sanford*, Stuart C. Sealfon, Gregory R. Smith*, Michael P. Snyder*^, Nikolai G. Vetr*, Bingqing Zhao*, Tiantian Zhang*

### Senior Leadership†

Joshua N. Adkins, Euan Ashley, Sue C. Bodine^, Charles Burant, Steven A. Carr^, Clary Clish, Gary Cutter, Karyn A. Esser^, Facundo M. Fernández, Robert Gerszten, Laurie J. Goodyear, William E. Kraus, Ian R. Lanza, Jun Li, Michael E. Miller, Stephen B. Montgomery^, K. Sreekumaran Nair, Christopher Newgard, Eric A. Ortlund, Wei-Jun Qian, Simon Schenk^, Stuart C. Sealfon, Michael P. Snyder^, Russell Tracy, Martin J. Walsh, Matthew T. Wheeler^

**†**Alphabetic order

**‡**Co-first author

*Contributed equally

^Co-corresponding author

## MoTrPAC Study Group

### Consortium Coordination**†**

**Bioinformatics Center - Stanford University, Stanford, CA**

David Amar**‡**, Euan Ashley, Karen P. Dalton, Trevor Hastie, Steven G. Hershman, David Jimenez-Morales, Malene E. Lindholm, Shruti Marwaha, Archana Natarajan Raja, Mihir Samdarshi, Christopher Teng, Rob Tibshirani, Matthew T. Wheeler^, Jimmy Zhen

**Biospecimens Repository - University of Vermont, Burlington, VT**

Elaine Cornell, Nicole Gagne, Sandy May, Jessica L. Rooney, Russell Tracy

**Administrative Coordinating Center - University of Florida, Gainesville, FL**

Brian Bouverat, Christiaan Leeuwenburgh, Ching-ju Lu, Marco Pahor

**Data Management, Analysis, and Quality Control Center - Wake Forest University School of Medicine, Winston-Salem, NC**

Fang-Chi Hsu, Michael E. Miller, Scott Rushing, Cynthia L. Stowe, Michael P. Walkup

**Exercise Intervention Core - Wake Forest University, Winston-Salem, NC**

Barbara Nicklas, W. Jack Rejeski

**National Institutes of Health, Bethesda, MD**

John P. Williams, Ashley Xia

### Preclinical Animal Study Sites**†**

**Joslin Diabetes Center, Boston, MA**

Brent G. Albertson, Tiziana Caputo, Laurie J. Goodyear, Michael F. Hirshman, Sarah J. Lessard, Nathan S. Makarewicz, Pasquale Nigro, David M. Presby, Krithika Ramachandran

**University of California, Los Angeles, CA**

Andrea Hevener

**University of Florida, Gainesville, FL**

Elisabeth R. Barton, Karyn Esser^, Roger Farrar, Scott Powers

**University of Iowa, Iowa City, IA**

Dam Bae, Sue C. Bodine^, Michael Cicha, Luis Gustavo Oliveira De Sousa, Bailey E. Jackson, Kyle S. Kramer, Ana K. Lira, Andrea Marshall, Collyn Z-T. Richards, Simon Schenk^

**University of Kansas Medical Center, Kansas City, KS**

John Thyfault

**University of Missouri, Columbia, MO**

Frank W. Booth, R. Scott Rector

**University of Virginia School of Medicine, Charlottesville, VA**

Benjamin G. Ke, Zhen Yan, Chongzhi Zang

### Chemical Analysis Sites**†**

**Broad Institute, Inc., Boston, MA**

Julian Avila-Pacheco, Steven Carr^, Clary B. Clish, Courtney Dennis, Robert E. Gerszten, Pierre M. Jean Beltran**‡**, Hasmik Keshishian, DR Mani, Charles C. Mundorff, Cadence Pearce, Jeremy M. Robbins

**Duke University, Durham, NC**

Olga Ilkayeva, Michael Muehlbauer, Christopher Newgard

**Emory University, Atlanta, GA**

Zhenxin Hou, Anna A. Ivanova, Xueyun Liu, Kristal M. Maner-Smith, Eric A. Ortlund, Karan Uppal, Tiantian Zhang

**Georgia Institute of Technology, Atlanta GA**

Facundo M. Fernández, David A. Gaul, Samuel G. Moore, Evan M. Savage

**Icahn School of Medicine at Mount Sinai, New York City, NY**

Mary Anne S. Amper, Ali Tugrul Balci, Maria Chikina, Yongchao Ge, Kristy Guevara, Nada Marjanovic, Venugopalan D. Nair, German Nudelman, Hanna Pincas, Irene Ramos, Stas Rirak, Aliza B. Rubenstein, Frederique Ruf-Zamojski, Stuart C. Sealfon, Nitish Seenarine, Gregory R. Smith, Yifei Sun, Sindhu Vangeti, Mital Vasoya, Alexandria Vornholt, Martin J. Walsh, Xuechen Yu, Elena Zaslavsky

**Pacific Northwest National Laboratory, Richland, WA**

Joshua N. Adkins, Marina A. Gritsenko, Joshua R. Hansen, Chelsea Hutchinson-Bunch, Gina M. Many, Matthew E. Monroe, Ronald J. Moore, Michael D. Nestor, Vladislav A. Petyuk, Paul D. Piehowski, Wei-Jun Qian, Tyler J. Sagendorf, James A. Sanford

**Stanford University, Stanford, CA**

Jose Juan Almagro Armenteros, Nasim Bararpour, Clarisa Chavez, Roxanne Chiu, Nicole R. Gay**‡**, Krista M. Hennig, Chia-Jui Hung, Christopher A. Jin, Stephen B. Montgomery^, Daniel Nachun, Kevin S. Smith, Michael P. Snyder^, Nikolai G. Vetr, Si Wu, Navid Zebarjadi, Bingqing Zhao

**The Mayo Clinic, Rochester, MN**

Surendra Dasari, Ian Lanza, K. Sreekumaran Nair

**University of Michigan, Ann Arbor, MI**

Charles F. Burant, Charles R. Evans, Maureen T. Kachman, Jun Z. Li, Alexander (Sasha) Raskind, Tanu Soni, Alec Steep

### Clinical Sites**†**

**AdventHealth Translational Research Institute, Orlando, FL**

Paul M. Coen, Bret H. Goodpaster, Lauren M. Sparks

**Ball State University, Muncie, IN**

Toby L. Chambers, Bridget Lester, Scott Trappe, Todd A. Trappe

**Duke University, Durham, NC**

Kim M. Huffman, William E. Kraus, Megan E. Ramaker

**Pennington Biomedical Research Center, Baton Rouge, LA**

Kishore Gadde, Melissa Harris, Neil M. Johannsen, Robert L. Newton Jr., Tuomo Rankinen, Eric Ravussin

**The University of Alabama at Birmingham, Birmingham, AL**

Marcas Bamman, Thomas W. Buford, Gary Cutter, Anna Thalacker-Mercer

**University of California, Irvine, CA**

Dan Cooper, Fadia Haddad, Shlomit Radom-Aizik

**University of Colorado Anschutz Medical Campus, Aurora, CO**

Bryan C. Bergman, Daniel H. Bessesen, Catherine M. Jankowski, Wendy M. Kohrt, Edward L. Melanson, Kerrie L. Moreau, Irene E. Schauer, Robert S. Schwartz

**University of Texas Health Science Center, San Antonio, TX**

Nicolas Musi

**University of Texas Medical Branch, Galveston, TX**

Blake B. Rasmussen, Elena Volpi

**†**Alphabetic order

**‡**Co-first author

^Co-corresponding author

## Methods

See Additional File 1 for experimental and computational methods.

## Supporting information

Additional File 2 - Author Contributions

## Data availability

MoTrPAC data for this study are publicly available via https://motrpac-data.org/data-access. Data access inquiries should be emailed to. Additional resources can be found at https://motrpac.org and https://motrpac-data.org. Processed data and analysis results are additionally available in the MotrpacRatTraining6moData R package (https://motrpac.github.io/MotrpacRatTraining6moData).

## Code availability

Code for reproducing the main analyses are conveniently provided in the MotrpacRatTraining6mo R package (<u>github.com/MoTrPAC/MotrpacRatTraining6mo</u>). MoTrPAC data processing pipelines for RNA-Seq, ATAC-seq, RRBS, and proteomics are publicly available: https://github.com/MoTrPAC/motrpac-rna-seq-pipeline, https://github.com/MoTrPAC/motrpac-atac-seq-pipeline, https://github.com/MoTrPAC/motrpac-rrbs-pipeline, https://github.com/MoTrPAC/motrpac-proteomics-pipeline. Normalization and QC scripts are available at https://github.com/MoTrPAC/MotrpacRatTraining6moQCRep.

## Funding

The MoTrPAC Study is supported by NIH grants U24OD026629 (Bioinformatics Center), U24DK112349, U24DK112342, U24DK112340, U24DK112341, U24DK112326, U24DK112331, U24DK112348 (Chemical Analysis Sites), U01AR071133, U01AR071130, U01AR071124, U01AR071128, U01AR071150, U01AR071160, U01AR071158 (Clinical Centers), U24AR071113 (Consortium Coordinating Center), U01AG055133, U01AG055137 and U01AG055135 (PASS/Animal Sites). This work was also supported by other funding sources: NHGRI Institutional Training Grant in Genome Science 5T32HG000044 (N.R.G.), National Science Foundation Graduate Research Fellowship Grant No. NSF 1445197 (N.R.G.), National Heart, Lung, and Blood Institute of the National Institute of Health F32 postdoctoral fellowship award F32HL154711 (P.M.J.B.), the Knut and Alice Wallenberg Foundation (M.E.L.), National Science Foundation Major Research Instrumentation (MRI) CHE-1726528 (F.M.F.), National Institute on Aging P30AG044271 and P30AG003319 (N.M.), and NORC at the University of Chicago Grant # P30DK07247 (E.R.).

Parts of this work were performed in the Environmental Molecular Science Laboratory, a U.S. Department of Energy national scientific user facility at Pacific Northwest National Laboratory in Richland, Washington.

The views expressed are those of the authors and do not necessarily reflect those of the NIH or the Department of Health and Human Services of the United States.

## Author contributions

Detailed author contributions are provided in Additional File 2.

## Additional information

Supplementary Information is available for this paper. Correspondence and requests for materials should be addressed to.

Additional file 1: Extended methods.

Additional file 2: Author contributions.

Additional file 3: Supplementary figures.

Additional file 4: Supplementary tables.

## Abbreviations

ACETYL: Acetylproteomics; protein site acetylation
acetylsites: Lysine acetylation
ADRNL: Adrenal gland
ATAC: Chromatin accessibility, ATAC-seq data
BAT: Brown adipose tissue
BLOOD: Whole blood; Blood RNA
COLON: Colon
CORTEX: Cerebral cortex
CV: Coefficient of variation
ECM: Extracellular matrix
FDR: False discovery rate
GSEA: Gene set enrichment analysis
HEART: Heart
HIPPOC: Hippocampus
HSP: Heat shock protein
HYPOTH: Hypothalamus
IBD: Inflammatory bowel disease
IHW: Independent hypothesis weighting
IMMUNO: Multiplexed immunoassays
KEGG: Kyoto Encyclopedia of Genes and Genomes
KIDNEY: Kidney
LC-MS/MS: Liquid chromatography-tandem mass spectrometry
LIVER: Liver
LUNG: Lung
METAB: Metabolomics and lipidomics
METHYL: DNA methylation, RRBS data
MHC: Major histocompatibility complex
NAFLD: Nonalcoholic fatty liver disease
NASH: Nonalcoholic steatohepatitis
NES: Normalized enrichment score
NK: Natural killer
OVARY: Ovaries
PC: Phosphatidylcholine
PCA: Principal component analysis
PE: Phosphatidylethanolamine
PHOSPHO: Phosphoproteomics; protein site phosphorylation
PLASMA: Plasma (from blood)
PROT: Global proteomics; protein abundance
PTM: Post-translational modification
PTM-SEA: PTM signature enrichment analysis
RIN: RNA integrity number
SKM-GN: Gastrocnemius (skeletal muscle)
SKM-VL: Vastus lateralis (skeletal muscle)
SMLINT: Small intestine
SPLEEN: Spleen
TAG: Triacylglycerol
TESTES: Testes
TF: Transcription factor
TRNSCRPT: Transcriptomics, RNA-Seq data
UBIQ: Ubiquitylome; protein site ubiquitination
ubiquitylsites: Lysine diglycine ubiquitin remnants
VENACV: Vena cava
WAT-SC: Subcutaneous white adipose tissue

