## Additional File 1 - Methods for "Temporal dynamics of the multi-omic response to endurance exercise training across tissues"


EXPERIMENTAL DESIGN AND SAMPLE HANDLING 4
Animal handling 4
Animals 4
Treadmill familiarization and training 4
Body composition 6
Maximum oxygen consumption (VO2max) 7
Tissue collection 7
Sample distribution 7
Archiving 7
Selection of animals for molecular assays 8
Cryopulverization 8
Aliquoting 8
Reference standards 8
DATA PRODUCTION AND QUANTIFICATION 9
DNA methylation by reduced representation bisulfite sequencing 9
DNA extraction and library preparation 9
DNA sequencing, data processing, and normalization 10
Assay for transposase-accessible chromatin using sequencing 11
Nuclei extraction and library preparation 11
DNA sequencing, data processing, and normalization 11
RNA sequencing 12
Extraction of total RNA 12
mRNA Sequence Library Preparation 12
RNA sequencing, quantification, and normalization 13
Liquid chromatography tandem mass spectrometry (LC-MS/MS) proteomics 13
Tissue lysis and protein extraction 13
Protein digestion 14
Tandem mass tag (TMT) labeling of peptides 14
Phosphotyrosine peptide enrichment 15
Offline bRP fractionation 15
Phosphopeptide enrichment 16
Acetylpeptide enrichment 16
K-ε-GG enrichment for ubiquitylome analysis 16
LC-MS/MS analysis of heart and liver tissue samples 17

LS-MS/MS analysis of the gastrocnemius, white adipose tissue, cortex, kidney, and lung 18
Raw MS/MS data processing 19
Proteomics data normalization 20
Non-targeted metabolomics 21
HILIC LC-MS positive ion mode non-targeted metabolomics 21
Reverse phase and ion pairing LC-MS non-targeted metabolomics 22
Materials and reagents 22
Sample preparation 22
LC-MS analysis 24
Data analysis 25
Non-targeted LC-MS/MS lipidomics 26
Sample preparation 26
Data collection 26
Data processing 27
Quality control procedures 27
Targeted metabolomics 28
LC-MS/MS analysis of branched-chain keto acids 28
Flow injection MS/MS analysis of acyl CoAs 28
LC-MS/MS analysis of nucleotides 29
LC-MS/MS analysis of amino acids and amino metabolites 29
GC-MS analysis of tricarboxylic acid cycle (TCA) metabolites 30
LC-MS/MS analysis of ceramides 30
LC-MS/MS analysis of acylcarnitines 30
Targeted lipidomics of low-level lipids 31
Sample preparation 31
Data collection 32
Data processing and quality control 32
Metabolomics data processing and normalization 32
Multiplexed bead-based immunoassays 34
Protein extraction 34
Targeted multiplexed bead-based immunoassays 34
Data preprocessing 35
STATISTICAL ANALYSES 35
Outlier identification 35
Differential analysis 36
Meta-analysis of metabolomics results 37

Graphical clustering of differential analysis results 38
Comprehensive feature-to-gene map 40
Pathway enrichment analysis of graphical clusters 40
Biological networks 41
Network connectivity analysis 42
Transcription factor enrichment analysis methods 42
Rat-to-human ortholog map 43
Gene and PTM set enrichment analysis 43
Mapping PTMs from rat to human proteins 43
Correlation of training-differential features with cell type markers 44
Immune cell type deconvolution 44
Enrichment of LM22 immune cell types 44
Quantification of sex differences in the training response 45
Comparison with human training gene expression data 45
Disease ontology enrichment analysis 46
**REFERENCES** 47

### EXPERIMENTAL DESIGN AND SAMPLE HANDLING

#### Animal handling

##### Animals

Adult male and female Fischer 344 (F344) inbred rats were obtained from the National Institute on Aging (NIA) rodent colony in three cohorts of 20-30 rats. Rats arrived on site at least 4 weeks prior to initiation of exercise training. Upon arrival at the University of Iowa, rats were adapted to a reverse dark-light cycle with lights off at 9:00 AM and lights on at 9:00 PM. Rats were housed two per cage (146.4 square inches of floor space) in ventilated racks (Thoren Maxi-Miser IVC Caging System) on Tekland 7093 Shredded Aspen bedding and fed the Lab Diet 5L79 pelleted diet. These are the standard bedding and diet used at the NIA rodent colony. The animal housing room was monitored daily and maintained at a temperature of 68-77°F and relative humidity of 25-55%. Red lights were used during the rat's dark cycle to provide adequate lighting for the staff to perform routine housing tasks and rodent handling and training. All animal procedures were approved by the Institutional Animal Care and Use Committee at the University of Iowa.

##### Treadmill familiarization and training

Treadmill exercise was performed on a Panlab 5-lane rat treadmill (Harvard Instruments, Model LE8710RTS). All animal handling and exercise was performed in the active, dark phase of the rats. Upon arrival, rats were acclimated to the reverse light cycle for a minimum of 10 days. Following the initial acclimation period, rats went through a 12-day treadmill familiarization protocol to expose the rats to the treadmill and to identify potential non-compliant rats. Those rats that were unable to run on the treadmill for 5 minutes at a speed of 10 m/min and grade of 0° were classified as non-compliant and removed from the study. Rats that successfully completed the 12-day familiarization protocol were entered in the rat database and randomized into a control or training group so that mean body weight of the groups were equal. The 8-week rats were randomly assigned to control or training within sex and tertile of weight. 4-week rats were assigned to control without randomization. 1- and 2- week rats were randomly assigned to 1- or 2-week training within sex and tertile of weight.

Exercise training began at 6 months of age in male and female rats and lasted for a duration of 1, 2, 4 or 8 weeks. Rats were exercised on the treadmill 5 days per week using a progressive training protocol designed to exercise the rats at approximately 70% of $VO_2$max. The starting treadmill speed was based on $VO_2$max measurements obtained following familiarization and prior to training in the compliant rats. Training occurred during the dark cycle of the rat, started no earlier than 10:00 AM, and ended no later than 5:00 PM over 5 consecutive days per week. Training was initiated with the treadmill set at a grade of 5°, speeds of 13m/min for males and 16m/min for females, and a duration of 20 minutes. As outlined in Table 1, the duration of exercise was increased by one minute each day until day 31 of training (start of week 7), when a duration of 50 min was reached. The treadmill grade was increased from 5° to 10° at the start of week 3 and stayed at 10° for the remainder of the training. The treadmill speed increased at the start of week 2, 4, 5, 6 and 7. At the start of week 7, speed, grade and duration were fixed and maintained for the final 10 days of the protocol to ensure steady-state had been achieved. If a rat was unable to perform at least 4 days of training per week, it was removed from the study and euthanized. Rats assigned to the control group were placed on the treadmill (0 m/min) for 15 min/day, 5 days per week, and followed a schedule similar to the 8-week training group.

Table 1. Progressive training protocol for male and female 6-month-old rats.

| **Week** | **Day** | **Speed (meters/min) Male/Female** | **Grade (degrees)** | **Duration (mins)** |
|---|---|---|---|---|
| 1 | 1 | 13/16 | 5 | 20 |
| | 2 | 13/16 | 5 | 21 |
| | 3 | 13/16 | 5 | 22 |
| | 4 | 13/16 | 5 | 23 |
| | 5 | 13/16 | 5 | 24 |
| 2 | 6 | 15/18 | 5 | 25 |
| | 7 | 15/18 | 5 | 26 |
| | 8 | 15/18 | 5 | 27 |
| | 9 | 15/18 | 5 | 28 |
| | 10 | 15/18 | 5 | 29 |
| 3 | 11 | 15/18 | 10 | 30 |
| | 12 | 15/18 | 10 | 31 |
| | 13 | 15/18 | 10 | 32 |
| | 14 | 15/18 | 10 | 33 |
| | 15 | 15/18 | 10 | 34 |
| 4 | 16 | 18/21 | 10 | 35 |
| | 17 | 18/21 | 10 | 36 |
| | 18 | 18/21 | 10 | 37 |
| | 19 | 18/21 | 10 | 38 |

| | | | | |
|---|---|---|---|---|
| | 20 | 18/21 | 10 | 39 |
| 5 | 21 | 20/23 | 10 | 40 |
| | 22 | 20/23 | 10 | 41 |
| | 23 | 20/23 | 10 | 42 |
| | 24 | 20/23 | 10 | 43 |
| | 25 | 20/23 | 10 | 44 |
| 6 | 26 | 23/26 | 10 | 45 |
| | 27 | 23/26 | 10 | 46 |
| | 28 | 23/26 | 10 | 47 |
| | 29 | 23/26 | 10 | 48 |
| | 30 | 23/26 | 10 | 49 |
| 7 | 31 | 25/28 | 10 | 50 |
| | 32 | 25/28 | 10 | 50 |
| | 33 | 25/28 | 10 | 50 |
| | 34 | 25/28 | 10 | 50 |
| | 35 | 25/28 | 10 | 50 |
| 8 | 36 | 25/28 | 10 | 50 |
| | 37 | 25/28 | 10 | 50 |
| | 38 | 25/28 | 10 | 50 |
| | 39 | 25/28 | 10 | 50 |
| | 40 | 25/28 | 10 | 50 |

#### Body composition

Body composition was determined for all rats 13 days prior to the start of training period using the minispec LF90II Body Composition Rat and Mice Analyzer (Bruker, 6.2 MHz Time Domain Nuclear Magnetic Resonance (TD-NMR) system). This analyzer was used for *in vivo* measurement of lean tissue, body fat, and body fluid in fully awake animals. Post-training body composition was determined for rats in the 4- and 8-week training groups, 5 days prior to tissue harvesting.

#### Maximum oxygen consumption ($VO_2max$)

$VO_2max$ was determined prior to the onset of training in all rats, and during the last week of training, for the 4- and 8-week exercise groups. Rats were acclimated to a single-lane enclosed treadmill (Columbus Instruments Metabolic Modular Treadmill) two days prior to testing. On the day of testing the rat was placed in the treadmill and testing began once oxygen consumption stabilized. Testing began with a warm up for 15 minutes with the treadmill set at a speed of 9 m/min and 0° incline. Following the warm up period, the incline was increased to 10° and treadmill speed was increased by 1.8 m/min every 2 minutes[1]. During the test, shock was used sparingly. Shock was used only when the rat stopped running and sat on the shock area. Testing stopped when the rat sat on the shock area 3 consecutive times and did not respond to increased shock. Upon cessation of the test, the rat was removed from the enclosure and blood was taken from the tail to measure lactate. Criteria for reaching $VO_2max$ was a leveling off of oxygen uptake despite increased workload, a respiratory exchange ratio above 1.05, and an unhemolyzed blood lactate concentration ≥6 mM.

#### Tissue collection

Tissues were collected from all rats 48 hours following the last exercise bout. On the day of collection, food was removed at 8:30 AM, three hours prior to the start of dissections, which occurred between 11:30 AM and 2:30 PM. Rats were sedated with inhaled isoflurane (1-2%). Under isoflurane anesthesia, blood was drawn via cardiac puncture. Under continued isoflurane anesthesia, the right triceps surae muscles (soleus, gastrocnemius, and plantaris), subcutaneous white fat on the right side, right lobe of the liver, heart, and lungs were removed in that specific order. Removal of the heart resulted in death. Immediately following removal of the heart, a guillotine was used for decapitation. The brain was removed from the skull, and the following regions were dissected out in the specified order: hypothalamus, right and left hippocampus, right and left cerebral cortex. Following decapitation, specific organs were removed in the following order: right kidney, right & left adrenal glands, spleen, brown adipose tissue, small intestine (jejunum), colon (transverse and descending) and feces, right testes or ovaries, right vastus lateralis, and tibia. All tissues were flash-frozen in liquid nitrogen immediately upon removal, placed in cryovials, and stored at -80°C. All tissues were subsequently shipped on dry ice to the Biospecimens Repository at the University of Vermont.

### Sample distribution

#### Archiving

Frozen tissue samples received by the Biospecimens Repository at the University of Vermont were logged in to Freezerworks (Dataworks Development Inc., Mountlake Terrace, Washington) and stored in -80°C freezers. All samples were labeled with a unique 11-digit barcode.

#### Selection of animals for molecular assays

Tissue samples were collected for between 12 and 20 animals per sex, per training group (i.e., sedentary controls or animals trained for 1, 2, 4, or 8 weeks). Six animals per sex per training group were randomly selected for molecular profiling. All six replicates were used for proteomic assays; a subset of five replicates were used for genomic and metabolomic assays. Additional rats were selected at random if there were insufficient aliquots for specific molecular assays, as was the case for the adrenal gland, brown adipose tissue, and ovary immunoassays.

#### Cryopulverization

Tissues samples were stored at -80°C until time of processing. Frozen tissue samples were transferred from storage vials to Covaris tissueTUBEs (Covaris, Inc, Woburn, Massachusetts) flash-chilled in liquid nitrogen. Larger tissues were first broken into smaller pieces on a chilled, foil-covered stainless steel block using a foil-covered hammer or pestle. The tissue piece(s) were centered in the tissueTUBE pouch (primary impact zone), and a pre-chilled glass transfer tube was attached. Filled tissueTUBEs were placed on dry ice while the Covaris CryoPREP CP02 (Covaris Inc, Woburn, Massachusetts) was set to the appropriate setting for the tissue type. TissueTUBEs were dipped in liquid nitrogen, and the glass transfer tube was loosened ¼ turn to prevent pressure build-up inside the tissueTUBE before being inserted in the Covaris CP02 to pulverize. Tissues requiring an additional round of pulverization were first inspected to ensure tissueTUBE integrity and dipped in liquid nitrogen before being put back into the Covaris CP02. After pulverization, tissueTUBEs were dipped in liquid nitrogen and then inverted to move the pulverized sample into the glass transfer tube. Pulverized samples were transferred to cryogenic storage vials for long term storage at -80°C.

#### Aliquoting

Frozen tissue storage vials were removed from the -80°C freezer and placed on dry ice. Aliquot vials were set up in a prechilled CoolRack XT CFT24 (Corning, Corning, New York) sitting in dry ice inside an AirClean 600 Dead Air Workstation (AirClean Systems, Creedmoor, North Carolina). Working with one storage vial at a time, disposable plastic transfer scoops of predetermined volumes were chilled in liquid nitrogen and then used to measure and transfer tissue to aliquot vials. A chilled metal spatula was used to break up any tissue clumps in the storage tubes before aliquoting. Aliquot vials were capped and stored at -80°C before shipping to chemical analysis labs. By experimentation we determined that the target weights could be aliquoted with a reproducibility of +/- 15% of the target. Different sized scoops were needed for different tissues depending on fat content.

#### Reference standards

Aliquots of assay- and tissue-specific reference standards were included in molecular assays in order to evaluate technical differences across batches. The pooled reference standards used for RNA-Seq, ATAC-seq, RRBS, and immunoassays were from non-compliant animals collected at the University of Florida for a separate MoTrPAC study. Samples from the same tissue and sex were pulverized together to create a homogenous pool. One aliquot each of the male and female reference standards was included on each plate of study samples. Tissue-matched reference standards were used when possible. Metabolomics and lipidomics reference standards were from pilot samples collected at the University of Florida, Joslin Diabetes Center, and the University of Iowa for a separate MoTrPAC study. Samples were split to create pools of sedentary and immediate-post-exercise reference standards. One aliquot each of the sedentary and exercised reference standards was included with each batch of study samples. For proteomics, a universal reference standard was generated within each tissue type by pooling equal amounts of peptide from each experimental sample. All samples were from Fischer 344 (F344) inbred rats obtained from the NIA rodent colony.

### DATA PRODUCTION AND QUANTIFICATION

#### DNA methylation by reduced representation bisulfite sequencing

##### DNA extraction and library preparation

Reduced representation bisulfite sequencing (RRBS)[2] was performed at the Icahn School of Medicine at Mount Sinai. Rat tissues (10-30 mg of white adipose tissue) were disrupted in GenFind v2 lysis buffer (Beckman Coulter, Indianapolis, IN) using a tissue ruptor (Omni International, Kennesaw, GA) and the genomic DNA was extracted in a BiomekFx automation workstation (Beckman Coulter, Chaska, Minnesota), according to the manufacturer's instructions. Two tissue-specific consortium reference standards were included to monitor the sample processing QC. DNA samples were quantified by Qubit assay (dsDNA HR assay, Thermo Fisher Scientific), and the quality was determined by the Nanodrop A260/280 and A260/230 ratios.

The Ovation® RRBS Methyl-Seq kit from Tecan Genomics (Baldwin Park, CA) was used to generate the RRBS libraries according to the manufacturer's instructions. All library preparations were carried out in an automated workstation (Biomek Fx, Beckman Coulter). Briefly, purified genomic DNA (100 ng) was digested by the methylation-insensitive restriction enzyme MspI to generate short fragments that contain CpG dinucleotides at the ends. After end-repair, A-tailing, and ligation to methylated Illumina adapters, the CpG-rich DNA fragments (40-220 base pairs) were size-selected and subjected to bisulfite conversion according to the manufacturer's instructions. The single-stranded uracil-containing DNA was purified after desulfonation using the magnetic beads provided in the kit, and eluted DNA was converted to double-stranded DNA by PCR with the following cycling parameters: 95°C for 2 minutes, followed by the optimal number of cycles of 95°C for 15 seconds, 60°C for 1 minute, and 72°C for 30 seconds, and a final step at 72°C for 10 minutes. The optimal number of cycles for the enrichment PCR was calculated based on qPCR values. A final clean-up step was performed by 1x AMPure XP beads. The quantity of the libraries was measured by Qubit High Sensitivity assays (ThermoFisher Scientific), and the quality was evaluated using Bioanalyzer High Sensitivity DNA Chip (Agilent Technologies, Santa Clara, CA). The libraries were pooled and sequenced in a MiSeq Nano flow cell to determine the sequencing quality of the libraries. Unmethylated lambda DNA (Promega, D1521) was spiked (0.05%) in each sample to monitor bisulfite conversion. Jurkat genomic DNA (ThermoFisher, SD1111) and CpG Methylated Jurkat genomic DNA (ThermoFisher, SD1121) were used as positive controls with each batch.

#### DNA sequencing, data processing, and normalization

Sequencing of RRBS libraries was performed on a NovaSeq 6000 platform (Illumina, San Diego, CA, USA) using a paired-end 100 base-pair run configuration. The pooled libraries were spiked in with 10% PhiX and sequenced to a minimal depth of 30 million paired-end reads per library using a custom 1-index primer as per Illumina guidelines. In addition to the 8-base barcode, the adapter contained 8-base unique molecular identifiers (UMIs) immediately following the library index. The UMIs were used for duplicate read determination (see below). To take advantage of this feature, the libraries were sequenced using 16 cycles for the i7 index read.

Reads were demultiplexed with bcl2fastq (version 2.20) using options --use-bases-mask Y*,I8Y*,I*,Y* --mask-short-adapter-reads 0 --minimum-trimmed-read-length 0 (Illumina, San Diego, CA, USA), and UMIs in the index FASTQ files were attached to the read FASTQ files. The regular 5' and 3' adapters were trimmed with TrimGalore (v1.18), and the diversity adapter that is about 0 to 3 bases of RDD (R={A or G} and D={A, G, or T}) that is added before YGG (Y={C or T} depending on the methylation) from the YGG MspI cut signature was trimmed with the NuGEN script "trimRRBSdiversityAdaptCustomers.py" (https://github.com/nugentechnologies/NuMetRRBS). FastQC (v0.11.8) was used to generate pre-alignment QC metrics[3]. Bismark (v0.20.0) was used to index and align reads to release 96 of the Ensembl *Rattus norvegicus* (rn6) genome and gene annotation[4]. As the lambda genome was spiked into each sample to determine the bisulfite conversion efficiency, the lambda genome (GenBank: J02459.1) was also indexed. Default parameters were used for Bismark's bismark_genome_preparation in the alignment step. Bismark output BAM files were first formatted using a custom script; Bismark's deduplicate_bismark with "-p --barcode" options was used to remove PCR duplicates from the bam files; and Bismark's "bismark_methylation_extractor --comprehensive --bedgraph" was used to quantify methylated and unmethylated coverages for all the CpG sites. Bowtie 2 (v2.3.4.3) was used to index and align reads to globin, rRNA, and phix sequences in order to quantify the percent of reads that mapped to these contaminants and spike-ins[5]. SAMtools (v1.3.1) was used to compute mapping percentages to different chromosomes[6]. UMIs were used to accurately quantify PCR duplicates with NuGEN's "nodup.py" script (https://github.com/tecangenomics/nudup). QC metrics from every stage of the quantification pipeline were compiled, in part with multiQC (v1.6)[7]. The openWDL-based implementation of the RRBS pipeline on Google Cloud Platform is available on GitHub (https://github.com/MoTrPAC/motrpac-rrbs-pipeline). Only CpG sites with methylation coverage of >=10 in all samples were included for downstream analysis, and normalization was performed separately in each tissue. Individual CpG sites were divided into 500 base-pair windows and were clustered using the Markov Clustering algorithm via the *MCL* R package[8]. To apply MCL, for each 500 base-pair window an undirected graph was constructed, linking individual sites if their correlation was >=0.7. MCL was chosen for this task as it: (1) determines the number of clusters internally, (2) identifies homogeneous clusters, and (3) keeps single sites that are not correlated with either sites as singletons (clusters of size one). The resulting sites/clusters were used as input for normalization and differential analysis with edgeR[9]. To generate normalized sample-level data, the methylation coverages of filtered sites/clusters were first $log_2$-transformed, and normalization was performed using preprocessCore's quantile normalization *preprocessCore::normalize.quantiles.robust*[10].

### Assay for transposase-accessible chromatin using sequencing

#### Nuclei extraction and library preparation

Assay for transposase-accessible chromatin using sequencing (ATAC-seq)[11] was performed at Stanford University and the Icahn School of Medicine at Mount Sinai. Only Stanford ATAC-seq data were used in this manuscript. Nuclei from aliquoted tissue samples (30 mg for white adipose, 15 mg for brown adipose, 10 mg for hippocampus, kidney, lung, gastrocnemius, heart, and liver tissues) were extracted using the Omni-ATAC protocol with modifications[12]. Two tissue-specific consortium reference standards were included for sample processing QC. The white adipose, brown adipose, and hippocampus tissues were processed using no-douncing nuclei extraction to prevent fat droplets from interfering with the subsequent transposition steps. Tissue powder was incubated in the homogenization buffer for 10 min at 4°C. The tube was inverted every 2-3 minutes. The heart, liver, kidney, lung, and gastrocnemius tissues were processed using the Omni-ATAC protocol with modifications. Tissue powder was incubated in the homogenization buffer for 5 minutes on ice and dounced 10 times using pestle A and 20 times with pestle B. For both protocols, the homogenate passed through a 40 µm cell strainer to collect nuclei. Nuclei were stained with DAPI and counted using an automated cell counter. 50,000 nuclei (or max. 500 µl nuclei) were added to 1 ml ATAC-RSB buffer and spun at 1000 g for 10 minutes, and the supernatant was removed. The nuclei pellet was resuspended in 50 µl of transposition mixture and incubated at 37°C for 30 minutes with 1000 rpm shaking. The transposed DNA was purified using Qiagen MinElute Purification kits (Qiagen # 28006). The DNA product was amplified using NEBnext High-Fidelity 2x PCR Master Mix (NEB, M0541L) and custom indexed primers. 1.8x SPRIselect beads were used to clean the PCR reaction and obtain the ATAC-seq library for sequencing.

#### DNA sequencing, data processing, and normalization

Pooled libraries were sequenced on an Illumina NovaSeq 6000 platform (Illumina, San Diego, CA, USA) to a target depth of 35 million read pairs (70 million paired-end reads) per sample using a paired-end 50 base-pair run configuration. Reads were demultiplexed with bcl2fastq2 (v2.20.0) (Illumina, San Diego, CA, USA). Data was processed with the ENCODE ATAC-seq pipeline (v1.7.0) (https://github.com/ENCODE-DCC/atac-seq-pipeline)[13]. Samples from a single sex and training time point, e.g., males trained for 2 weeks, were analyzed together as biological replicates in a single workflow. Briefly, adapters were trimmed with cutadapt v2.5[14] and aligned to release 96 of the Ensembl *Rattus norvegicus* (rn6) genome[4] with Bowtie 2 v2.3.4.3[5]. Duplicate reads and reads mapping to the mitochondrial chromosome were removed.

Signal files and peak calls were generated using MACS2 v2.2.4[15], both from reads from each sample and pooled reads from all biological replicates. Pooled peaks were compared with the peaks called for each replicate individually using Irreproducibility Discovery Rate[16] and thresholded to generate an optimal set of peaks. The cloud implementation of the ENCODE ATAC-seq pipeline and source code for the post-processing steps are available at https://github.com/MoTrPAC/motrpac-atac-seq-pipeline. Optimal peaks (overlap.optimal_peak.narrowPeak.bed.gz) from all workflows were concatenated, trimmed to 200 base pairs around the summit, and sorted and merged with bedtools v2.29.0[17] to generate a master peak list. This peak list was intersected with the filtered alignments from each sample using bedtools coverage with options -nonamecheck and -counts to generate a peak by sample matrix of raw counts. The remaining steps were applied separately on raw counts from each tissue. Peaks from non-autosomal chromosomes were removed, as well as peaks that did not have at least 10 read counts in four samples. Filtered raw counts were then quantile-normalized with limma-voom[18]. This version of the normalized data was used for downstream analyses.

### RNA sequencing

#### Extraction of total RNA

RNA sequencing (RNA-Seq) was performed at Stanford University and the Icahn School of Medicine at Mount Sinai. Rat tissues (30 mg for white adipose, 15 mg for brown adipose, 10 mg for other solid tissues, and 0.47 ml blood) were disrupted in Agencourt RNAdvance tissue lysis buffer (Beckman Coulter, Brea, CA) using a tissue ruptor (Omni International, Kennesaw, GA, # 19-040E). Total RNA was extracted in a BiomekFx automation workstation according to the manufacturer’s instructions for tissue-specific extraction. Total RNA from blood collected in PAXgene tubes (BD Biosciences, Franklin Lakes, NJ, # 762165) was extracted using the Agencourt RNAdvance blood specific kit (Beckman Coulter). Two tissue-specific consortium reference standards were included to monitor the sample processing QC. The RNA was quantified by NanoDrop (ThermoFisher Scientific, # ND-ONE-W) and Qubit assay (ThermoFisher Scientific), and the quality was determined by either Bioanalyzer or Fragment Analyzer analysis.

#### mRNA Sequence Library Preparation

Universal Plus mRNA-Seq kit from NuGEN/Tecan (# 9133) were used for generation of RNA-Seq libraries derived from poly(A)-selected RNA according to the manufacturer’s instructions. Universal Plus mRNA-Seq libraries contain dual (i7 and i5) 8 bp barcodes and an 8 bp unique molecular identifier (UMI), which enable deep multiplexing of NGS sequencing samples and accurate quantification of PCR duplication levels. Approximately 500ng of total RNA were used to generate the libraries. The Universal Plus mRNA-Seq workflow consists of poly(A) RNA selection, RNA fragmentation and double-stranded cDNA generation using a mix-ture of random and oligo(dT) priming, end repair to generate blunt ends, adaptor ligation, strand selection, AnyDeplete workflow to remove unwanted ribosomal and globin transcripts, and PCR amplification to enrich final library species. All library preparations were performed using a

Biomek i7 laboratory automation system (Beckman Coulter). Tissue-specific reference standards provided by the consortium were included with all RNA isolations to QC the RNA.

##### RNA sequencing, quantification, and normalization

Pooled libraries were sequenced on an Illumina NovaSeq 6000 platform (Illumina, San Diego, CA, USA) to a target depth of 35 million read pairs (70 million paired-end reads) per sample using a paired-end 100 base pair run configuration. In order to capture the 8-base UMIs, libraries were sequenced using 16 cycles for the i7 index read and 8 cycles for the i5 index read. Reads were demultiplexed with bcl2fastq2 (v2.20.0) using options --use-bases-mask Y*,I8Y*,I*,Y* --mask-short-adapter-reads 0 --minimum-trimmed-read-length 0 (Illumina, San Diego, CA, USA), and UMIs in the index FASTQ files were attached to the read FASTQ files. Adapters were trimmed with cutadapt (v1.18), and trimmed reads shorter than 20 base pairs were removed[14]. FastQC (v0.11.8) was used to generate pre-alignment QC metrics[3]. STAR (v2.7.0d) was used to index and align reads to release 96 of the Ensembl Rattus norvegicus (rn6) genome and gene annotation[4]. Default parameters were used for STAR's genomeGenerate run mode; in STAR's alignReads run mode, SAM attributes were specified as NH HI AS NM MD nM, and reads were removed if they did not contain high-confidence collapsed splice junctions (--outFilterType BySJout). RSEM (v1.3.1) was used to quantify transcriptome-coordinate-sorted alignments using a forward probability of 0.5 to indicate a non-strand-specific protocol[19]. Bowtie 2 (v2.3.4.3) was used to index and align reads to globin, rRNA, and phix sequences in order to quantify the percent of reads that mapped to these contaminants and spike-ins[5]. UCSC's gtfToGenePred was used to convert the rn6 gene annotation (GTF) to a refFlat file in order to run Picard CollectRnaSeqMetrics (v2.18.16) with options MINIMUM_LENGTH=50 and RRNA_FRAGMENT_PERCENTAGE=0.3[20]. UMIs were used to accurately quantify PCR duplicates with NuGEN's "nodup.py" script (https://github.com/tecangenomics/nudup). QC metrics from every stage of the quantification pipeline were compiled, in part with multiQC (v1.6)[7]. The openWDL-based implementation of the RNA-Seq pipeline on Google Cloud Platform is available at https://github.com/MoTrPAC/motrpac-rna-seq-pipeline. Filtering of lowly expressed genes and normalization were performed separately in each tissue. RSEM gene counts were used to remove lowly expressed genes, defined as having 0.5 or fewer counts per million in all but one sample. These filtered raw counts were used as input for differential analysis with DESeq2[21], as described below. To generate normalized sample-level data, filtered gene counts were TMM-normalized using *edgeR::calcNormFactors*, followed by conversion to log counts per million with *edgeR::cpm*[9].

#### Liquid chromatography tandem mass spectrometry (LC-MS/MS) proteomics

##### Tissue lysis and protein extraction

LC-MS/MS analyses of 60 samples per tissue representing full time course samples for 6 female and 6 male rats were performed at the Broad Institute of MIT and Harvard (heart and liver) and the Pacific Northwest National Laboratory (gastrocnemius, white adipose tissue, cerebellum, lung, and kidney). Sample processing for proteomic analysis was based on a protocol previously described[22], with modifications detailed below. Cryopulverized tissue samples were suspended in cold, freshly-prepared lysis buffer (8 M urea (Sigma-Aldrich, St. Louis, Missouri), 50 mM Tris pH 8.0, 75 mM sodium chloride, 1 mM EDTA, 2 µg/ml Aprotinin (Sigma-Aldrich, St. Louis, Missouri), 10 µg/ml Leupeptin (Roche CustomBiotech, Indianapolis, Indiana), 1 mM PMSF in EtOH, 10 mM sodium fluoride, 1% phosphatase inhibitor cocktail 2 and 3 (Sigma-Aldrich, St. Louis, Missouri)), vortexed for 10 seconds, and incubated for 15 min on a thermomixer set to 4°C and 800 rpm. The lysis buffer for heart and liver tissues was supplemented with deacetylase inhibitors for acetylome analysis (10 mM Sodium Butyrate, 2 µM SAHA, and 10 mM nicotinamide). Samples were vortexed for an additional 10 seconds and incubated for 15 more minutes with the same settings, then centrifuged for 10 minutes at 4°C and 18000 rcf to remove debris. Supernatant was removed, and protein concentrations were determined by BCA assay (ThermoFisher, Waltham, Massachusetts).

#### Protein digestion

Protein lysate concentrations were normalized within samples of the same tissue type, and protein was reduced with 5 mM dithiothreitol (DTT, Sigma-Aldrich) for 1 hour in a thermomixer set to 37°C and 1000 rpm. Protein was then alkylated with iodoacetamide (IAA, Sigma-Aldrich) for 45 minutes in the dark in a thermomixer set to 25°C and 1000 rpm. Samples were then diluted 1:4 with 50 mM Tris-HCl, pH 8.0 to lower the urea concentration below 2 M. LysC endopeptidase (1mAU/uL, Wako Chemicals, Richmond, Virginia) was added at a 1:50 enzyme:substrate ratio, and samples were then digested for 2 hours in a thermomixer set to 25°C and 850 rpm. Sequencing grade modified trypsin (Promega, MAdison, Wisconsin) was then added at an enzyme:substrate ratio of 1:50 (or 1:10 for white adipose tissue), and samples were digested for 14 hours in a thermomixer set to 25°C and 850 rpm. Digestions were quenched by adding formic acid (FA) to a final concentration of 1%, and samples were centrifuged at 1500 rcf for 15 minutes at 4°C. Supernatants were collected into new tubes, diluted to 3 ml total volume with 0.1% FA, and desalted using Sep-Pac C18 SPE cartridges (Waters, Milford, MA). Clean peptides were concentrated in a speedvac, and peptide concentrations were determined by BCA assay.

#### Tandem mass tag (TMT) labeling of peptides

400 µg aliquots of every sample were prepared and dried down to be used for TMT labeling. Additionally, a common reference sample was generated within each tissue type by pooling equal amounts of peptide from each experimental sample; one 400 µg aliquot of this reference was included in each multiplex. Within each tissue type male and female rat time course samples were randomized into 6 TMT11 plexes making sure single time course samples are in the same plex and keeping 1 male and 1 female in each plex. Samples were resuspended in 200 mM HEPES pH 8.5 to a final concentration of 5 µg/uL for labeling and a reduced TMT reagent labeling approach was used[23]. Experimental samples were randomized across the first ten TMT channels (126C to 131N) and common reference aliquots were assigned to the last TMT channel (131C). TMT reagents were resuspended in anhydrous acetonitrile to a

concentration of 20 µg/uL, and 400 µg of reagent was added to appropriate peptide aliquots (resulting in a 1:1 peptide:tag ratio). Labeling was allowed to proceed for 1 hour in a thermomixer set to 25°C and 400 rpm. After 1 hour, samples were diluted to 2.5 µg/uL with 20% acetonitrile, a small aliquot was removed from each sample for labeling efficiency and mixing tests, and the remaining sample was snap-frozen in liquid $N_2$ and stored at -80°C. Following QC checks, reactions were quenched with hydroxylamine and samples from each multiplex were combined, concentrated in a speedvac, and desalted using Sep-Pac C18 SPE cartridges (Waters). Heart and liver samples were subjected to phosphotyrosine enrichment, while this step was skipped for the remaining tissues to perform offline bRP fractionation.

#### Phosphotyrosine peptide enrichment

Phosphotyrosine (pY) enrichment was performed on liver and heart tissues for improved characterization of tyrosine kinase signaling pathways, as described previously[24]. TMT-labeled and combined peptides were resuspended in 1.5 ml of IAP buffer (50 mM MOPS, 10 mM sodium phosphate dibasic, 50 mM NaCl) and cleared by centrifugation (5 min, 5000 rcf). The pellet was dissolved in 1% FA and combined with the pY enrichment flowthrough for downstream bRP fractionation, while the supernatant was used for pY enrichment. PTMScan Phospho-Tyrosine Rabbit mAb beads (Cell Signaling Technologies #8803, Danvers, Massachusetts) were washed three times with 1.5 ml IAP buffer (30 s, 2000 rcf) and incubated with the solubilized peptides (1 hour, 4°C, end-over-end rotation). Beads with pY peptides bound were centrifuged (1 minute, 1500 rcf), and the flowthrough was collected, desalted using Sep-Pac C18 SPE cartridges, and used for downstream bRP offline fractionation. The beads were washed four times using 1.5 ml PBS (30 seconds, 2000 rcf) and pY peptides were eluted by two consecutive 5-minute incubations with 0.15% trifluoroacetic acid (TFA). Eluted pY peptides were desalted using stage tips containing 2x C18 discs (Empore, CDS Analytical, Oxford, Pennsylvania). The stage tip column was conditioned with 1x 100 µL methanol, 1x 100 µL 50% acetonitrile (ACN) / 1% FA, and 2x 100 µL 0.1% FA washes. Acidified peptides were bound to the column, washed with 2x 100 µL 0.1% FA, and eluted with 50 µL 50% ACN / 0.1% FA. Eluted peptides were dried using a vacuum concentrator. Phosphotyrosine peptides were suspended in 9 µl of 0.1% FA and 3% ACN, and 2 x 4 µl were injected in an LC-MS/MS instrument.

#### Offline bRP fractionation

Each combined multiplex sample was fractionated by high pH reversed phase separation using a 3.5 µm Agilent Zorbax 300 Extend-C18 column (4.6 mm ID x 250 mm length). Samples were resuspended in mobile phase A (5 mM ammonium formate, pH 10, in 2% acetonitrile), centrifuged to remove debris, loaded onto the column and eluted off the column for 96 minutes at a flow rate of 1 ml/minute using mobile phase B (5 mM ammonium formate, pH 10, in 90% acetonitrile) with the following gradient (time(min): %B): 0:0; 7:0; 13:16; 73:40; 77:44; 82:60; 96:60. A total of 96 fractions were collected, immediately acidified to 0.1% formic acid, and concatenated down to 24 fractions by combining non-sequential fractions. For heart and liver tissues, the complete gradient was collected and only fractions 15-91 showing robust peptide signals were concatenated. Fractions 3-14 were collected and used as an extra fraction “A” for phosphopeptide enrichment. For the remaining tissues, 96 fractions were collected between elution time of 2.4 to 91 minutes and all fractions were used for concatenation. Five percent was removed for global proteome analysis. For phosphopeptide enrichment, the remaining 95% was further concatenated to 12 fractions plus an additional fraction “A” for heart and liver only. These fractions were frozen in liquid nitrogen, vacuum-centrifuged to dryness, and stored at -80°C until ready for phosphopeptide enrichment.

#### Phosphopeptide enrichment

Phosphopeptide enrichment was performed through immobilized metal affinity chromatography (IMAC) using $Fe^{3+}$-NTA-agarose beads, freshly prepared from Ni-NTA-agarose beads (Qiagen, Hilden, Germany) by sequential incubation in 100 mM EDTA to strip nickel, washing with HPLC water, and incubation in 10 mM iron (III) chloride). Peptide fractions were resuspended to 0.5 µg/uL in 80% ACN + 0.1% TFA and incubated with beads for 30 minutes in a thermomixer set to 1000 rpm at room temperature. After 30 minutes, beads were spun down (1 minute, 1000 rcf) and supernatant was removed and saved as flow-through for subsequent enrichments. Phosphopeptides were eluted off IMAC beads in 3x 70 µl of agarose bead elution buffer (500 mM $K_2HPO_4$, pH 7.0), desalted using C18 stage tips, eluted with 50% ACN, and lyophilized. Samples were then reconstituted in 3% ACN / 0.1% FA for LC-MS/MS analysis (12 µl reconstitution / 5 µl injection for gastrocnemius, white adipose, lung, kidney, and cortex samples; 9 µl reconstitution / 4 µl injection for heart and liver samples). Flow-through from the 12 IMAC fractions were further concatenated into 4 fractions to be used for acetylpeptide enrichment only for heart and liver tissues.

#### Acetylpeptide enrichment

Acetylated lysine peptides were enriched using an antibody against the acetyl-lysine motif (Cell Signaling Technologies #13416, Danvers, Massachusetts). Peptide fractions were reconstituted with 1.4 ml of IAP buffer per fraction and incubated for 2 h at 4°C with pre-washed (3 times with IAP buffer) agarose beads bound to acetyl-lysine motif antibody. Peptide-bound beads were washed 4 times with ice-cold PBS followed by elution with 2 × 100 µl of 0.15% TFA. Eluents were desalted using C18 stage tips, eluted with 50% ACN, lyophilized, and reconstituted in 9 µl 3% ACN / 0.1% FA. Four µl of each fraction were injected for LC-MS/MS analysis.

#### K-ε-GG enrichment for ubiquitylome analysis

Ubiquitin enrichment was performed based on the UbiFast protocol[25]. Anti-K-ε-GG bead-bound antibodies from the PTM-Scan ubiquitin remnant motif kit (Cell Signaling Technologies #5562) were cross-linked using the following procedure. Beads were washed 3 times with 100 mM sodium borate (pH 9.0) and incubated with 20 mM DMP for 30 minutes at room temperature. Beads were then washed 2 times with 200 mM ethanolamine and incubated overnight at 4°C in 200 mM ethanolamine with end-over-end rotation. Following incubation, beads were washed three times with an IAP buffer and stored at 4°C at a concentration of 0.5 µg/µL. For each 11-plex experiment, 31.25 µg of cross-linked anti-K-ε-GG bead-bound antibody at 0.5 µg/µL in IAP per channel was aliquoted into 1.5 ml Eppendorf tubes on ice. Eight hundred and one thousand micrograms of total peptides were used for liver and heart tissue samples, respectively. Each sample was reconstituted to 0.5 mg/ml concentration in IAP buffer and vortexed for 10 minutes. Peptides were then centrifuged for 5 minutes at 5000 g. Each peptide solution was added to a tube of antibody and gently rotated end-over-end at 4°C for 1 hour. Following enrichment, samples were centrifuged (1 minute, 2000 rcf) and the supernatant was removed. Beads were washed with 1.5 ml ice cold IAP followed by 1.5 ml ice cold PBS (30 seconds, 2000 rcf) and reconstituted in 200 µl 100 mM HEPES buffer. For each sample, 400 µg of TMT labeling reagent in 10 µl acetonitrile was added. For heart samples, 800 µg of TMT labeling reagent was added to improve labeling efficiency. Peptides were TMT labeled on-beads while shaking vigorously (1400 rpm) at 20°C for 10 minutes, then quenched with 8 µl 5% hydroxylamine and shaken vigorously for another 5 minutes, washed once with 1.3 ml cold IAP, and again with 1.5 ml cold IAP. Samples for each channel were resuspended and transferred to a combination tube with 130 µl cold IAP. Following combination, each now-empty tube was serially washed with 1.5 ml cold IAP to remove remaining beads. This 1.5 ml IAP fraction was added to the combination tube and used to wash the combined beads. Combined beads were then washed one final time with 1.5 ml ice cold PBS. Once the channels were combined and washed, peptides were eluted twice from the beads by resuspending with 150 µL room-temperature 0.15% TFA and incubating for 5 minutes at room temperature. Each round of acid-eluted K-ε-GG-modified peptides was desalted on an equilibrated two-punch C18 stage tip. Both elutions of K-ε-GG peptides were loaded sequentially, washed 2 times with 100 µl 0.1% FA, and eluted into an LC-MS vial with 50 µl 50% ACN / 0.1% FA. The eluted peptides were frozen, lyophilized, and reconstituted in 9 µl 3% ACN / 0.1% FA, with 4 µl injected twice for two consecutive LC-MS/MS runs.

#### LC-MS/MS analysis of heart and liver tissue samples

Online separation was conducted with a nanoflow Proxeon EASY-nLC 1200 UHPLC system (Thermo Fisher Scientific). In this setup, the LC system, column, and platinum wire used to deliver the electrospray source voltage were connected via a stainless steel cross (360 mm, IDEX Health & Science, UH-906x). An in-house packed 22 cm x 75 µm internal diameter C18 silica picofrit capillary column (1.9 mm ReproSil-Pur C18-AQ beads, Dr. Maisch GmbH, r119.AQ; Picofrit 10 µm tip opening, New Objective, PF360-75-10-N-5) heated at 50°C using a column heater sleeve (Phoenix-ST) was used for chromatographic separation. For global proteome analysis, ~1 µg was loaded on-column in a 2-µL volume (based on peptide-level BCA with uniformly-distributed fractionation presumed). For phosphoproteome, acetylome, and ubiquitylome analysis, 50% of each fraction sample was injected in a 4-µl volume. Mobile phase flow rate was 200 nL/min, composed of 3% acetonitrile / 0.1% formic acid (Solvent A) and 90% acetonitrile / 0.1% formic acid (Solvent B). The 110-minute LC-MS/MS method used for global proteome and IMAC phosphoproteome analysis consisted of a 10-minute column-equilibration procedure; a 20-minute sample-loading procedure; and the following gradient profile: (time (minutes):%B) 0:1; 1:6; 63:20, 85:30; 94:60; 95;90; 100:90; 101:50; 110:50. For acetylproteome, ubiquitylome, and phosphotyrosine enrichment analysis, the same LC and column setup was used, but the gradient was extended to 154 minutes with the following gradient profile: (time (minutes):%B) 0:2; 2:6; 122:30; 130:60; 133;90; 143:90; 144:50; 154:50.

For proteome analysis, samples were analyzed with a Q-Exactive Plus mass spectrometer (Thermo Fisher Scientific). Data-dependent MS/MS acquisition was performed using the following relevant parameters: positive ion mode at a spray voltage of 1.8 kV, MS1 resolution of 60,000, an AGC target of 3e6, a mass range from 300 to 1800 *m/z*, MS2 resolution of 45,000, MS2 AGC target of 5e4, isolation window of 0.7 *m/z*, MS2 maximum injection time of 105 ms, HCD collision energy of 29%, dynamic exclusion for 20 seconds, peptide match was set to preferred for monoisotopic peak determination, charge states 2-6.

For analysis of IMAC-enriched peptides (phosphoproteome), samples were analyzed with a Q-Exactive HFX mass spectrometer (Thermo Fisher Scientific). Data-dependent MS/MS acquisition was performed using the following relevant parameters: positive ion mode at a spray voltage of 1.5 kV, MS1 resolution of 60,000, an AGC target of 3e6, a mass range from 350 to 1800 *m/z*, MS2 resolution of 45,000, MS2 AGC target of 5e4, isolation window of 0.7 *m/z*, MS2 maximum injection time of 105 ms, HCD collision energy of 31%, dynamic exclusion for 15 seconds, peptide match was set to preferred for monoisotopic peak determination, charge states 2-6. The advanced precursor determination feature (APD)[26] was turned off in the tune file using a software patch provided by Thermo Fisher Scientific (Tune version 2.11). For analysis of phosphotyrosine enriched peptides (phosphoproteome), the same settings were used with the following modifications: MS2 AGC target of 2e5 and MS2 maximum injection time of 150 ms. For analysis of acetyl-lysine enriched peptides (acetylome), the same settings were used with the following modifications: MS2 AGC target of 5e5 and MS2 maximum injection time of 150 ms. For analysis of K-ε-GG enriched peptides (ubiquitylome), the same settings were used with the following modifications: MS2 AGC target of 5e4 and MS2 maximum injection time of 150 ms.

#### LS-MS/MS analysis of the gastrocnemius, white adipose tissue, cortex, kidney, and lung

For global proteome analysis, online separation was performed with a nanoAcquity M-Class UHPLC system (Waters) equipped with a 250 mm x 4.6 mm internal diameter Jupiter 5 µm C18 trapping column (Phenomenex). An in-house packed, 25 cm x 75 µm internal diameter C18 silica picofrit column (1.7 µm UPLC BEH particles, Waters Acquity) heated to 50°C was used for chromatographic separation. 0.25-0.5 µg per fraction (based on peptide-level BCA) was loaded on-column in a 5uL volume. Mobile phase flow rate was 200 nL/min, composed of 0.1% formic acid in $H_2O$ (Solvent A) and 0.1% formic acid in acetonitrile (Solvent B). Following a 6-minute trap time and 27-minute sample load time in 100% solvent A, the following 120-minute gradient profile was applied: (minutes:%B) 0:8; 83:20; 96:35; 101:75; 104:95; 110:95; 111:50; 113:95. Samples were analyzed with a Q Exactive HF mass spectrometer (Thermo Fisher Scientific). Data-dependent MS/MS acquisition was performed with the following parameters: positive ion mode at a spray voltage of 1.8 kV, MS1 resolution of 60,000, an AGC target of 3e6, a mass range from 300 to 1800 *m/z*, MS2 resolution of 30,000, MS2 AGC target of 1e5, isolation window of 0.7 *m/z*, MS2 maximum injection time of 100 ms, HCD collision energy of 30%, dynamic exclusion for 45 seconds, peptide match was set to preferred for monoisotopic peak determination, charge states 2-6.

For IMAC-enriched phosphopeptide analysis, online separation was performed with a Dionex Ultimate 3000 UHPLC direct-inject system (Thermo). An in-house packed, 30 cm x 75 µm internal diameter C18 silica picofrit column (1.7 µm UPLC BEH particles, Waters Acquity) at room temperature was used for chromatographic separation. Samples were resuspended in 12 µl Solvent A, and 5 µl was loaded on-column. Mobile phase flow rate was 200 nL/min, composed of 0.1% formic acid in $H_2O$ (Solvent A) and 0.1% formic acid in acetonitrile (Solvent B). Following a 60-minute sample load time in 100% solvent A, the following 120-minute gradient profile was applied: (time (minutes):%B) 0:8; 85:25; 95:35; 100:75; 105:5; 110:95; 115:1. Samples were analyzed with a Q-Exactive HF-X mass spectrometer (Thermo Fisher Scientific). Data-dependent MS/MS acquisition was performed with the following parameters: positive ion mode at a spray voltage of 1.8 kV, MS1 resolution of 60,000, an AGC target of 3e6, a mass range from 300 to 1800 *m/z*, MS2 resolution of 45,000, MS2 AGC target of 2e5, isolation window of 0.7 *m/z*, MS2 maximum injection time of 100 ms, HCD collision energy of 30%, dynamic exclusion for 45 seconds, peptide match was set to preferred for monoisotopic peak determination, charge states 2-6.

#### Raw MS/MS data processing

Raw MS/MS data from gastrocnemius, white adipose, lung, kidney, and cortex were processed using a cloud-based proteomics pipeline developed for this project and executed in the Google Cloud Platform. Briefly, we used the Workflow Description Language (WDL) to define the workflow and parallelize the execution using Cromwell as the workflow management system. The pipeline includes the following steps and open source methods:

1. MASIC: Extract reporter ion peaks from MS2 spectra and create Selected Ion Chromatograms for each MS/MS parent ion.
2. MSConverted: Convert Thermo .raw files to .mzML files.
3. MS-GF+: Identify peptides using a fully tryptic search (for speed).
4. MZrefiner filter in MSConvert: Use mass error histograms to in-silico re-calibrate the *m/z* values in the .mzML file.
5. PPMErrorCharter: Plot the mass error histograms before and after in-silico recalibration.
6. MS-GF+: Identify peptides using a partially tryptic search.
7. MzidToTSVConverter: Create a tab-separated value file listing peptide IDs.
8. PeptideHitResultsProcessor: Create tab-delimited files required for step 7; files contain peptide IDs, unique sequence info, and residue modification details.
9. Ascore (PTM only): Localize the position of Phosphorylation on S, T, and Y residues in phosphopeptides.

For a more extensive description, please visit the pipeline source code available here https://github.com/MoTrPAC/motrpac-proteomics-pipeline.

MS2 spectra were processed and searched against the rat RefSeq protein database (downloaded in November 2018) and common contaminants by the MS-GF+ tool[27] for peptide sequence identification. Fixed modifications were cysteine carbamidomethylation and TMT11 on N-terminal and lysine residues; variable modifications were methionine oxidation for global proteomics datasets, and S,T,Y phosphorylation for phosphoproteomics datasets. Localization of phosphorylation modifications was performed using the Ascore algorithm[28]. A target-decoy approach was used to control false discovery rate to <1%. TMT reporter ion intensities were extracted using MASIC[29] with the following thresholds: signal-to-noise ratio = 0; interference score = 0.5. Data from all fractions of the same multiplex were aggregated to peptide-centric, site-centric, or protein-centric levels, ratioed to the common reference, and summarized in results tables using the R tool "PlexedPiper" (https://github.com/PNNL-Comp-Mass-Spec/PlexedPiper).

Raw MS/MS data from heart and liver samples were processed using Spectrum Mill v.7.09.215 (Broad Institute). MS2 spectra were extracted from RAW files and merged if originating from the same precursor, or within a retention time window of +/- 60 s and *m/z* range of +/- 1.4, followed by filtering for precursor mass range of 750-6000 Da and sequence tag length > 0. MS/MS search was performed against the rat RefSeq protein database downloaded on November 2018 and common contaminants, with digestion enzyme conditions set to "Trypsin allow P", <5 missed cleavages, fixed modifications (cysteine carbamidomethylation and TMT11 on N-term and lysine), and variable modifications (oxidized methionine, acetylation of the protein N-terminus, pyroglutamic acid on N-term Q, and pyro carbamidomethyl on N-term C). Additional variable modifications were added for analysis of phosphoproteome (S,T, and Y phosphorylation), acetylome (K acetylation), and ubiquitylome (di-glycine residual in K). Matching criteria included a 30% minimum matched peak intensity and a precursor and product mass tolerance of +/- 20 ppm. Peptide-level matches were validated if found to be below the 1.0% false discovery rate (FDR) threshold and within a precursor charge range of 2-6. A second round of validation was then performed for protein-level matches for proteome datasets, requiring a minimum protein score of 13 and protein level FDR of 0%. Post-translational modification (PTM) site-centric tables for PTM datasets and protein-centric tables for proteome dataset, including TMT intensity values and ratio to the common reference, were exported using Spectrum Mill and used for further normalization and statistical analysis..

#### Proteomics data normalization

$Log_2$ TMT ratios to the common reference were used as quantitative values for all proteomics features (proteins, phosphosites, acetylsite, and ubiquitylsites). Proteomics datasets were examined for sample outliers by looking at the top principal components and by examining median protein abundance across samples. Outlier samples were identified for acetylome samples labeled with channel 130C, and these were suspected to originate from contaminated 130C-TMT reagent. All acetylome samples labeled with TMT channel 130C were excluded from downstream analysis. Proteomics features not fully quantified in at least two plexes within a tissue and non-rat contaminants were removed. $Log_2$ TMT ratios were sample-normalized by median-centering and mean absolute deviation scaling. Plex batch effects were removed using linear models implemented by the *limma::removeBatchEffect* function in R (v 3.48.0). The PTM datasets were corrected for changes in protein abundances by fitting a global linear model between the PTM-site and the cognate protein and extracting the residuals. Protein-corrected PTM values are available for exploration, but these were only used for the ubiquitylome differential analysis.

### Non-targeted metabolomics

#### HILIC LC-MS positive ion mode non-targeted metabolomics

Hydrophilic interaction liquid chromatography (HILIC) analyses of polar metabolites in the positive ionization mode were conducted at the Broad Institute of MIT and Harvard using an LC-MS system comprised of a Shimadzu Nexera X2 UHPLC (Shimadzu Corp., Kyoto, Japan) coupled to a Q-Exactive hybrid quadrupole Orbitrap mass spectrometer (Thermo Fisher Scientific). Powdered tissue samples (10 mg) were homogenized in 300 µL of 10/67.4/22.4/0.018 v/v/v/v water/acetonitrile/methanol/formic acid containing stable isotope-labeled internal standards (valine-d8, Sigma-Aldrich; and phenylalanine-d8, Cambridge Isotope Laboratories; Andover, MA) at 4°C using a TissueLyser II (QIAGEN) bead mill set to two 2 min intervals at 20 Hz. Plasma samples (10 µL) were samples were extracted using 90 µL of 74.9/24.9/0.2 v/v/v acetonitrile/methanol/formic acid containing valine-d8 and phenylalanine-d8 internal standards. Samples were centrifuged (10 minutes, 9,000 x g, 4°C), and the supernatants were injected directly onto a 150 x 2 mm, 3 µm Atlantis HILIC column (Waters). The column was eluted isocratically at a flow rate of 250 µL/min with 5% mobile phase A (10 mM ammonium formate and 0.1% formic acid in water) for 0.5 minute followed by a linear gradient to 40% mobile phase B (acetonitrile with 0.1% formic acid) over 10 minutes, then held at 40% B for 4.5 minutes. MS analyses were carried out using electrospray ionization in the positive ion mode using full scan analysis over 70-800 *m/z* at 70,000 resolution and 3 Hz data acquisition rate. Other MS settings were: sheath gas 40, auxiliary gas 10, sweep gas 2, spray voltage 3.5 kV, capillary temperature 350°C, S-lens RF 40, heater temperature 300°C, microscans 1, automatic gain control target 1e6, and maximum ion time 250 ms. Data quality was assured by i) initially confirming LC-MS system performance by analyzing a mixture of >140 well-characterized synthetic reference compounds as well as repeated analyses of extracts from human pooled plasma (BioIVT, Westbury, New York); ii) daily evaluation of internal standard signals to ensure that each sample injected properly and to monitor MS sensitivity; and iii) analysis of four pairs of pooled extract samples per sample type that were inserted in the analysis queue at regular intervals. One sample from each pair was used to correct for instrument drift using “nearest neighbor” scaling while the second reference sample served as a passive QC for determination of the analytical coefficient of variation of every identified metabolite and unknown. Raw data were processed using TraceFinder software (Thermo Fisher Scientific) for targeted peak integration and manual review of a subset of identified metabolites and Progenesis QI (Nonlinear Dynamics, Waters) for peak detection and integration of both metabolites of known identity and unknowns. Metabolite identities were confirmed using authentic reference standards.

#### Reverse phase and ion pairing LC-MS non-targeted metabolomics

Reverse-phase and ion pairing profiling of polar metabolites was conducted at the University of Michigan.

##### Materials and reagents

All solvents and mobile phase additives were LC-MS grade and purchased from Sigma-Aldrich. All chemical standards were the highest grade available and purchased from Sigma Aldrich or Cambridge Isotope Labs.

##### Sample preparation

**Plasma samples:** Samples were stored at -80°C until extraction and were thawed and maintained on wet ice throughout processing steps. Plasma (50 µl aliquot) was extracted by adding 200 µL of extraction solvent (1:1:1 v:v methanol:acetonitrile:acetone containing internal standards listed in Table 2). Samples were vortexed 10 seconds, incubated in ice for 10 minutes, then centrifuged for 10 minutes at 15,000 rcf at 4°C to pellet precipitated proteins. 150 µL supernatant was transferred to a glass autosampler vial with a low-volume insert and brought to dryness using a nitrogen blower at ambient temperature. Dried samples were reconstituted in 37.5 µL water:methanol (8:2 v:v) for LC-MS analysis. A QC sample was generated by pooling residual supernatant from multiple samples, then drying and reconstituting as above.

Table 2. Internal standard concentrations in extraction solvent

| Internal standard compound | Concentration in extraction solvent for plasma (µM, except as noted) | Concentration in extraction solvent for tissues (µM, except as noted) |
|---|---|---|
| $^{13}C_3$ lactic acid | 50 | 10 |
| $^{13}C_5$ alpha-ketoglutaric acid | 0.5 | 0.5 |
| $^{13}C_6$ citric acid | 5 | 0.5 |
| $^{13}C_4$ succinic acid | 0.5 | 0.5 |
| $^{13}C_4$ malic acid | 0.5 | 0.5 |
| $^{13}C_6$ fructose 6-phosphate | - | 1 |
| $^{13}C_6$ fructose 1,6-bisphosphate | - | 0.5 |
| $^{13}C_{10}{}^{15}N_5$ adenosine monophosphate | - | 0.5 |
| $^{13}C_{10}{}^{15}N_5$ adenosine triphosphate | - | 1 |

| | | |
|---|---|---|
| U-$^{13}$C amino acid mix (Sigma 426199) | 10 µg/ml | 5 µg/ml |
| $^{13}C_5$ glutamine | 25 | 5 |
| $^{13}C_6$ cystine | 5 | 1 |
| $^{15}N_2$ asparagine | 5 | 1 |
| $^{15}N_2$ tryptophan | 5 | 1 |
| $^{13}C_6$ glucose | 250 | 2 |
| $D_9$ L-carnitine | 0.38 | 0.38 |
| $D_4$ thymine | 1 | 1 |
| $^{15}N$ anthranilic acid | 1 | 1 |
| Gibberelic acid | 1 | 1 |
| Epibrassinolide | 1 | 1 |
| $D_3$ acetylcarnitine (Cambridge NSK-B) | 0.095 | 0.095 |
| $D_3$ propionylcarnitine (Cambridge NSK-B) | 0.019 | 0.019 |
| $D_3$ butyrylcarnitine (Cambridge NSK-B) | 0.019 | 0.019 |
| $D_9$ isovalerylcarnitine (Cambridge NSK-B) | 0.019 | 0.019 |
| $D_3$ octanoylcarnitine (Cambridge NSK-B) | 0.019 | 0.019 |
| $D_9$ myristoylcarnitine (Cambridge NSK-B) | 0.019 | 0.019 |
| $D_3$ palmitoylcarnitine (Cambridge NSK-B) | 0.038 | 0.038 |

**Tissue samples:** Frozen tissue samples were rapidly weighed into pre-tared, pre-chilled Eppendorf tubes and tissue mass was recorded to the nearest 0.1 mg. Extraction solvent was 1:1:1:1 methanol:acetonitrile:acetone:water containing internal standards. To extract samples, chilled extraction solvent was added to a tissue sample at the ratio of 1 ml solvent to 50 mg wet tissue mass. Immediately following solvent addition, the sample was homogenized using a Branson 450 probe sonicator set to output level 4, 40% duty cycle, for 30 seconds. Tubes were subsequently mixed several times by inversion and then incubated on ice for 10 minutes. Samples were centrifuged at 15,000 rcf for 10 minutes. 300 µL of supernatant was transferred to two autosampler vials with flat-bottom inserts, dried using the nitrogen blower, and stored at -80C until the day of analysis. Samples were reconstituted in 60 µL of 8:2 water:methanol and submitted for LC-MS analysis. A QC sample was generated by pooling residual supernatant from multiple samples, then drying and reconstituting as above.

#### LC-MS analysis

**Non-targeted reverse phase LC-MS:** Samples were analyzed on an Agilent 1290 Infinity II / 6545 qTOF MS system with a JetStream electrospray ionization (ESI) source (Agilent Technologies, Santa Clara, California) using a Waters Acquity HSS T3 column, 1.8 µm 2.1 x 100 mm equipped with a matched Vanguard precolumn (Waters Corporation). Mobile phase A was 100% water with 0.1% formic acid and mobile phase B was 100% methanol with 0.025% formic acid. The gradient was as follows: Linear ramp from 0% to 100% B from 0-10 minutes, hold 100% B until 17 minutes, linear return to 0% B from 17 to 17.1 minutes, hold 0% B until 20 minutes. The flow rate was 0.45 ml/min, the column temperature was 55°C, and the injection volume was 5 µL. Each sample was analyzed twice, once in positive and once in negative ion mode MS, scan rate 2 spectra/sec, mass range 50-1200 *m/z*. Source parameters were: drying gas temperature 350°C, drying gas flow rate 10 L/min, nebulizer pressure 30 psig, sheath gas temperature 350°C and flow 11 L/minute, capillary voltage 3500 V, internal reference mass correction enabled. A QC sample run was performed at minimum every tenth injection.

**Non-targeted ion pairing LC-MS:** Samples were analyzed on an identically-configured LC-MS system using an Agilent Zorbax Extend C18 1.8 µm RRHD column, 2.1 x 150 mm ID, equipped with a matched guard column. Mobile phase A was 97% water, 3% methanol. Mobile phase B was 100% methanol. Both mobile phases contained 15 mM tributylamine and 10 mM acetic acid. Mobile phase C was 100% acetonitrile. Elution was carried out using a linear gradient followed by a multi-step column wash including automated (valve-controlled) backflushing (see Table 3). Column temperature was 35°C and the injection volume was 5 µL. MS acquisition was performed in negative ion mode, scan rate 2 spectra/sec, mass range 50-1200 *m/z*. Source parameters were: drying gas temperature 250°C, drying gas flow rate 13 L/min, nebulizer pressure 35 psig, sheath gas temp 325°C and flow 12 L/min, capillary voltage 3500V, internal reference mass correction enabled. A QC sample run was performed at minimum every tenth injection.

Table 3. IPC-MS gradient program

| Time (minutes) | %A | %B | %C | Flow (ml/min) | Flow direction |
|---|---|---|---|---|---|
| 0 | 100 | 0 | 0 | 0.25 | Normal |
| 2 | 100 | 0 | 0 | 0.25 | Normal |
| 12 | 1 | 99 | 0 | 0.25 | Normal |
| 18 | 1 | 99 | 0 | 0.25 | Normal |
| 18.05 | 5 | 0 | 95 | 0.25 | Backflush |
| 21 | 5 | 0 | 95 | 0.25 | Backflush |
| 21.50 | 5 | 0 | 95 | 0.80 | Backflush |
| 23 | 5 | 0 | 95 | 0.80 | Backflush |

| | | | | | |
|---|---|---|---|---|---|
| 23.2 | 5 | 0 | 95 | 0.60 | Backflush |
| 24.00 | 100 | 0 | 0 | 0.40 | Backflush |
| 27.99 | 100 | 0 | 0 | 0.40 | Backflush |
| 28.0 | 100 | 0 | 0 | 0.40 | Normal |
| 29.9 | 100 | 0 | 0 | 0.40 | Normal |
| 30 | 100 | 0 | 0 | 0.25 | Normal |

**Iterative Data Dependent MS/MS data acquisition (iDDA):** To aid in compound identification, iterative MS/MS data was acquired for both reverse phase and ion pairing methods using the pooled sample material. Eight repeated LC-MS/MS runs of the QC sample were performed at three different collision energies (10, 20, and 40) with iterative acquisition enabled. The software excluded precursor ions from MS/MS acquisition within 0.5 minute of their MS/MS acquisition time in prior runs, resulting in deeper MS/MS coverage of lower-abundance precursor ions.

#### Data analysis

**Feature detection and alignment:** Data analysis was performed using a hybrid targeted/untargeted approach. Targeted compound detection and relative quantitation was performed by automatic integration followed by manual review and correction using Profinder v8.0 (Agilent Technologies, Santa Clara, CA.) Non-targeted feature detection was performed using custom scripts that automate operation of the “find by molecular feature” workflow of the Agilent Masshunter Qualitative Analysis (v7) software package. Feature alignment and recursive feature detection were performed using Agilent Mass Profiler Pro (v8.0) and Masshunter Qualitative Analysis (“find by formula” workflow), yielding an aligned table including *m/z*, RT, and peak areas for all features.

**Data Cleaning and Degeneracy Removal:** A combined feature set was generated by merging untargeted features and named metabolites into a single feature list. Features missing from over 50% of all samples in a batch or over 30% of QC samples were removed prior to subsequent normalization steps. Next, the combined feature set underwent data reduction using Binner[30]. Briefly, Binner first performs RT-based binning, followed by clustering of features by Pearson’s correlation coefficient, and then assigns annotations for isotopes, adducts or in-source fragments by searching for known mass differences between highly correlated features.

**Normalization and Quality Control:** Data were normalized using a Systematic Error Removal Using Random Forest (SERRF) approach[31], which helps correct for drift in peak intensity over the batch using data from the QC sample runs. When necessary to correct for residual drift, peak area normalization to closest-matching internal standard was also applied to selected compounds. Both SERRF correction and internal standard normalization were implemented in R. Parameters were set to minimize batch effects and other observable drift, as visualized using principal component analysis score plots of the full dataset. Normalization performance was also validated by examining relative standard deviation values for additional QC samples not included in the drift correction calculations. Quality control reports containing these data were generated for all datasets and uploaded to the MoTrPAC data repository along with raw and processed data.

**Compound identification:** Metabolites from the targeted analysis workflow were identified with high confidence (MSI level 1)[32] by matching retention time (+/- 0.1 minute), mass (+/- 10 ppm) and isotope profile (peak height and spacing) to authentic standards. MS/MS data corresponding to unidentified features of interest from the untargeted analysis were searched against a spectral library (NIST 2020 MS/MS spectral database or other public spectral databases) to generate putative identifications (MSI level 2) or compound-class level annotations (MSI level 3) as described previously[33].

#### Non-targeted LC-MS/MS lipidomics

##### Sample preparation

Non-targeted lipid analysis was conducted at the Georgia Institute of Technology. Powdered tissue samples (10 mg) were extracted in 400 µl isopropanol containing stable isotope-labeled internal standards (IS) by freeze-thawing in liquid nitrogen followed by sonication in an ice bath, repeated three times. Samples were then centrifuged (5 min, 21,100xg), and the supernatants transferred to autosampler vials. Plasma samples (25 µL) were extracted by mixing with 75 µL isopropanol containing the IS mix followed by centrifugation. Sample blanks, pooled extract samples used as quality controls (QC), and consortium reference samples, were prepared for analysis using the same methods. The IS mix consisted of PC (15:0-18:1(d7)), Catalog No. 791637; PE (15:0-18:1(d7)), Catalog No. 791638; PS (15:0-18:1(d7)), Catalog No. 791639; PG(15:0-18:1(d7)), Catalog No. 791640; PI(15:0-18:1(d7)), Catalog No. 791641; LPC(18:1(d7)), Catalog No. 791643; LPE(18:1(d7)); Catalog No. 791644; Chol Ester (18:1(d7)), Catalog No. 700185; DG(15:0-18:1(d7)), Catalog No. 791647; TG(15:0-18:1(d7)-15:0), Catalog No. 791648; SM(18:1(d9)), Catalog No. 791649; Cholesterol (d7), Catalog No. 700041. All internal standards were purchased from Avanti Polar Lipids (Alabaster, Alabama) and added to the extraction solvent at a final concentration in the 0.1-8 µg/ml range.

##### Data collection

Lipid LC-MS data were acquired using a Vanquish (ThermoFisher Scientific) chromatograph fitted with a ThermoFisher Scientific AccucoreTM C30 column (2.1 × 150 mm, 2.6 µm particle size), coupled to a high-resolution accurate mass Q-Exactive HF Orbitrap mass spectrometer (ThermoFisher Scientific) for both positive and negative ionization modes. The mobile phases were 40:60 water:acetonitrile with 10 mM ammonium formate and 0.1% formic acid (mobile phase A), and 10:90 acetonitrile:isopropyl alcohol, with 10 mM ammonium formate and 0.1% formic acid (mobile phase B). The chromatographic method used the following gradient program: 0 minutes 80% A; 1 minute 40% A; 5 minutes 30% A; 5.5 minutes 15% A; 8 minutes 10% A; held 8.2 minutes to 10.5 minutes 0% A; 10.7 minutes 80% A; and held until 12 minutes. The flow rate was set at 0.40 ml/min. The column temperature was set to 50°C, and the injection volume was 2 µL.

For analysis of the organic phase the electrospray ionization source was operated at a vaporizer temperature of 425°C, a spray voltage of 3.0 kV for positive ionization mode and 2.8 kV for negative ionization mode, sheath, auxiliary, and sweep gas flows of 60, 18, and 4 (arbitrary units), respectively, and capillary temperature of 275°C. The instrument acquired full MS data with 240,000 resolution over the 150-2000 *m/z* range. LC-MS/MS experiments were acquired using a DDA strategy. MS2 spectra were collected with a resolution of 120,000 and the dd-MS2 were collected at a resolution of 30,000 and an isolation window of 0.4 *m/z* with a loop count of top 7. Stepped normalized collision energies of 10%, 30%, and 50% fragmented selected precursors in the collision cell. Dynamic exclusion was set at 7 seconds and ions with charges greater than 2 were omitted.

#### Data processing

Data processing steps included peak detection, spectral alignment, grouping of isotopic peaks and adduct ions, drift correction, and gap filling. Compound Discoverer V3.0 (ThermoFisher Scientific) was used to process the raw LC-MS data. Drift correction was performed on each individual feature, where a linear curve was fitted to the pooled QC sample peak areas across the batch and was then used to correct the peak area for that specific feature in the samples. Detected features were filtered with background and QC filters. Features with abundance lower than 5x the background signal in the sample blanks and that were not present in at least 50% of the QC pooled injections with a coefficient of variance (CV) lower than 30% were removed from the dataset. Lipid annotations were accomplished based on accurate mass and relative isotopic abundances (to assign elemental formula), retention time (to assign lipid class), and MS2 fragmentation pattern matching to local spectral databases built from curated experimental data. Lipid nomenclature followed that described by Fahy et al.[34,35].

#### Quality control procedures

System suitability was assessed prior to the analysis of each batch. A performance baseline for a clean instrument was established before any experiments were conducted. The mass spectrometers were mass calibrated, mass accuracy and mass resolution were checked to be within manufacturer specifications, and signal-to-noise ratios for the suite of IS checked to be at least 75% of the clean baseline values. For LC-MS assays, an IS mix consisting of 12 standards was injected to establish baseline separation parameters for each new column. The performance of the LC gradient was assessed by inspection of the column back pressure trace, which had to be stable within an acceptable range (less than 30% change). Each IS mix component was visually evaluated for chromatographic peak shape, retention time (lower than 0.2 minute drift from baseline values) and FWHM lower than 125% of the baseline measurements. The CV of the average signal intensity and CV of the IS (<=15%) in pooled samples were also checked. These pooled QC samples were used to correct for instrument sensitivity drift over the various batches using a procedure similar to that described by the Human Serum Metabolome (HUSERMET) Consortium[36]. To evaluate the quality of the data for the samples themselves, the IS signals across the batch were monitored, PCA modeling for all samples and features before and after drift correction was conducted, and Pearson correlations calculated between each sample and the median of the QC samples.

### Targeted metabolomics

#### LC-MS/MS analysis of branched-chain keto acids

Targeted profiling of branched-chain keto acid metabolites was conducted at Duke University. Ten µl of plasma containing isotopically labeled ketoleucine (KIC)-d3, ketoisovalerate (KIV)-$^{13}C_5$ (Cambridge Isotope Laboratories), and 3-methyl-2-oxovalerate (KMV)-d8 (Toronto Research Chemicals, Canada) internal standards were deproteinated with 150 µl of 3M perchloric acid. Two hundred µl of tissue homogenate prepared at 100 mg/ml in 3M perchloric acid were centrifuged at 14,000 x g for 5 minutes. Two hundred µl of 25 M o-phenylenediamine (OPD) in 3M HCl were added to the plasma and tissue supernatants, and the samples were incubated at 80°C for 20 minutes. Keto acids were extracted with ethyl acetate as previously described[37,38]. The extracts were dried under nitrogen, reconstituted in 200 mM ammonium acetate, and analyzed on a Xevo TQ-S triple quadrupole mass spectrometer coupled to an Acquity UPLC (Waters) controlled by the MassLynx 4.1 operating system. The analytical column (Waters Acquity UPLC BEH C18 Column, 1.7 µm, 2.1 × 50 mm) was used at 30°C. 10 µl of the sample were injected onto the column and eluted at a flow rate of 0.4 ml/min. The gradient consisted of 45% mobile phase A (5 mM ammonium acetate in water) and 55% mobile phase B (methanol) for 2 minutes, followed by a linear gradient to 95% B from 2 to 2.5 minutes, held at 95% B for 0.7 minutes, returned to 45% A, and finally the column was re-equilibrated at initial conditions for 1 minute. The total run time was 4.7 minutes. Mass transitions of *m/z* 203 → 161 (KIC), 206 → 161 (KIC-d3), 189 → 174 (KIV), 194 → 178 (KIV-$^{5}C_{13}$), 203 → 174 (KMV), and 211 → 177 (KMV-d8) were monitored in positive ion mode. The endogenous keto acids were quantified using calibrators prepared by spiking dialyzed fetal bovine serum with authentic keto acids (Sigma-Aldrich).

#### Flow injection MS/MS analysis of acyl CoAs

Targeted profiling of acyl CoAs was conducted at Duke University. 500 µl of tissue homogenate prepared at 50 mg/ml in isopropanol/0.1 M $KH_2PO_4$ (1:1) was extracted with an equal volume of acetonitrile and centrifuged at 14,000 x g for 10 min as previously described[39,40]. The supernatants were acidified with 0.25 ml of glacial acetic acid, and acyl CoAs were further purified by solid phase extraction (SPE) using 2-(2-pyridyl) ethyl functionalized silica gel (Sigma-Aldrich) as described[41]. The SPE columns were conditioned with 1 ml of acetonitrile/isopropanol/water/glacial acetic acid (9/3/4/4 : v/v/v/v). Following application and flow through of the supernatant, the SPE columns were washed with 2 ml of acetonitrile/ isopropanol/water/glacial acetic acid (9/3/4/4 : v/v/v/v). Acyl CoAs were then eluted with 2 ml of methanol/250 mM ammonium formate (4/1 : v/v) and analyzed by flow injection MS/MS analysis using positive ion mode on a Xevo TQ-S, triple quadrupole mass spectrometer (Waters), employing methanol/water (80/20, v/v) containing 30 mM ammonium hydroxide as the mobile phase. Spectra were acquired in the multichannel acquisition mode monitoring the neutral loss of 507 amu (phosphoadenosine diphosphate) and scanning from *m/z* 750 to 1100.

Heptadecanoyl CoA was employed as an internal standard. The endogenous Acyl CoAs were quantified using calibrators prepared by spiking tissue homogenates with authentic Acyl CoAs (Sigma-Aldrich) having saturated acyl chain lengths C0 - C18. Corrections for the heavy isotope effects, mainly $^{13}C$, to the adjacent m+2 spectral peaks in a particular chain length cluster were made empirically by referring to the observed spectra for the analytical standards.

#### LC-MS/MS analysis of nucleotides

Targeted profiling of nucleotide metabolites was conducted at Duke University. 300 µl of tissue homogenates prepared at 50 mg/ml in 70% methanol were spiked with nine internal standards: $^{13}C^{10}$,$^{15}N^{5}$-adenosine monophosphate, $^{13}C^{10}$,$^{15}N^{5}$-guanosine monophosphate, $^{13}C^{10}$,$^{15}N^{2}$-uridine monophosphate, $^{13}C^{9}$,$^{15}N^{3}$-cytidine monophosphate, $^{13}C^{10}$-guanosine triphosphate, $^{13}C^{10}$-uridine triphosphate, $^{13}C^{9}$-cytidine triphosphate, $^{13}C^{10}$-adenosine triphosphate, and nicotinamide-1, $N^{6}$-ethenoadenine dinucleotide (eNAD) (Sigma-Aldrich). Nucleotides were extracted using an equal volume of hexane as described by Cordell et al. and Gooding et al.[42,43]. The samples were vortexed and centrifuged at 14,000 x g for 5 minutes. The bottom layer was collected and centrifuged again. Chromatographic separations and MS analysis of the supernatants were performed using an Acquity UPLC system (Waters) coupled to a Xevo TQ-XS quadrupole mass spectrometer and (Waters). The analytical column (Chromolith FastGradient RP-18e 50-2mm column, EMD Millipore, Billerica, MA, USA) was maintained at 40°C. The injection volume was 2 µL. Nucleotides were separated using a mobile phase A consisting of 95% water, 5% methanol, and 5 mM dimethylhexylamine adjusted to pH 7.5 with acetic acid and a mobile phase B consisting of 20% water, 80% methanol, and 10 mM dimethylhexylamine. Flow rate was set to 0.3 ml/min. The 22-minute gradient (t=0, %B=0; t=1.2, %B=0; t=22, %B=40) was followed by a 3-minute wash and 7-minute equilibration. Nucleotides were detected in the negative ion MRM mode based on characteristic fragmentation reactions. The endogenous nucleotides were quantified using calibrators prepared by spiking tissue homogenates with authentic nucleotides (Sigma-Aldrich).

#### LC-MS/MS analysis of amino acids and amino metabolites

Targeted profiling of amino acids and amino metabolites was conducted at the Mayo Clinic by LC-MS as previously described[44,45]. Briefly, either 20 ml of plasma samples or 5 mg of tissue homogenates were spiked with an internal standard solution consisting of isotopically labeled amino acids (U-$^{13}C_4$ L-aspartic acid,U-$^{13}C_3$ alanine, U-$^{13}C_4$ L-threonine, U-$^{13}C$ L-proline, U-$^{13}C_6$ tyrosine, U-$^{13}C_5$ valine, U-$^{13}C_6$ leucine, U-$^{13}C_6$ phenylalanine, U-$^{13}C_3$ serine, U-$^{13}C_5$ glutamine, U-$^{13}C_2$ glycine, U-$^{13}C_5$ glutamate, U-$^{13}C_6$,$^{15}N_2$ lysine, U-$^{13}C_5$,$^{15}N$ methionine, 1,1U-$^{13}C_2$ homocysteine, U-$^{13}C_6$ arginine, U-$^{13}C_5$ ornithine, $^{13}C_4$ asparagine, $^{13}C_2$ ethanolamine, d3 sarcosine, d6 4-aminobutyric acid) . The supernatant was immediately derivatized with 6-aminoquinolyl-N-hydroxysuccinimidyl carbamate using a MassTrak kit (Waters). A 10-point calibration standard curve underwent a similar derivatization procedure after the addition of internal standards. Both derivatized standards and samples were analyzed on a Quantum Ultra triple quadrupole mass spectrometer (ThermoFischer) coupled with an Acquity liquid chromatography system (Waters). Data acquisition was conducted by using selected ion monitoring (SRM) in positive ion mode. Concentrations of 42 analytes of each unknown were calculated against their respective calibration curves.

#### GC-MS analysis of tricarboxylic acid cycle (TCA) metabolites

Targeted profiling of TCA metabolites was conducted at the Mayo Clinic by gas chromatography mass spectrometry (GC-MS) as previously described[46,47], with a few modifications. Briefly, 5 mg of tissue were homogenized in 1X PBS on an Omni bead ruptor (Omni International, Kennesaw, GA) prior to adding 20 µl of internal solution containing U-$^{13}C$ labeled analytes ($^{13}C_3$ sodium lactate, $^{13}C_4$ succinic acid, $^{13}C_4$ fumaric acid, $^{13}C_4$ alpha ketoglutaric acid, $^{13}C_4$ malic acid, $^{13}C_4$ aspartic acid, $^{13}C_5$ 2-hydroxyglutaric acid, $^{13}C_5$ glutamic acid, $^{13}C_6$ citric acid, $^{13}C_2$,$^{15}N$ glycine, $^{13}C_2$ sodium pyruvate). For plasma, 50 µl were used. The proteins were removed by adding 300 µl of chilled methanol and acetonitrile solution to the sample mixture. After drying the supernatant in the speedvac, the sample was derivatized with ethoxime and then with MtBSTFA + 1% tBDMCS (N-Methyl-N-(t-Butyldimethylsilyl)-Trifluoroacetamide + 1% t-Butyldimethylchlorosilane) before it was analyzed on an Agilent 5977B GC/MS (Santa Clara, California) under single ion monitoring conditions using electron ionization. Concentrations of lactic acid (*m/z* 261.2), fumaric acid (*m/z* 287.1), succinic acid (*m/z* 289.1), ketoglutaric acid (*m/z* 360.2), malic acid (*m/z* 419.3), aspartic acid (*m/z* 418.2), 2-hydroxyglutaratic acid (*m/z* 433.2), cis aconitic acid (*m/z* 459.3), citric acid (*m/z* 591.4), and isocitric acid (*m/z* 591.4), glutamic acid (*m/z* 432.4) were measured against 7-point calibration curves that underwent the same derivatization procedure.

#### LC-MS/MS analysis of ceramides

Targeted profiling of ceramides was conducted at the Mayo Clinic. Plasma and tissue ceramides, sphinganine, sphingosine, sphingosine-1-phosphate (S1P) were measured by previously described methods[48,49]. Briefly, a 25 µl aliquot of plasma or 5 mg tissue homogenate was spiked with an internal standard mixture prior to undergoing extraction. Data acquisition was conducted in SRM mode after chromatographic separation in an electron impact Thermo Quantiva mass spectrometer coupled to a Waters Acquity UPLC system. Concentrations of each analyte were calculated against their respective calibration curves. Coefficients of variation for a healthy plasma control analyzed with each batch of 40 samples over a one-month period were 6.3%, 6.2%, 3.1%, 5.0%, 5.7%, 3.2%, 4.9% and 3.3% for sphingosine, sphingosine-1-phosphate, C16:0-ceramides, C18:0-ceramides, C20:0-ceramide, C22:0-ceramide, C24:1-ceramide and C24:0-ceramide, respectively.

#### LC-MS/MS analysis of acylcarnitines

Targeted profiling of acylcarnitines was conducted at the Mayo Clinic. Acylcarnitines (specifically C0-C18:1) were measured by LC-MS using established methods[47,50]. Briefly, 25 µl of plasma or 5 mg homogenized tissue were spiked with an IS consisting of isotopically-labeled acylcarnitines. The samples were then extracted with cold MeOH:DCM (1:1) followed by centrifugation at 12,000 g for 10 minutes. The supernatant was transferred to another vial, dried down and reconstituted in mobile phase. A calibration curve was built from a purchased acyl carnitine mix aliquoted at various concentrations, and spiked with the same IS as the samples. The samples and calibration standards were analyzed on a triple quadrupole mass spectrometer coupled with an UHPLC system. Data acquisition was conducted in SRM mode. Analyte concentrations of each unknown were calculated against their perspective standard curves.

#### Targeted lipidomics of low-level lipids

##### Sample preparation

Targeted profiling of lipids was conducted at Emory University using previously published methods[51,52]. Powdered tissue samples (10 mg) were homogenized in 100 µl PBS with Bead Ruptor (Omni International, Kennesaw, GA). Homogenized samples were diluted with 300 µL 20% methanol and spiked with 1% BHT solution to a final BHT concentration of 0.1% and pH of 3.0 by acetic acid addition. Samples were then centrifuged (10 minutes, 14000 rmp), and the supernatants were transferred to 96-well plates for further extraction. The supernatants were loaded to Isolute C18 SPE columns (conditioned with 1000 µL ethyl acetate and 1000 µL 5% methanol). The SPE columns were then washed with 800 µL water and 800 µL hexane. The oxylipins were eluted with 400 µl methyl formate. SPE was conducted automatically with a Biotage Extrahera (Uppsala, Sweden). The eluate was dried with nitrogen and then reconstituted with 200 µL methanol prior to LC-MS analysis. Sample blanks, pooled extract samples used as quality controls (QC), and consortium reference samples, were prepared for analysis using the same methods. The external standards consisted of prostaglandin E2 ethanolamide (Catalog No. 100007212), oleoyl ethanolamide (Catalog No. 90265), palmitoyl ethanolamide (Catalog No. 10965), arachidonoyl ethanolamide (Catalog No. 1007270), docosahexaenoyl ethanolamide (Catalog No. 10007534), linoleoyl ethanolamide (Catalog No. 90155), stearoyl ethanolamide (Catalog No. 90245), oxy-arachidonoyl ethanolamide (Catalog No. 10008642), 2-arachidonyl glycerol (Catalog No. 62160), docosatetraenoyl ethanolamide (Catalog No. 90215), α-linolenoyl ethanolamide (Catalog No. 902150), oleamide (Catalog No. 90375), dihomo-γ-linolenoyl ethanolamide (Catalog No. 09235), decosanoyl ethanolamide (Catalog No. 10005823), 9,10 DiHOME (Catalog No. 53400), prostaglandin E2-1-glyceryl ester (Catalog No. 14010), 20-HETE (Catalog No. 10007269), 9-HETE (Catalog No. 34400), 14,15 DiHET (Catalog No. 10007267), 5(S)-HETE (Catalog No. 34210), 12(R)-HETE (Catalog No. 10007247), 11(12)-DiHET (Catalog No. 10007266), 5,6-DiHET (Catalog No. 10007264), thromboxane B2 (Catalog No. 10007237), 12(13)-EpOME (Catalog No. 52450), 13 HODE (Catalog No. 38600), prostaglandin F2α (Catalog No. 10007221), 14(15)-EET (Catalog No. 10007263), 8(9)-EET (Catalog No. 10007261), 11(12)-EET (Catalog No. 10007262), leukotriene B4 (Catalog No. 20110), 8(9)-DiHET (Catalog No. 10007265), 13-OxoODE (Catalog No. 38620), 13(S)-HpODE (Catalog No. 48610), 9(S)-HpODE (Catalog No. 48410), 9(S)-HODE (Catalog No. 38410), resolvin D3 (Catalog No. 13834), resolvin E1 (Catalog No. 10007848), resolvin D1 (Catalog No. 10012554), resolvin D2 (Catalog No. 10007279), 9(S)HOTrE (Catalog No. 39420), 13(S)HOTrE (Catalog No. 39620), 8-iso Progstaglandin F2α (Catalog No. 25903). All external standards were purchased from Cayman Chemical (Ann Arbor, Michigan) at a final concentration in the range 0.01-20 µg/ml.

##### Data collection

LC-MS data were acquired using an ExionLC (SCIEX, Waltham, MA) chromatograph fitted with a ThermoFisher Scientific AccucoreTM C18 column (100 mm × 4.6, 2.6 µm particle size), coupled to a SCIEX QTRAP 5500 mass spectrometer for both positive and negative ion modes. The mobile phases were water with 10 mM ammonium acetate (mobile phase A), and acetonitrile with 10 mM ammonium acetate (mobile phase B). The chromatographic method used the following gradient program: 0.5 minute with 10% A; 1 minute to 2 minutes with 50% A; 2.1 minutes to 5.0 minutes with 75% A; 7 minutes to 13 minutes with 85% A; and 14.1 minutes until 18 minutes with 10% A. The flow rate was set at 0.50 ml/min. The column temperature was set to 50°C, and the injection volume was 10 µL with negative ion mode and 2 µL with positive ion mode. For mass spectrometry analysis, the heated electrospray ionization source was operated at a vaporizer temperature of 650°C, a spray voltage of 5.5 kV for positive ion mode and 4.5 kV for negative ion mode, curtain gas, ion source gas 1, ion source gas 2 were 20, 60 and 50, respectively. The decluttering potential, entrance potential, collision energy, and collision cell exit potential were 200, 10, 40, and 10 for the negative ion mode, respectively and 90, 10, 47 and 18, respectively, for positive ion mode.

##### Data processing and quality control

Sciex OS (AB SCIEX, Version 1.6.1) was used to process the raw LC-MS data. Standard curves were built for each oxlipin/ethanolamide and checked, i.e, all concentration points should be in the linear portion of the curve with an R-squared value no less than 0.9. Additionally, features with a high coefficient of variation (CV) among the QCs were removed from the dataset. Pearson correlation among the QCs for each tissue type were calculated with the *Hmisc* R library, and figures documented in QC report were plotted with the *corrplot* R library[53,54].

#### Metabolomics data processing and normalization

Metabolomics sample pre-processing was performed by each Chemical Analysis Site (CAS) as described above for each platform. These datasets were separated into two data types: “named”, for chemical compounds confidently identified, and “unnamed”, for compounds with specific chemical properties but without a standard chemical name. Only named metabolites were included in downstream analyses. The resulting named datasets were the input for the data processing and normalization procedures described in this section. Table 4 provides a summary of features measured by each platform, and the tissues and samples analyzed by each platform are presented in Figure S1E. The large number of features identified by reversed-phase positive untargeted lipidomics (lrppos) is due to a large number of multi-peak isomers.

For both untargeted and targeted datasets, measurements were $log_2$-transformed, negative values were converted to missing values, and features with >20% missing values were removed. For untargeted datasets and targeted datasets with more than 12 features, remaining missing values were imputed using K-nearest neighbors (k=10 samples). Missing values in targeted datasets with 12 or fewer features were not imputed. For outlier identification in each dataset, we calculated each sample's median correlation value against the other N-1 samples and selected a threshold to designate outliers as those with below-threshold median correlation values. All outliers were reviewed by Metabolomics CAS, and only 21 confirmed technical outliers in the untargeted datasets were removed.

To normalize untargeted datasets, we median-centered samples if neither sample medians nor upper quartiles were significantly associated with sex or sex-stratified training group (Kruskal-Wallis p-value < 0.01). Targeted datasets were not normalized as they were quantified using absolute concentrations.

A total of 1116 metabolites were measured by two or more platforms. We retained all features with repeated measurements and integrated their differential analysis results using meta-regression (see “Meta-analysis of metabolomics results”).

Table 4. Description of metabolomics platforms.

| ASSAY | DESCRIPTION | SITE | TARGETED | N TISSUES | MAX N NAMED FEATURES |
|---|---|---|---|---|---|
| hilicpos | HILIC positive | Broad | 0 | 19 | 306 |
| lrppos | Untargeted lipidomics, reversed-phase positive | GTech | 0 | 9 | 1341 |
| rppos | Reversed-phase positive | UMich | 0 | 9 | 211 |
| ionpneg | Ion pairing negative | UMich | 0 | 9 | 80 |
| rpneg | Reversed-phase negative | UMich | 0 | 9 | 160 |
| lrpneg | Untargeted lipidomics, reversed-phase negative | GTech | 0 | 9 | 523 |
| amines | Amines | Mayo | 1 | 9 | 39 |
| tca | TCA cycle | Mayo | 1 | 9 | 12 |
| etamidpos | Ethanolamides | Emory | 1 | 9 | 10 |
| oxylipneg | Oxylipins | Emory | 1 | 9 | 27 |
| ka | Ketoacids | Duke | 1 | 5 | 3 |
| acoa | Acyl-CoA | Duke | 1 | 4 | 67 |
| nuc | Nucleotides | Duke | 1 | 4 | 36 |

### Multiplexed bead-based immunoassays

#### Protein extraction

Tissue lysates for the immunoassays were prepared at Stanford University. All 30 samples per tissue (15+15 for testes and ovaries, 3 samples per sex per time point, 5 time points) were processed in a single batch. Upon retrieval from storage at -80°C, tissue samples were kept on dry ice until addition of a lysis buffer. Cold lysis buffer (25mM Tris-HCl, pH 7.5; 100mM NaCl; 0.01% Triton-X 100; 0.0025% NP-40) was supplemented with Halt protease inhibitor cocktail (Thermo Fisher # 78429) to 1x just prior to use. 550 µl of chilled lysis buffer were added to each tissue sample; the tube was inverted several times and placed on ice. To facilitate lysis, each sample was sonicated using a Branson Sonifier 250 with 3 1-second bursts at an output setting of 1 and constant duty cycle. Samples were rocked at 4°C for 1 hour. Sample tubes were then centrifuged at max speed (>20,000 xg) at 4°C for 10 minutes to pellet debris. The cleared supernatant from each sample was then transferred into an Eppendorf Protein Lo-Bind 1.5-ml tube. Protein concentrations of lysates were measured in duplicates using the BCA Assay (Thermo Fisher # 23225). A Tecan Infinite M1000 plate reader was used to read absorbance at 562 nm. Microsoft Excel was used to generate a standard curve based on the background-subtracted absorbance of provided BSA protein standards, and this curve was used to extrapolate the protein concentrations in each tissue sample lysate. All samples for a given tissue were then normalized to the protein concentration of the lowest sample through addition of an appropriate amount of lysis buffer. Lysates from each tissue were then plated into 2 replicate 96-well master plates (Greiner # 651201). Plates were sealed with PCR seals and stored at -80°C prior to analysis.

#### Targeted multiplexed bead-based immunoassays

The immunoassays were performed by the Stanford Human Immune Monitoring Center (HIMC). The rat tissue lysates were kept at -80°C until used for immunoassays. Levels of 54 cytokines were measured in rat samples using five Luminex panels: MILLIPLEX MAP Rat Cytokine/Chemokine Magnetic Bead Panel (Millipore, RECYTMAG-65K); MILLIPLEX MAP Rat Myokine Magnetic Bead Panel (Millipore, RMYOMAG-88K); MILLIPLEX MAP Rat Metabolic Hormone Magnetic Bead Panel (Millipore, RMHMAG-84K); MILLIPLEX MAP Rat Putuitary Magnetic Bead Panel (Millipore, RPTMAG-86K); MILLIPLEX MAP Rat Adipokine Magnetic Bead Panel (Millipore, RADPKMAG-80K). All samples from the same tissue were analyzed on the same plate to avoid plate-to-plate variability. Custom Assay CHEX control beads (Radix BioSolutions, Georgetown, Texas) were added to all wells to monitor instrument performance, application of the detection antibody, application of the fluorescent reporter, and nonspecific binding (CHEX1, CHEX2, CHEX3, and CHEX4, respectively) [55]. Samples were mixed with antibody-linked magnetic beads on a 96-well plate and incubated overnight at 4°C with shaking. Cold and room-temperature incubation steps were performed on an orbital shaker at 500-600 rpm. Plates were washed twice with wash buffer in a Biotek ELx405 washer. Following one-hour incubation at room temperature with biotinylated detection antibody, streptavidin-PE was added for 30 minutes with shaking. Plates were washed as described above and PBS was added to wells for reading in the Luminex FlexMap3D Instrument with a lower bound of 50 beads per sample per cytokine. Reports were generated using Masterplex QT/Mirai Bio/Hitachi software and exported as csv files. Samples were measured in singlets.

#### Data preprocessing

Raw mean fluorescence intensities (MFI) were $log_2$-transformed, and measurements for analytes with fewer than 20 beads in a well were removed. $Log_2$-MFI for analytes measured by multiple panels were correlated to determine reproducibility. TNFA measurements were close to background levels in the Rat Metabolic Hormone Panel likely due to dilutions, while TNFA measured in Rat Cytokine/Chemokine Magnetic Bead Panel showed a dynamic range of signal intensities. Thus, TNFA measurements from the Rat Metabolic Hormone Panel were removed. For remaining measurements in each panel and tissue, samples with more than 50% missing values (i.e., due to low bead count) were removed; features with at least two missing values for a single experimental group (e.g., males trained for 2 weeks) were removed (this affected four colon analytes and one spleen analyte, all in the Rat Cytokine/Chemokine Panel). Remaining missing values were imputed with k-nearest neighbors (k=5 features). This was the version of the data used for visualization (e.g., principal component analysis). Within each panel, tissue, and analyte, we calculated the mean and standard deviation and removed outlying measurements more than 4 standard deviations away from the mean. This version of the data with extreme feature-specific outliers removed was used for differential analysis.

### STATISTICAL ANALYSES

#### Outlier identification

Outlier identification for metabolomics datasets is described in “Metabolomics data processing and normalization.” Otherwise, outliers were identified through a combination of principal component analysis of normalized datasets and examination of assay-specific quality control metric scores. For all proteomics, transcriptomics, RRBS, and ATAC-seq datasets, we first examined the top three principal components of each tissue separately. Samples were flagged if they fell outside of three times the interquartile range for at least one of the first three principal components. Refined methods were used for remaining data types to identify extreme outliers for exclusion from differential analysis. All identified outliers were manually inspected before removal from the final dataset used for downstream analysis.

For transcriptomics datasets, principal component analysis (PCA) was run separately on the top 1000 most variable genes in each tissue, using the TMM-normalized data described above. Outliers were defined as samples outside of five times the interquartile range for any principal component that explained at least five percent of variance in the data. A total of 16 samples were removed.

For ATAC-seq datasets, PCA was run separately on the top 10,000 most variable features in each tissue and sex, using the normalized data described above. Outliers were defined as samples outside of three times the interquartile range for any principal component that explained at least 7.5% of variance in the data. A total of 11 samples were removed.

Five RRBS samples from the kidney dataset were excluded based on their sequencing quality scores (e.g., having a low total number of reads). Two additional samples, one from lung and one from hippocampus, were removed as they fell outside of five times the interquartile range of at least one of the three first principal components.

For proteomics datasets, the following samples were removed as they were flagged by PCA and also showed large differences in median TMT reporter ion intensity: two gastrocnemius samples from the global proteome and phosphoproteome datasets; one heart sample from global proteome, phosphoproteome, and acetylome datasets; and one additional heart sample only from the ubiquitylome dataset. As detailed in the experimental methods, all acetylome samples labeled with the TMT channel 130C were suspected to be failed reactions caused by the reagent lot and excluded from downstream analysis.

1-week and 2-week female vena cava samples were excluded from all analyses due to apparent brown adipose tissue contamination, as indicated by gene expression levels of *Ucp1*, a brown adipose marker, and confirmation that brown adipose contamination was sometimes visible during vena cava dissection.

### Differential analysis

All differential analyses were performed separately in each tissue and ome dataset (e.g. heart RNA-Seq). Males and females in a dataset were analyzed separately as we often observed significantly different residual variances between males and females when both sexes were fitted together. Limma with empirical Bayes variance shrinkage was used to perform differential analyses for all proteomics, metabolomics, and ATAC-seq data[56]; the edgeR pipeline for methylation analysis was used for RRBS[57]; DESeq2 was used for RNA-Seq[21]; base R linear models, *lmtest*, and *multcomp* were used for the immunoassays[58,59]. For all proteomics, ATAC-seq, immunoassays, and RRBS, the input for differential analysis was the normalized data described above. For RNA-Seq, the input was filtered raw counts, in accordance with the DESeq2 workflow. For targeted metabolomics, KNN-imputed $\log_2$-transformed data were used for datasets with more than 12 features; otherwise, $\log_2$-transformed data were used. For untargeted metabolomics, $\log_2$ KNN-imputed data were used; the sample data were median-centered if neither sample medians nor upper quartiles were significantly associated with sex or sex-stratified training group (Kruskal-Wallis p-value < 0.01).

In order to select analytes that changed over the training time course, we performed F-tests (limma, *edgeR::glmQLFTest*) or likelihood ratio tests (*DESeq2::nbinomLRT*, *lrtest*) to compare a full model with ome-specific technical covariates and training group as a factor variable (i.e. sedentary, 1 week, 2 weeks, 4 weeks, 8 weeks) against a reduced model with only technical covariates. For the immunoassays, a training group was excluded from the model if there were fewer than two non-missing values. For each feature, male- and female-specific p-values were combined using Fisher's sum of logs meta-analysis to provide a single p-value, referred to as the *training p-value*. To account for false discovery rate across all statistical tests, the training p-values were adjusted across all datasets within each ome using Independent Hypothesis Weighting (IHW) with tissue as a covariate[60]. Training-differential features were selected at 5% IHW FDR.

Given the regression model of each analyte computed as explained above, we used the contrasts of each training time point versus the sex-matched sedentary controls to calculate time- and sex-specific effect sizes, their variance, and their p-values (e.g., using linear F-tests), referred to as the *timewise summary statistics*. Specifically, for limma models we used *limma::contrasts.fit* and *limma::eBayes*, for DESeq2 models we used *DESeq2::DESeq*, for edgeR models we used *edgeR::glmQLFTest*, and for the immunoassays we used *aov()* and *multcomp::glht()*. For the immunoassays, a contrast was not tested if there were fewer than two non-missing values in either training group.

Covariates were selected from assay-specific technical metrics that explained variance in the data and were not correlated with exercise training: $\log_2$ CHEX4 for the immunoassays, which is a measure of non-specific binding[55]; RNA integrity number (RIN), median 5'-3' bias, percent of reads mapping to globin, and percent of PCR duplicates as quantified with Unique Molecular Identifiers (UMIs) for RNA-Seq; fraction of reads in peaks and library preparation batch for ATAC-seq. The same covariates were used in both differential analysis approaches described above.

#### Meta-analysis of metabolomics results

The metabolomics data were collected using different platforms from six different sites. Moreover, some platforms were *targeted* (i.e., quantified a predefined small set of metabolites of interest), whereas other platforms were *untargeted*. There were 1116 cases in which at least two sites measured the same metabolite in the same tissue. In these cases, we used meta-regression to integrate the differential analysis results, implemented using R's *metafor* package[61].

For a given metabolite m, the input to this analysis included the timewise effect sizes $y_{g,p}$ and their variances $v_{g,p}$ where g denotes the analysis group, which is a combination of the training time point and the sex for which the summary statistics were computed using the regression models explained above, and p ($\in 1,...,n_m$) denotes the platform. If m had data from at least three platforms, of which at least one was untargeted and at least one was targeted, then we added nested random effects for both the platform and the targeted status. That is, in metafor's notation we used: "mods ~ 0+analysis_group" and "random = list(~analysis_group|platform, ~analysis_group|is_targeted)". The inner|outer notation defines a blockwise dependence structure for the random effects, where different outer values are assumed to be independent, and the same outer values may be dependent based on their inner values. If the targeted status was redundant (e.g., we only had two platforms, one was targeted and the other was untargeted) then we kept the platform-level random effects only.

In practice, we observed that in 154 cases the default metafor optimizer failed to converge. In these cases, we modified the default parameters to optimizer = “nloptr”, algorithm = “NLOPT_LN_SBPLX”. Still, optimization failed in 61 additional cases. When both the default and alternative optimizers failed, we opted for a standard fixed effect model without random effects. Other than these cases, we had 361 models with random effects for both the platform and the targeted status, and 694 models with a platform-only random effect.

As a summary of each model we kept the overall model p-value (i.e., the modifiers p-value) *QMp* (used as the training p-value), and the residual heterogeneity p-value *QEp*. We flagged 103 models with QEp < 0.001 as having excessive heterogeneity. Using this definition we partitioned the meta-analysis results into three classes: (1) excessive heterogeneity, and has a targeted platform (57 cases), (2) excessive heterogeneity, without a targeted platform (46 cases), and (3) low heterogeneity (1013 cases). For class (2) we discarded the meta-analysis and kept the all platform-level results for m as-is. For class (1) we discarded the meta-analysis and kept only the targeted platform-level results for m. For class (3) we kept the meta-analysis results only (i.e., discarded the platform-level results), and used the meta-regression model to calculate the timewise summary statistics, i.e., time- and sex-specific meta-analysis effect sizes, their variance, and their p-values.

#### Graphical clustering of differential analysis results

Given the timewise differential analysis summary statistics for each analyte, computed as explained above, we next clustered the analytes into homogeneous patterns. Ovary, testis, and vena cava results were excluded from this analysis because not all time points and sexes were represented in the samples, which is a prerequisite for this longitudinal approach.

Standard clustering analysis can be used for identifying the main patterns in the data. For example, by generating a matrix of z-scores from the timewise summary statistics, where analytes are the rows and the eight training groups (e.g., 1-week males) are the columns, standard clustering algorithms can be applied. However, this methodology has three main limitations. First, off-the-shelf clustering algorithms use standard distance or correlation metrics to compute analyte pairwise distances. Thus, this may give the same distance between -1 to 1 as between 1 and 3. In the z-score space, the latter may be more important as it can signify a difference between a significant discovery (z=3) and a borderline result (z=1). Moreover, standard distance metrics do not directly account for correlations over sex and time. Second, standard clustering and fuzzy clustering analyses lead, by definition, to information loss. For example, clustering the analytes has a limited descriptive power in the presence of *split* or *convergence* points over the longitudinal trajectories. For an example of a split point in time, consider a group of features that have the same upregulation pattern up to week 4, but only a subset of the features returns to baseline levels in week 8, while the rest remain up-regulated. Clustering analysis will likely result in two clusters, without showing a direct connection between them. Finally, clustering algorithms typically disagree about the number of clusters in the underlying data, especially in noisy data. To illustrate this difficulty in our dataset we tested a few clustering algorithms and used standard evaluation metrics, as can be seen in the plot below.

A) Silhouette (kmeans, clara, hclust) B) Mixture of gaussians, BIC

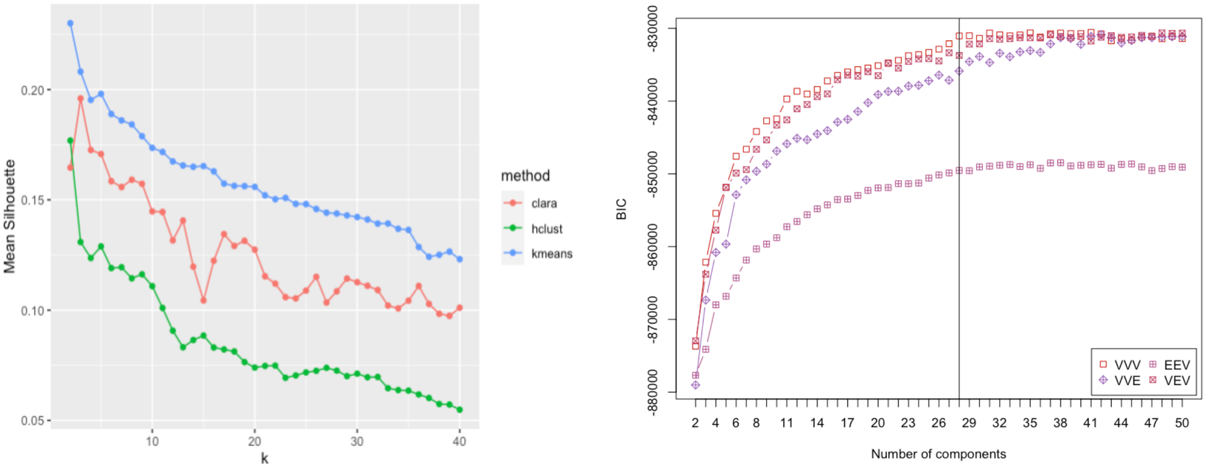


These results show that the silhouette score decreases with k for all tested algorithms, and the maximal score is <0.3, which are both indications for poor separation. Parametric mixture of gaussians clustering using the *mclust* R package[62] tends to estimate the number of clusters at 28-30, with some moderate improvement in the BIC scores at greater numbers (algorithm variants are marked as VVV, EEV, VVE, and VEV in the plot above). However, in practice we found that the solutions of these algorithms tend to result in a very large cluster around the origin, with excessive standard deviations in each axis, resulting in many significant z-scores in this “null” cluster. Thus, this algorithm does not provide reasonable results for our matrix of z-scores. Instead, we implemented a solution that overcomes the limitations of standard clustering. First, we model z-scores directly using a mixture distribution to separate null from non-null cases, thereby circumventing the limitation of naive distance metrics explained above. Second, we learn the joint distribution of events over the groups, thereby accounting for correlations over time between sexes. Finally, we utilize a graphical framework for representing the time-course of differential events. This provides an enhanced representation that allows for the identification of both split and convergence events over time.

To set the stage for our graphical clustering solution we start with some notation. Let $Z \in R^{n\ x\ t\ x\ 2}$ be the timewise z-scores, where $z_{i,j,k}$ denotes the z-score of analyte *i* ($\in$ 1,...,n) computed at the training time point *j* ($\in$ 1,...,t) of sex *k* ($\in$ {m,f}, for males and females, respectively). Assuming that $z_{i,j,k}$ follows a mixture distribution of null and non-null z-scores (i.e., a standard two-groups model), then each $z_{i,j,k}$ has a latent configuration $h_{i,j,k} \in \{-1,0,1\}$, where -1 denotes down-regulation, 0 denotes null (no change), and 1 denotes up-regulation. For simplicity, let $\mathbf{h} \in \{-1,0,1\}^{t\ x\ 2}$ denote a *full configuration matrix* (e.g., specifying if a z-score is null, up, or down for each time point in each sex), and let $\mathbf{z}_i \in R^{t\ x\ 2}$ be the matrix of all z-scores of analyte *i*. We used the expectation-maximization (EM) process of the repfdr algorithm[63,64] to estimate for each possible **h** both its prior probability π(**h**) and its posterior Pr(**h**|$\mathbf{z}_i$), for every analyte *i*. In this process, repfdr infers the marginal mixture distribution of each time point *j* and sex *k*. That is, all z-scores $z_{*jk}$ are used to estimate the densities: $f_{j,k}(z|H_{i,j,k} = -1) = f_{-1,j,k}(z)$ , $f_{j,k}(z|H_{i,j,k} = 0) = N(0,1)$, and $f_{j,k}(z|H_{i,j,k} = -1) = f_{1,j,k}(z)$. This is done using the *locfdr* R package (see [65] for details). Given these densities, the EM process is used such that Z is the observed data and H is the missing data. It iteratively updates the estimates for π(**h**) and Pr(**h**|**z**$_i$) to increase the overall complete composite likelihood (see [64] for details).

Once the algorithm converges, we discard configurations **h** with π(**h**) < 0.001 and normalize Pr(**h**|**z**$_i$) to sum to 1 (i.e., all posteriors given the same **z**$_i$). The main output of the EM for our analysis are these new posteriors that can be interpreted as a soft clustering solution, where the greater the value is, the more likely it is for analyte *i* to participate in cluster **h**. Of note, repfdr makes two simplifying assumptions about the data: (1) the z-score patterns of the analytes are independent, and (2) for a specific analyte *i*, the z-scores across the groups are independent conditioned on **h**.

Given the posteriors Pr(**h**|**z**$_i$) computed above, we assign analytes to "states". A state is a tuple ($s_{m,j}$, $s_{f,j}$), where $s_{m,j}$ is the differential abundance state null, up, or down (0,1, and -1 in the notation above, respectively) in males at time point *j*, and $s_{f,j}$ is defined similarly for females (at time point *j*). Thus, we have nine possible states in each time point. For example, assume we inspect analyte *i* in time point *j*, asking if the abundance is up-regulated in males while null in females. Then, we sum over all posteriors Pr(h|$z_i$) such that $h_{m,j}$=1 and $h_{f,j}$=0. If the resulting value is greater than 0.5, then we say that analyte *i* belongs to the node set S($s_{m,j}$, $s_{f,j}$). Thus, we use S($s_{m,j}$, $s_{f,j}$) to denote all analytes that belong to a state ($s_{m,j}$, $s_{f,j}$). Then, for every pair of states from adjacent time points *j* and *j+1* we define their edge set E($s_{m,j}$, $s_{f,j}$, $s_{m,j+1}$, $s_{f,j+1}$) as the intersection of S($s_{m,j}$, $s_{f,j}$) and S($s_{m,j+1}$, $s_{f,j+1}$). Thus, the sets S and E together define a tree structure that represent different differential patterns over sex and time.

#### Comprehensive feature-to-gene map

We compiled a feature-to-gene map that associates every feature tested in our differential analysis with gene identifiers. All proteomics feature IDs (RefSeq accessions) were mapped to gene symbols and Entrez IDs using NCBI's "gene2refseq" mapping files (https://ftp.ncbi.nlm.nih.gov/gene/DATA/gene2refseq.gz, accessed 12/18/2020). Epigenomics features were mapped to the nearest gene using the R function *ChIPseeker::annotatePeak()*[66] with Ensembl gene annotation (*Rattus norvegicus* release 96). Gene symbols, Entrez IDs, Ensembl IDs, and Rat Genome Database (RGD) IDs were mapped to each other using the RGD rat gene annotation (https://download.rgd.mcw.edu/data_release/RAT/GENES_RAT.txt, accessed 11/12/2021).

#### Pathway enrichment analysis of graphical clusters

All non-metabolite training-differential features (5% IWH FDR) were mapped to Ensembl gene symbols using the feature-to-gene map described above. Training-differential metabolites were mapped to KEGG IDs. For each graphical cluster of interest (i.e., the ten largest paths, two largest nodes, and two largest single edges with at least 20 features in each tissue), we performed pathway enrichment analysis separately for the Ensembl genes (or KEGG IDs for metabolomics) associated with differential features in each ome. For most omes, background gene (or metabolite) sets were specified as the set of Ensembl IDs (or KEGG IDs) associated with all features tested within the corresponding tissue during differential analysis. For epigenomics (ATAC-seq and RRBS), the background gene sets were defined as the set of all genes expressed in each tissue, taken from our RNA-Seq data. For gene-centric omes (i.e., all but metabolomics) we performed enrichment analysis of KEGG and REACTOME rat pathways (organism “rnorvegicus”) using the *gprofiler2::gost* function in R with the custom backgrounds defined above[67]. Only pathways with at least 10 and up to 200 members were tested. Because *gprofiler2::gost* only returns adjusted p-values, we recalculated nominal p-values using a one-tailed hypergeometric test, which is consistent with how *gprofiler2::gost* calculates enrichments. For metabolites, we performed enrichment of KEGG pathways using the hypergeometric method in the R *FELLA* package with custom backgrounds as defined above[68]. Pathway enrichment analysis p-values were adjusted across all results using Independent Hypothesis Weighting (IHW) with tissue as a covariate. We defined significantly enriched pathways as those with q-value < 0.1. Significant pathway enrichments driven by a single gene were removed from the results (Table S9).

#### Biological networks

We considered three resources for biological interactions: BioGRID (v BIOGRID-ORGANISM-4.2.193.tab3)[69], STRING (v 10116.protein.physical.links.v11.5)[70], and biological pathways. BioGRID and STRING were used for protein-protein interactions. Biological pathways contain various types of interactions, including undirected protein interactions, directed signaling interactions, and gene-metabolite pairs. For the analysis below we ignored the type (and direction). Pathway interactions were retrieved using the *graphite* R package (v 1.37.1)[71,72]. We used all directed and undirected (but not indirect) pathway interactions from Reactome, PathBank, and KEGG[73–75]. Metabolite IDs from these pathways were mapped to KEGG and InChiKey IDs (if ones were not provided). Metabolites that only had ChEBI or CAS identifiers were mapped to KEGG and InChiKey IDs using the CTS online tool (http://cts.fiehnlab.ucdavis.edu/batch). Finally, KEGG and InChiKey IDs were mapped to RefMet IDs using the Metabolomics Workbench REST service. A few extra missing KEGG IDs were added manually (as provided by chemical analysis sites).

The rat STRING network contained 115,389 high confidence interactions (score > 500), covering 10,322 genes. The graphite rat gene-gene network contained 161,130 interactions, covering 8942 genes and metabolites. The rat gene-metabolite pathways network contained 95,295 interactions, covering 5791 genes and metabolites. In contrast, the BioGRID rat network was much smaller with 6408 interactions only. To account for this limitation we used the human and mouse networks to extend the rat BioGRID network. By mapping gene symbols from these organisms to rat gene symbols via the RGD ortholog mapping (v39), we were able to generate a BioGRID rat network with 461,685 interactions, covering 15,856 genes. When analyzing data from a specific tissue, all networks were reduced to their induced subgraphs using the background gene and metabolite sets explained above.

#### Network connectivity analysis

Given an interaction network G=<V,E> (where V is a set of nodes, and E is a set of undirected edges), a “reference” set of nodes S, and a node of interest v, we tested the connectivity between v and S in G using a simple hypergeometric test. Assume a null hypothesis in which all neighbors of v in G are selected at random. Under this null hypothesis, the number of neighbors of v that are in S follows a hypergeometric distribution HG(N,K,n), where N is the number of nodes in the graph G (i.e., |V| the number of nodes in V), K is the number of neighbors of v in G, and n=|S| is the number of nodes in S. Thus, for each node v in G we can obtain a p-value for the connectivity of v and S.

To illustrate the tight connectivity of our multi-omic results in the gastrocnemius (SKM-GN), we examined the sex-consistent, up-regulated analytes identified in week 8. These data contained 64 multi-omic genes identified by at least two different omes. We used this set of genes as the reference set S, and for each of the three gene networks above we computed the p-value for every node. For a node v in S the p-value was computed by taking S\{v} as the reference set (i.e., by excluding v from S). In each network we selected the significant edges at 5% Benjamini-Hochberg FDR adjustment, resulting in 95, 51, and 326 genes in the STRING, pathways, and BioGRID networks, respectively. These sets were significantly enriched both in the multi-omic genes in S and in genes that were discovered by a single ome in our analyses (p < 10^-10 for each set using Fisher’s exact test; see Figure S5C).

Based on the analysis above we took the BioGRID network interactions and used them to generate an interaction graph of all genes and metabolites identified in SKM-GN 8w_F1_M1 (i.e., up-regulated in both sexes in week 8). We clustered the graph using the leading eigenvector clustering algorithm[76], implemented in the *igraph* R package[77]. This resulted in three large connected clusters that had at least 10 nodes (Table S10).

#### Transcription factor enrichment analysis methods

We conducted transcription factor (TF) motif enrichment analysis on sets of genes that satisfied the IHW-adjusted training p-value threshold of 0.05 for differential expression in a given tissue, isolating the 13 tissues that had at least 300 genes meet the differential expression threshold. The analysis was carried out by findMotifs.pl (HOMER v4.11.1)[78]. Enrichment of a motif in a tissue is quantified by the percent of promoter regions of differentially expressed genes containing that motif, with enrichment p-values calculated by comparing enrichment among target genes to enrichment among a background gene set that is normalized for GC%. To determine tissue similarity in motif enrichment, we calculated the mean absolute value of differences in enrichment over all TFs between each tissue and conducted hierarchical clustering of the tissues on their enrichment differences.

For more detailed analyses, we selected the top ten enriched TFs from each tissue and then removed any TFs whose corresponding genes were not found to be expressed in the RNA-Seq data. Pearson correlation was calculated between tissue-standardized TF motif enrichment scores and tissue-standardized TF gene expression across the set of control samples in each of the 13 tissues.

#### Rat-to-human ortholog map

We compiled a map between rat Ensembl IDs and human gene symbols using several external sources. GENCODE metadata and annotation files were used to map between human Ensembl transcript IDs, Entrez IDs, GENCODE IDs, and Ensembl gene IDs[79]. RGD files were used to map between human and rat gene symbols as well as between various rat gene identifiers[80].

#### Gene and PTM set enrichment analysis

Gene set enrichment analysis (GSEA) and PTM set enrichment analysis (PTM-SEA) were performed using the ssGSEA2.0 implementation by Krug et al.[81]. For GSEA, differential enrichment analysis t-scores (trained versus control) were used as input. Feature-level data was summarized into gene-level data using the maximum absolute t-score. The gene set database used included canonical pathways available through the Human Matrisome database available through MatrisomeDB (http://matrisome.org/), and the MitoPathways database available through MitoCarta 3.0[82–84]. Human gene symbols were mapped to rat orthologs before running the analysis. The *ssGSEA2* function was run using parameters that avoid normalization, require at least 5 overlapping features with the gene set, and use the area under the curve as the enrichment metric (sample.norm.type = "none", weight=0.75, correl.type = "rank", statistic = "area.under.RES", output.score.type = "NES", min.overlap=5).

For PTM-SEA, phosphosite-level t-scores were used as input. Phosphosite flanking sites were mapped from rats to humans as described below. The PTM set database used was the human PTMSigDB[81]. The *ssGSEA2* function was run using parameters that avoid sample normalization, require at least 5 overlapping features with the gene set, and use the area under the curve as the enrichment metric (sample.norm.type = "none", weight = 0.75, correl.type = "rank", statistic = "area.under.RES", output.score.type = "NES", min.overlap = 5).

#### Mapping PTMs from rat to human proteins

We used the NCBI Reference Protein Sequence database (RefSeq) to annotate protein IDs. Most of the Post-Translational Modification (PTM) resources and tools available are for humans; rat annotation is lacking. To leverage information from humans, we mapped PTM sites from rats to humans following a bioinformatics approach. Briefly, we used BLASTp[85,86] to align all rat sequences to the human review UniProt fasta sequence database (download date: 02/03/2021)[87]. The median protein sequence identity between rats and humans is 85%. Only alignments with a sequence identity greater than 60% were included for mapping. For most proteins, BLASTp outputs multiple pairwise alignments (one-to-many). In those cases, we selected the alignment with the larger "positives" and "identities" values and required an exact match for the S/T/Y residues identified in this study. As a result, we could map with confidence 73.5% of all the phosphorylation sites we identified.

#### Correlation of training-differential features with cell type markers

In order to characterize the extent to which the presence of different cell type populations were reflected in trajectories of specific clusters of training-differential features, we correlated these trajectories with transcript- and protein-level expression of cell markers. Well-documented, immune cell-type-specific surface markers classically employed in tissue staining and cell sorting procedures were selected through literature review and using resources from companies such as ThermoFisher and Biolegend (Table S17). Hemoglobin genes were added to the list of erythrocyte markers. Marker of proliferation Ki-67 (*Mki67*) was also included to reflect cell proliferation[88], and markers of lymphatic tissue were curated from the literature[89]. For each marker, we calculated the Pearson correlation between the normalized expression of the marker and the average normalized expression across all features in the given cluster of interest. Correlations were performed on sample-level data, separately for each tissue and ome (transcriptomics and proteomics). For each cell or tissue type with more than one marker, we performed a one-sample t-test to assess whether or not the corresponding correlations were significantly different from zero, separately for each tissue and ome (5% BY FDR).

#### Immune cell type deconvolution

In addition to correlating gene expression with specific immune cell type markers, we performed immune cell type deconvolution on bulk, tissue-level transcriptomic data using CIBERSORTx[90]. Ensembl gene IDs were mapped human gene symbols using the rat-to-human ortholog map described above, and raw read counts from each tissue were deconvoluted using the LM22 immune cell signature matrix generated by CIBERSORT developers[91] with the following parameters: absolute mode, B-mode batch correction, no quantile normalization, 0 permutations for significance analysis. The resulting absolute scores for each cell type in each sample were used for statistical analyses and comparisons within each tissue (Figure S8E).

#### Enrichment of LM22 immune cell types

In a parallel approach to immune cell type deconvolution, we performed enrichment of the CIBERSORT LM22 immune cell types using cell type specificity scores calculated from the LM22 leukocyte gene expression matrix[91,92]. The specificity index (SI) was calculated for each gene and cell type on a gene by cell-type matrix of the average expression for each cell type using the *pSI::specificity.index* R function with arguments e_min=0, bts=1, and SI=TRUE[93]. The resulting SI were $\log_2$-transformed and multiplied by -1 so that larger scores corresponded to greater cell type specificity. The rat-to-human ortholog map described above was used to map human gene symbols from the LM22 dataset to rat gene identifiers. For each immune cell type, tissue, and graphical node of interest (e.g., memory B cells, white adipose tissue, 8w_F1_M1), we tested for enrichment of immune cell type signatures by comparing the cell type specificity scores corresponding to the training-differential transcripts in the given tissue and cluster to the cell type specificity scores corresponding to all other genes expressed in that tissue using a one-sided Mann-Whitney U test (*wilcox.test* function in R with argument alternative = “greater”). Significant enrichments were reported at q-value < 0.05 across all tests (Figure S8F).

#### Quantification of sex differences in the training response

The extent of sex differences in the training response were characterized in two ways. First, we correlated the training $\log_2$ fold-changes between males and females for each training-differential feature in order to characterize differences in the direction of the training effect. A zero was prepended to the time-ordered vector of $\log_2$ fold-changes for each sex to indicate baseline (i.e., the $\log_2$ fold-change between the controls trained for zero weeks and themselves is zero). In order to account for spurious association due to autocorrelation[94], we first took the lagged difference of values in each vector and divided those values by the square root of the lagged differences in the time points (i.e., 0, 1, 2, 4, and 8 weeks). Then we calculated the Pearson coefficient between these transformed vectors (Figure 7A-D, Figure S9). Second, we calculated the difference between the male and female areas under the curve ($\Delta_{AUC}$, males - females) made by plotting time versus $\log_2$ fold-change for each training-differential feature, including a (0,0) point. This characterized the difference in the magnitude of response between the sexes. All $\log_2$ fold-changes for a feature were scaled to have unit variance before calculating sex-specific AUC using the *pracma::trapz* function in R[95]. Because $\Delta_{AUC}$ does not account for time, it is difficult to interpret alone. Therefore, we plotted the $\Delta_{AUC}$ against the correlation between male and female training $\log_2$ fold-changes (Figure S9) to visualize the sex difference in both magnitude (x-axis) and direction (y-axis) of the training response for all training-differential features in each tissue.

#### Comparison with human training gene expression data

We used the results from the gene expression data meta-analysis of Amar et al.[96]. In this paper, a meta-analysis was used to synthesize skeletal muscle gene expression data from 26 cohorts (n=430 subjects). Each cohort had a group of subjects that went through a training program and had a pre-training time point and at least one post-training time point. Most of the studies collected data from vastus lateralis. The paper identified 114 genes that were consistently up-regulated across the studies. However, these 114 genes were selected at extremely strict cutoffs, and extensive effect heterogeneity of many other genes were observed.

We compared our vastus lateralis transcriptomics differential analysis results with the results of the meta-analysis. First, we computed the significance of the Spearman correlation between our estimated $\log_2$ fold-changes and the estimated $\log_2$ fold-change from the meta-analysis (Figure 8A). Second, we ranked all available human genes according to their meta-analysis statistics and then performed gene set enrichment analysis (GSEA) using our identified gene sets as the sets of interest. For this analysis we used our sex-consistent differentially expressed gene sets (i.e., F1_M1, F-1_M-1, and F0_M0 according to our graphical analysis notation) from all weeks (Figure 8B). For this GSEA, the human genes were ranked using: ($-\log_{10}$ meta-analysis p-value)*(meta-analysis $\log_2$ fold-change). Finally, we checked the overlap between our gene sets and the set of genes with excessive true heterogeneity from the human data. These were defined as genes with a meta-analysis I2 > 75%. We used Fisher's exact test for testing the null hypothesis of a random overlap (Figure 8C).

#### Disease ontology enrichment analysis

We first preprocessed the disease ontology database before applying the enrichment analyses. Our rationale here was that many disease terms may be enriched with general biological processes that are relevant for many tissues both in health and disease states (e.g., cell proliferation in cancer disease terms), and are thus not likely to reflect a true association between our exercise-specific results and diseases. We therefore generated tissue-specific disease ontology terms by utilizing gene expression data from GTEx v8[97]. For each disease ontology term and a tissue (covered by GTEx) we computed the p-value for the overlap between the term's gene set and the tissue's gene set. If the p-value was greater than 0.001 then we omitted the term from the tissue's analyses. Disease ontology enrichment analysis was then performed using the *DOSE* R package[98] for each of our tissue- and ome-specific gene sets that had at least 20 genes. Identified associations were declared as significant if their FDR-adjusted p-value was <0.05 and they had at least three genes in the intersection between our set and the disease term gene set.
