## Additional File 2 - Author Contributions for "Temporal dynamics of the multi-omic response to endurance exercise training across tissues"

Joshua Adkins: Led the Proteomics Chemical Analysis Site at PNNL; collaborated with the consortium to generate the experimental design; oversaw the experimental design and implementation of proteomics workflows; reviewed data analysis platform for proteomics and proteomics interpretation; reviewed the manuscript and made suggestions for edits

Jose Juan Almagro Armenteros: Contributed to the immunological analysis of the small intestine and interpretation of the results. Contribute to the immune figure.

David Amar: Co-led the manuscript working group; compiled analysis resources and documentation. Led the statistical and computational design and analysis throughout the paper. Contributed to RNA-seq, RRBS, proteomics, and metabolomics quantification, normalization pipelines, and optimization of all differential analysis pipelines. Performed the metabolomics meta-analysis; repfdr-based Bayesian clustering of the differential analysis results; network biology analyses and tissue comparisons. Helped write, distribute, and revise the manuscript.

Mary Anne Amper: Processed GET assays, data generation, and QC.

Julian Avila-Pacheco: Performed metabolomic data QC, data preparation, analyses, and compound ID harmonization across sites. Contributed in the discussion of metabolomic related analyses in the manuscript and global tally of project samples used for each data generation platform.

Nasim Bararpour: Contributed to metabolome-related data discussion & interpretation. Contributed to supplementary immune figure panel

Bryan C Bergman: Contributed to design of study, reviewed manuscript as part of publications committee.

Sue C. Bodine: Co-lead preclinical animal study site: Contributed to the development of the animal protocol and design of the study; oversaw the implementation of the animal training protocol, participated in tissue collections and data analysis, contributed to the writing and editing of the manuscript

Frank W Booth: Contributed to data interpretation and reading and editing manuscript

Thomas Buford: Contributed to study analytic discussions; reviewed the manuscript and made suggestions for edits

Toby Chambers: Contributed to the design of the study; participated in data analysis calls, reviewed the manuscript

Clarisa Chavez: Contributed to the immunoassay literature review

Paul M. Coen: Contributed to the design of the study; participated in data analysis calls.

Elaine Cornell: University of Vermont Biorepository Senior Supervisor; oversaw all administrative aspects of the Biorepository, including MoTrPAC

Gary Cutter: Contributed to comments on analysis of results

Surendra Dasari: Processed the targeted metabolomics data.

Luis G. O. De Sousa: Assisted in animal endurance training protocols, tissue collection, data acquisition, and data entry.

Courtney Dennis: Sample processing, data generation, and QC of LC-MS metabolomics data.

Karyn A. Esser: Co-lead preclinical animal study site: Contributed to the development of the animal protocol and design of the study; helped revise and edit the manuscript

Charles Evans: Led development of cross-site MoTrPAC metabolomics method of procedure (MOP) document, prepared site-specific standard operating procedure (SOP) documents including developing and validating protocols for internal standard controlled metabolite extraction, and supervised use of these protocols to prepare tissue and plasma samples and perform untargeted metabolomics analysis

Facundo M. Fernandez: Contributed to the design and implementation of the overall lipidomics measurement approach. Led the CAS team at the Georgia Institute of Technology involved in non-targeted lipidomics measurements and lipid annotation using high resolution mass spectrometry. Conceived data dictionary approach for lipid annotation. Reviewed the manuscript and made edits.

Nicole Gagne: Biorepository technician; worked closely with Sandra May on all technical aspects of the Biorepository

David A. Gaul: Implemented non-targeted lipidomics measurement platform for a variety of tissues. Curated data dictionary for lipid annotation. Quality oversight of lipidomics datasets.

Nicole R. Gay: Co-led the manuscript working group and 6 of 8 figure subgroups; compiled analysis resources and documentation. Analyzed RNA-seq and ATAC-seq pilot data; handled all data generated by Stanford GET. Contributed to RNA-seq, ATAC-seq, immunoassay, and metabolomics quantification and/or normalization pipelines and optimization of all differential analysis pipelines. Performed all RNA-seq, all ATAC-seq, and some immunoassay differential analysis; RNA-seq, ATAC-seq, and immunoassay QC; all pathway enrichment of graphical clusters; immune cell type correlation or enrichment analyses; analyses related to global, brain, adrenal, lung, and adipose sex differences; and some multi-tissue analyses. Helped write, distribute, and revise the manuscript. Developed the MotrpacRatTraining6mo and MotrpacRatTraining6moData R packages.

Yongchao Ge: Developed the snakemake pipelines and the method of procedures (MOP) for rna-seq and rrbs data data processing and the QC.

Bret H. Goodpaster: Contributed to the design of the study; reviewed the manuscript and made suggestions for edits

Laurie J Goodyear: One of the lead preclinical animal study site Investigators. Contributed to the development of the animal protocol and design of the study; discussion and development of data.

Josh Hansen: Processed proteomic samples at PNNL

Andrea L. Hevener: Contributed to data interpretation, relevant citations, and review of the manuscript

Fang-Chi Hsu: Participated on data analysis calls, reviewed the manuscript and made suggestions for edits

Kim M. Huffman: Contributed to the design of the study; reviewed the manuscript and made suggestions for edits

Olga Ilkayeva: Led the Duke Metabolomics team that prepared samples and performed the targeted mass spec analysis across five tissue types.

Bailey E. Jackson: Assisted in animal endurance training protocols; tissue collection, data acquisition, and data entry.

Pierre M Jean Beltran: Co-led the manuscript writing and data analysis working groups. Led the analysis and biological interpretation of proteomics data and integration with other omics. Contributed to design and development of proteomics and metabolomics data processing, analysis, and visualization pipelines. Co-led the study design for proteomics, including sample processing, and data generation at the Broad Institute Proteomics Platform.

David Jimenez-Morales: Led the development of the data transfer guidelines. Developed the MotrpacBicQC package for QA/QC metabolomics and proteomics submissions. Developed the cloud implementation of the proteomics data analysis pipeline. Contributed to metabolomics and proteomics normalization and differential analysis pipelines. Coordinated and performed data preparation and releases. Co-supervised BIC staff and trainees. Co-chair the manuscript working group calls. Reviewed the manuscript and made suggestions for edits.

Christopher Jin: Contributed to the analysis/literature review of the Cytokine portion of the manuscript

Maureen Kachman: Led the Michigan MoTrPAC CAS team that prepared and analyzed samples on three Untargeted mass spectrometry platforms, which provided semiquantitative data for both unannotated and annotated metabolites across tissue types

Benjamin G. Ke: Contributed to data analysis; reviewed the manuscript

Hasmik Keshishian: Co-led the study including study design for proteomics, overseeing sample processing, data generation and analysis at the Broad Proteomics site.  Reviewed the manuscript and suggested edits

Kyle S. Kramer: Assisted in animal endurance training protocols; tissue collection, data acquisition, data entry, determined training protocol adjustments when necessary.

William E. Kraus: Contributed to the design of the study; reviewed the manuscript and made suggestions for edits

Christiaan Leeuwenburgh: Contributed to the development of the initial animal protocol and design of the study

Bridget Lester: Contributed to the design of the study

Malene E. Lindholm: Co-led the sex differences figure subgroup, performed analyses for main and supplemental figure panels, helped write and revise the manuscript, contributed to study design discussions

Ana Lira: Assisted in animal endurance training protocol, data acquisition, tissue collection, and data entry.

Xueyun Liu: Contributed to lipidomics sample prep, data collections, and data analysis at Emory University.

Kristal Maner-Smith: Contributed to data interpretation and reading and editing manuscript at Emory University

Gina M Many: Contributed to data analysis and interpretation and writing the manuscript, with primary focus on immunological aspects (e.g. small intestine, adipose); reviewed the manuscript and made suggestions for edits

Shruti Marwaha: Contributed to immunoassay QC

Sandra May: Biorepository senior technician; oversaw the development and implementation of all technical aspects of the Biorepository including kit design, sample receipt, cryopulverization, aliquot preparation, etc.

Michael E. Miller: Contributed to the design of the study; reviewed the manuscript and made suggestions for edits

Ronald J Moore: Contributed to LC-MS methods development and data acquisition at PNNL

Samuel G Moore: Performed lipidomics sample prep, data collection, and data analysis at Georgia Tech.

Kerrie L. Moreau: Contributed to the design of the study; reviewed the manuscript and made suggestions for edits

Michael Muehlbauer: Formatted and downloaded data from the Duke Metabolomics team for the MoTrPAC BIC database.

Charles C. Mundorff: Sample processing and data analysis contributions at Broad Proteomics site

Nicolas Musi: Participated on data analysis calls, reviewed the manuscript and made suggestions for edits

Venugopalan D. Nair: Co-led the Mount Sinai GET team. Oversaw assay development and implementation of GET assays, data generation, and QC.

Archana Natarajan Raja: Contributed to the development of cloud based pipelines for the analysis of epigenomics and transcriptomics data in colloboration with Sinai and Stanford GET CAS sites. Processed rna-seq, atac-seq and rrbs datasets through these standardized cloud based pipelines. Contributed to QC of the data, standardized data submission and ingestion.

Michael Nestor: Contributed to data analysis pipeline

Eric A. Ortlund: Contributed to the design and implementation of the overall targeted lipidomics measurement approach. Led the CAS team at Emory University involved in targeted lipidomics measurements using mass spectrometry and implementation of bioinformatics analyses for both targeted and untargeted data. Reviewed the manuscript and made edits.

Marco Pahor: Contributed to the design of the study; reviewed the manuscript

Paul Piehowski: Study design and oversight of data production for proteomics at PNNL

Wei-Jun Qian: Co-led the Proteomics Chemical Analysis Site at PNNL; oversaw the experimental design and implementation of proteomics workflows; reviewed the manuscript and made suggestions for edits

Megan E. Ramaker: Participated on data analysis calls, reviewed the manuscript and made suggestions for edits

Alexander Raskind: Initial processing of untargeted metabolomics data and IT support for MoTrPAC data management by the Michigan CAS

R. Scott Rector: Contributed to data interpretation and reading and editing manuscript

Collyn Z-T. Richards: Assisted in animal endurance training protocols; tissue collection, data acquisition, and data entry.

Jessica L. Rooney: Biorepository MoTrPAC Project Manager; directly responsible for all logistic (kits, labels, etc.) and technical (sample distribution design, etc.) aspects of the Biorepository

Frederique Ruf-Zamojski: Contributed to the development of experimental protocols and quality control assessment of the data, reviewed the manuscript

Scott Rushing: Oversight of the programming of data collection, performed data transfer to Bioinformatics core, reviewed the manuscript and made suggestions for edits

Tyler J. Sagendorf: Contributed R code to generate enrichment heatmaps

James A. Sanford: Study design and oversight of data production for proteomics at PNNL; contibuted to the development of proteomics data analysis pipeline and interpretation; co-led the immune/inflammatory figure subgroup; participated in figure generation, manuscript writing and revision.

Simon Schenk: Co-investigator for preclinical animal study site: Contributed to the development of the animal protocol and design of the study; contributed to data interpretation, and the writing and editing of the manuscript

Stuart C. Sealfon: Co-led a data generation site. Writing group member. Revised the manuscript.

Nitish Seenarine: Performed and processed GET assays, data generation, and QC as a member of the Mount Sinai GET team. Developed automated Methyl Capture protocol for use

Gregory R. Smith: Co-led the sex differences figure subgroup, aided with design of analysis plans, conducted transcription factor motif enrichment analysis, studied cross-tissue sex differences, and helped write the manuscript

Kevin  S. Smith: Stanford GET Project Manager

Tanu Soni: Processed Michigan metabolomics data QC and normalization.

Alec Steep: Processed metabolomics datasets comprehensively and generated official metabolomics data freezes. Performed and developed QC, cross-technology correlation, and differential analyses to guide metabolomics data processing decisions.

Cynthia L. Stowe: Contributed to the design of the study, developed manual of procedures, oversaw the development of data collection forms, performed quality control of data, reviewed the manuscript and made suggestions for edits

Yifei Sun: Contributed data analysis and data generation. Reviewed protocols used in manuscript

John Thyfault: Contributed to interpreting and development of the data and editing the manuscript

Russell Tracy: Biorepository Director; as Vice-Chair of the Steering Committee, contributed to overall study design, and all aspects of sample handling and distribution to analytical sites

Scott Trappe: Contributed to the design of the study, participated in data analysis calls, reviewed the manuscript

Todd Trappe: Contributed to the design of the study, participated in data analysis calls, reviewed the manuscript

Mital Vasoya: Processed samples and QC for epigenetic data generation

Nikolai G. Vetr: Performed analyses for and generated multiple iterations of main and supplemental figure panel drafts, contributed final figure panels, provided advice and implementation details for multiple additional panels, wrote corresponding manuscript text

Michael P. Walkup: Performed all rat randomizations, performed quality control of data, reviewed the manuscript and made suggestions for edits

Martin J. Walsh: Co-led a data generation site. Reviewed manuscript and made minor suggestions for edits.

Matthew T. Wheeler: Co-led the Bioinformatics Center. Contributed to design, QC and data analysis. Supervised BIC staff and trainees. Oversaw motrpac-datahub development and data repository. Reviewed the manuscript and made suggestions for edits.

Si Wu: Contribute to metabolic-centered figure and discussion, molecule clustering analysis exploration

Ashley Xia: Contributed to the design of the study, planning of omics assay and data management; reviewed the manuscript and made suggestions for edits

Zhen Yan: Contribute to the editing of the manuscript

Chongzhi Zang: Contributed to data analysis; reviewed the manuscript

Elena Zaslavsky: Contributed to the development of GET QC data standards and processing pipelines, reviewed the manuscript

Tiantian Zhang: Contributed to the Emory Cas Site QC analysis, data processing and performed metabolite chemical class enrichment analysis of landscape paper

Bingqing Zhao: Led or co-led the Stanford GET team on protocol optimization, method development, automation and data generation of ATAC-seq, RNA-seq and immunoassays. Co-led analysis of ATAC-seq and immunoassays. Co-led immuno figure subgroup. Contributed to manuscript writing.

Jimmy Zhen: Contributed to development of MoTrPAC Data Hub and cloud computing infrastructure; contributed to design of figure in the manuscript.
