## Additional File 3 - Supplementary Figures for "Temporal dynamics of the multi-omic response to endurance exercise training across tissues"

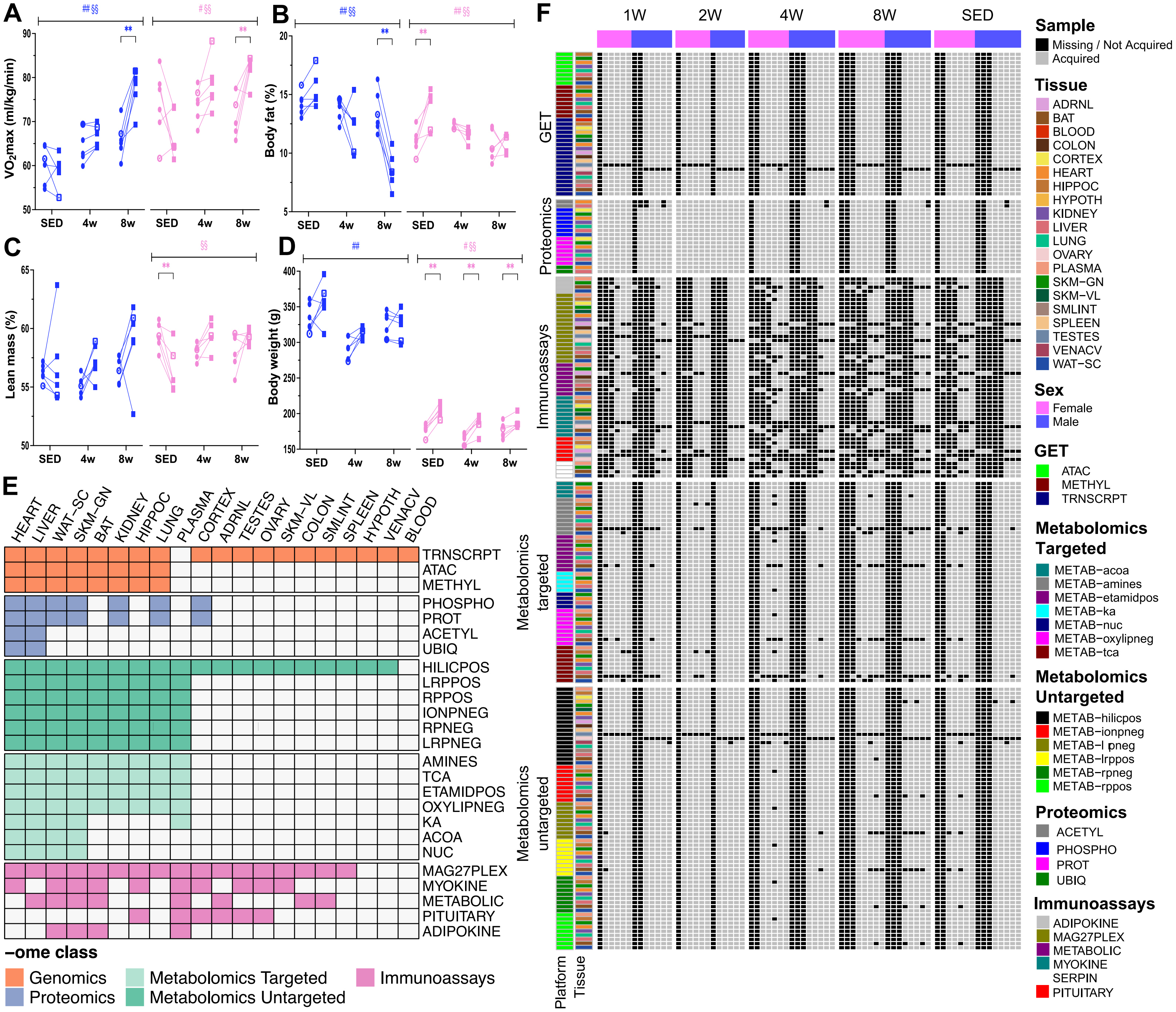

A
B
C
D
E
F
VO2max (ml/kg/min)
Body fat (%)
Lean mass (%)
Body weight (g)
SED
4w
8w
HEART LIVER WAT-SC SKM-GN BAT KIDNEY HIPPOC LUNG PLASMA CORTEX ADRNL TESTES OVARY SKM-VL COLON SMLINT SPLEEN HYPOTH VENACV BLOOD
TRNSCRPT
ATAC
METHYL
PHOSPHO
PROT
ACETYL
UBIQ
HILICPOS
LRPPOS
RPPOS
IONPNEG
RPNEG
LRPNEG
AMINES
TCA
ETAMIDPOS
OXYLIPNEG
KA
ACOA
NUC
MAG27PLEX
MYOKINE
METABOLIC
PITUITARY
ADIPOKINE
-ome class
Genomics
Proteomics
Metabolomics Targeted
Metabolomics Untargeted
Immunoassays
1W
2W
4W
8W
SED
GET
Proteomics
Immunoassays
Metabolomics targeted
Metabolomics untargeted
Platform
Tissue
Sample
Missing / Not Acquired
Acquired
Tissue
ADRNL
BAT
BLOOD
COLON
CORTEX
HEART
HIPPOC
HYPOTH
KIDNEY
LIVER
LUNG
OVARY
PLASMA
SKM-GN
SKM-VL
SMLINT
SPLEEN
TESTES
VENACV
WAT-SC
Sex
Female
Male
GET
ATAC
METHYL
TRNSCRPT
Metabolomics Targeted
METAB-acoa
METAB-amines
METAB-etamidpos
METAB-ka
METAB-nuc
METAB-oxylipneg
METAB-tca
Metabolomics Untargeted
METAB-hilicpos
METAB-ionpneg
METAB-lrpneg
METAB-lrppos
METAB-rpneg
METAB-rppos
Proteomics
ACETYL
PHOSPHO
PROT
UBIQ
Immunoassays
ADIPOKINE
MAG27PLEX
METABOLIC
MYOKINE
SERPIN
PITUITARY

### Supplementary Figure 1. Animal phenotyping and data availability.

**A)** Clinical measurements before and after the training intervention in untrained control rats (SED), the 4-week trained rats (4w), and the 8-week trained rats (8w). Data are displayed pre and post for each individual rat (connected by a line), with males in blue and females in pink. Filled symbols (n=5 per sex and time point) represent rats used for all omics analyses, whereas the rat utilized for proteomics only (n=1 per sex and time point) is represented by a non-filled symbol. Significant results by ANOVA of the overall group effect (#, $p < 0.05$; ##, $p < 0.01$) and interaction between group and time (§, $p < 0.05$; §§ $p < 0.01$) are indicated. Significant within-group differential responses from a Bonferroni post hoc test are indicated (*, q-value < 0.05; **, q-value < 0.01). **A)** Aerobic capacity through a $VO_2$max test until exhaustion . Data are reported in ml/(kg.min) for all individual rats and time points. **B)** Body fat percentage. **C)** Percent lean mass assessed through nuclear magnetic resonance spectroscopy. **D)** Body weight (in grams). **E)** Description of available datasets. Colored cells indicate that data are available for that tissue and assay. Individual panels and platforms are shown for metabolomics and the multiplexed immunoassays. **F)** Detailed availability of sample-level data across assays. Each column represents an individual animal, ordered by training group and colored by sex. Gray cells indicate that data were generated for that animal and assay; black cells indicate that data were not generated. Rows are ordered by ome and colored by assay and tissue.

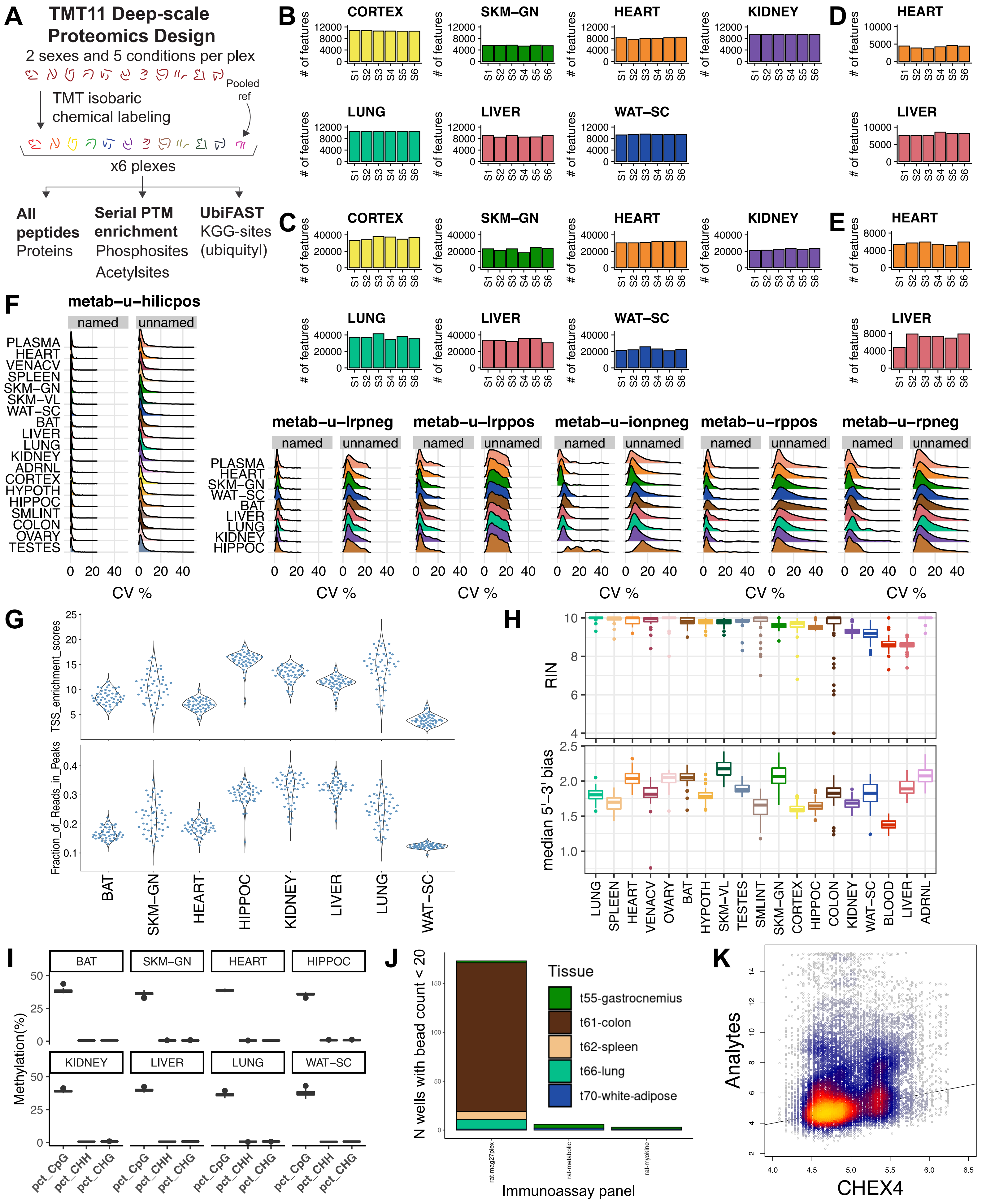
A
TMT11 Deep-scale Proteomics Design
2 sexes and 5 conditions per plex
TMT isobaric chemical labeling
Pooled ref
x6 plexes
All peptides
Proteins
Serial PTM enrichment
Phosphosites
Acetylsites
UbiFAST
KGG-sites (ubiquityl)
B
C
D
E
CORTEX
SKM−GN
HEART
KIDNEY
LUNG
LIVER
WAT−SC
### of features
S1 S2 S3 S4 S5 S6
F
metab−u−hilicpos
metab−u−lrpneg
metab−u−lrppos
metab−u−ionpneg
metab−u−rppos
metab−u−rpneg
named
unnamed
PLASMA
HEART
VENACV
SPLEEN
SKM−GN
SKM−VL
WAT−SC
BAT
LIVER
LUNG
KIDNEY
ADRNL
CORTEX
HYPOTH
HIPPOC
SMLINT
COLON
OVARY
TESTES
CV %
G
TSS_enrichment_scores
Fraction_of_Reads_in_Peaks
BAT
SKM−GN
HEART
HIPPOC
KIDNEY
LIVER
LUNG
WAT−SC
H
RIN
median 5'−3' bias
LUNG
SPLEEN
HEART
VENACV
OVARY
BAT
HYPOTH
SKM−VL
TESTES
SMLINT
SKM−GN
CORTEX
HIPPOC
COLON
KIDNEY
WAT−SC
BLOOD
LIVER
ADRNL
I
Methylation(%)
BAT
SKM−GN
HEART
HIPPOC
KIDNEY
LIVER
LUNG
WAT−SC
pct_CpG
pct_CHH
pct_CHG
J
N wells with bead count < 20
Tissue
t55-gastrocnemius
t61-colon
t62-spleen
t66-lung
t70-white-adipose
rat-mag27plex
rat-metabolic
rat-myokine
Immunoassay panel
K
Analytes
CHEX4

### Supplementary Figure 2. Quality control metrics for omics data.

**A)** The proteomics multiplexing design using TMT11 reagents for isobaric tagging and a pooled reference sample. The diagram describes processing of a single tissue. Following multiplexing, peptides were used for protein abundance analysis, serial PTM enriched for phosphosite and optional acetylsite quantification, or ubiquitylsite quantification through enrichment of lysine-diglycine ubiquitin remnants. **B)** The total number of fully quantified proteins per plex in each global proteome dataset. **C-E)** The total number of fully quantified phosphosites **(C)**, acetylsites **(D)**, and ubiquitylsites **(E)** per plex in each dataset. **F)** Distributions of coefficients of variation (CVs) calculated from metabolomics features identified in pooled samples and analyzed periodically throughout liquid chromatography-mass spectrometry runs. CVs were aggregated and plotted separately for named and unnamed metabolites. **G)** Transcription start site (TSS) enrichment (top) and fraction of reads in peaks (FRiP, bottom) across ATAC-seq samples per tissue. **H)** Distributions of RNA integrity numbers (RIN, top) and median 5' to 3' bias (bottom) across samples in each tissue in the RNA-Seq data. The average RIN was 9.6, indicating high-quality RNA. The average median 5' to 3' bias was 1.83. **I)** Percent methylation of CpG, CHG and CHH sites in the RRBS data. Non-CpG methylation of CHH and CHG sites averaged to 0.74 and 0.56 respectively, whereas average CpG methylation was 37.7. **J)** Number of wells across multiplexed immunoassays with fewer than 20 beads. Measurements from these 182 wells were excluded. **K)** 2D density plot of targeted analytes' mean fluorescence intensity (MFI) versus corresponding CHEX4 MFI from the same well for each multiplexed immunoassay measurement, where CHEX4 is a measure of non-specific binding.

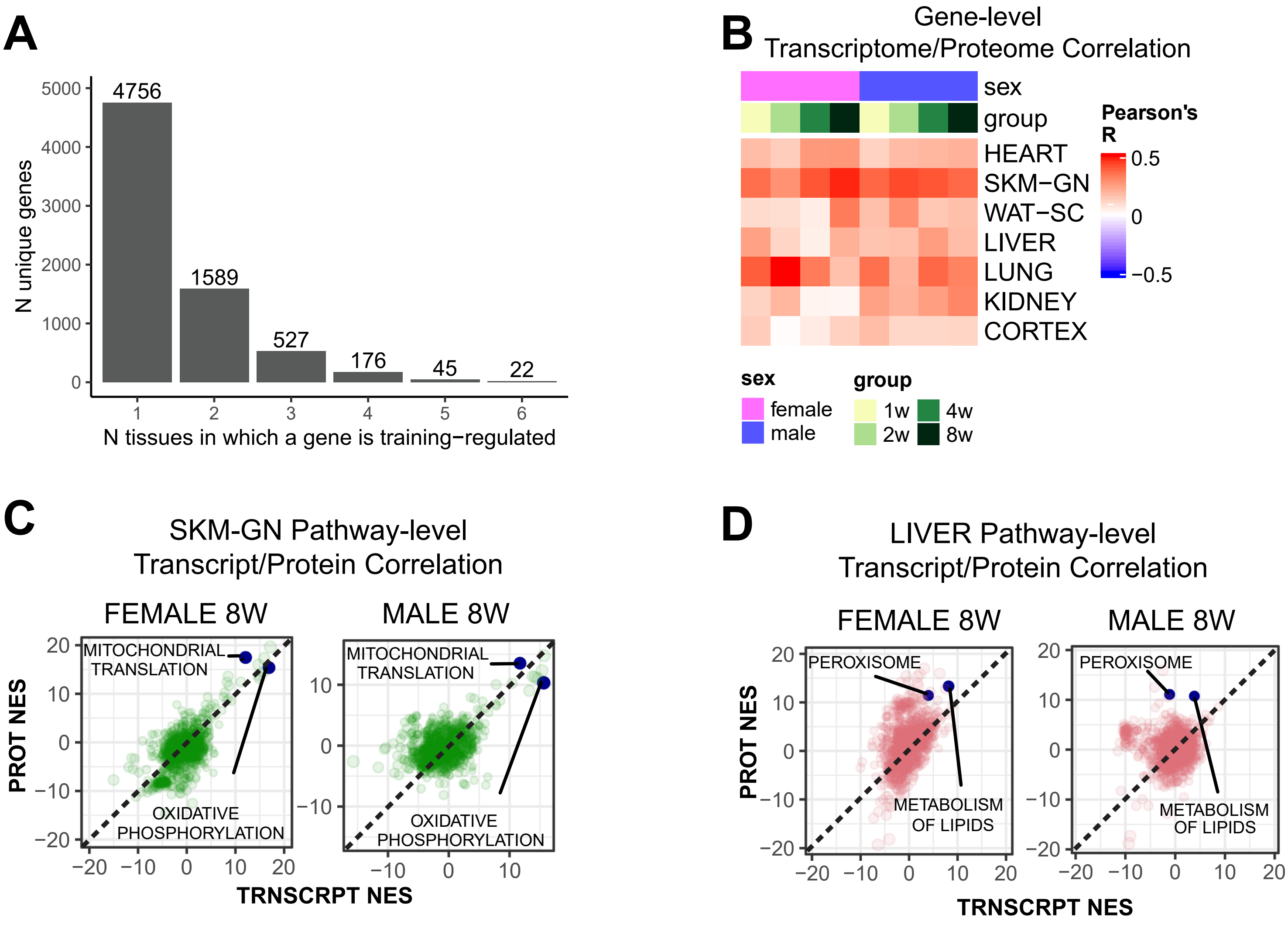
A
N unique genes
5000
4000
3000
2000
1000
0
4756
1589
527
176
45
22
1
2
3
4
5
6
N tissues in which a gene is training−regulated
B
Gene-level
Transcriptome/Proteome Correlation
sex
group
HEART
SKM−GN
WAT−SC
LIVER
LUNG
KIDNEY
CORTEX
Pearson's R
0.5
0
−0.5
sex
female
male
group
1w
2w
4w
8w
C
SKM-GN Pathway-level
Transcript/Protein Correlation
FEMALE 8W
MALE 8W
MITOCHONDRIAL TRANSLATION
OXIDATIVE PHOSPHORYLATION
PROT NES
TRNSCRPT NES
D
LIVER Pathway-level
Transcript/Protein Correlation
FEMALE 8W
MALE 8W
PEROXISOME
METABOLISM OF LIPIDS
PROT NES
TRNSCRPT NES

### Supplementary Figure 3. Correlations between proteins and transcripts throughout endurance training.

**A)** Number of tissues in which each gene, including features mapped to genes from all omes, is training-regulated. Only differential features from the subset of tissues with deep molecular profiling (lung, gastrocnemius, subcutaneous white adipose, kidney, liver, and heart) and the subset of omes that were profiled in all six of these tissues (DNA methylation, chromatin accessibility, transcriptomics, global proteomics, phosphoproteomics, multiplexed immunoassays) were considered. Numbers above each bar indicate the number of genes that are differential in exactly the number of tissues indicated on the x-axis. **B)** Heatmap showing the Pearson correlation between the TRNSCRPT and PROT timewise summary statistics (z- and t-scores, respectively). Pearson correlation was calculated for each tissue, sex, and time point combination **C-D)** Scatter plots of pathway GSEA NES of the TRNSCRPT and PROT datasets in representative tissues: **C)** gastrocnemius (SKM-GN) and **D)** liver. Labeled points represent selected pathways showing similar enrichment across both ome and sex for SKM-GN and only in females for LIVER.

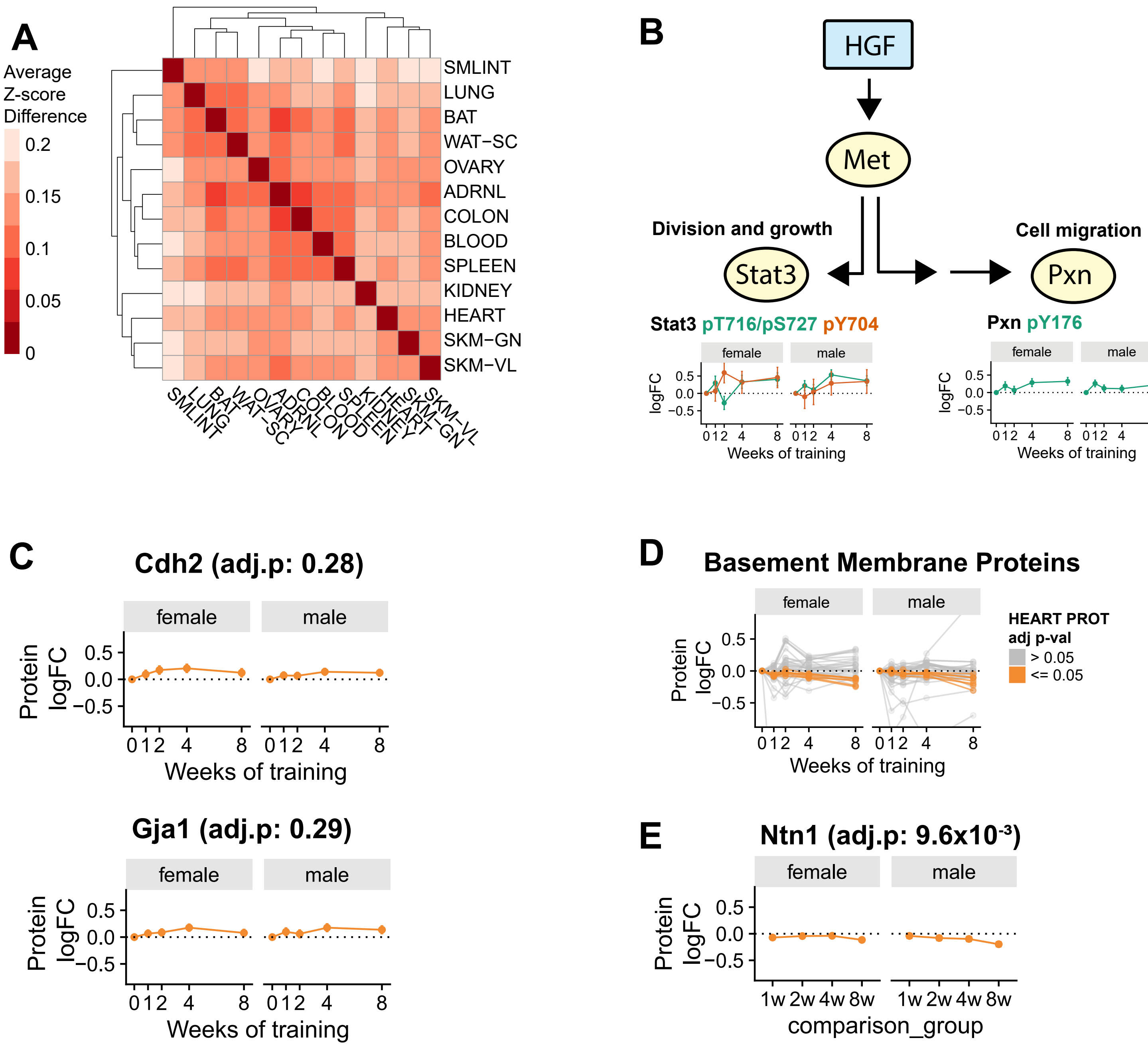

A
Average Z-score Difference
0.2
0.15
0.1
0.05
0
SMLINT
LUNG
BAT
WAT−SC
OVARY
ADRNL
COLON
BLOOD
SPLEEN
KIDNEY
HEART
SKM−GN
SKM−VL
B
HGF
Met
Division and growth
Cell migration
Stat3
Pxn
Stat3 pT716/pS727 pY704
Pxn pY176
female
male
logFC
0.5
0.0
−0.5
0 1 2 4 8
Weeks of training
C
Cdh2 (adj.p: 0.28)
Gja1 (adj.p: 0.29)
Protein logFC
D
Basement Membrane Proteins
HEART PROT adj p-val
> 0.05
<= 0.05
E
Ntn1 (adj.p: 9.6x10-3)
1w 2w 4w 8w
comparison_group

### Supplementary Figure 4. Regulatory signaling pathways modulated by endurance training.

**A)** Heatmap of differences in TF motif enrichment in training-regulated genes across tissues. Each value reflects the average difference in motif enrichment for shared transcription factors. Tissues are clustered with complete linkage hierarchical clustering. **B)** Hypothetical model of HGF signaling effects during exercise training. Phosphorylation of STAT3 and PXN is known to modulate cell growth and cell migration, respectively. **C)** $Log_2$ protein fold-change of GJA1 and CDH2 protein abundance in the heart. No significant response to exercise training was observed for these proteins (q-value > 0.05). **D)** $Log_2$ fold-change for basement membrane proteins in heart. Proteins showing a significant response to exercise training are highlighted in orange (q-value < 0.05). **E)** $Log_2$ protein fold-change of NTN1 protein abundance in heart. A significant response to exercise training was observed for these proteins (q-value < 0.05).

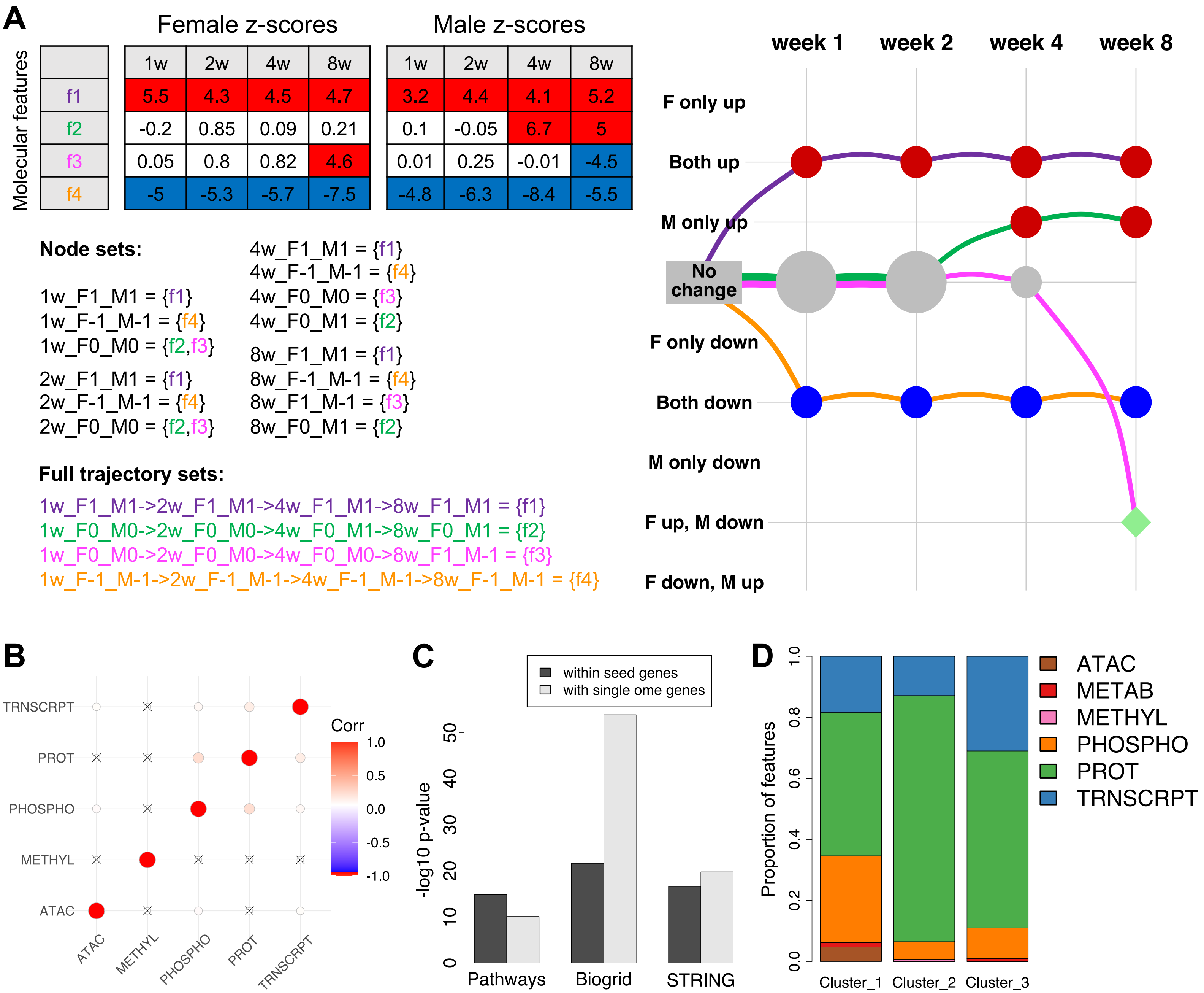

A
Female z-scores
Male z-scores
Molecular features
1w
2w
4w
8w
f1 5.5 4.3 4.5 4.7 3.2 4.4 4.1 5.2
f2 -0.2 0.85 0.09 0.21 0.1 -0.05 6.7 5
f3 0.05 0.8 0.82 4.6 0.01 0.25 -0.01 -4.5
f4 -5 -5.3 -5.7 -7.5 -4.8 -6.3 -8.4 -5.5
Node sets:
1w_F1_M1 = {f1}
1w_F-1_M-1 = {f4}
1w_F0_M0 = {f2,f3}
2w_F1_M1 = {f1}
2w_F-1_M-1 = {f4}
2w_F0_M0 = {f2,f3}
4w_F1_M1 = {f1}
4w_F-1_M-1 = {f4}
4w_F0_M0 = {f3}
4w_F0_M1 = {f2}
8w_F1_M1 = {f1}
8w_F-1_M-1 = {f4}
8w_F1_M-1 = {f3}
8w_F0_M1 = {f2}
Full trajectory sets:
1w_F1_M1->2w_F1_M1->4w_F1_M1->8w_F1_M1 = {f1}
1w_F0_M0->2w_F0_M0->4w_F0_M1->8w_F0_M1 = {f2}
1w_F0_M0->2w_F0_M0->4w_F0_M0->8w_F1_M-1 = {f3}
1w_F-1_M-1->2w_F-1_M-1->4w_F-1_M-1->8w_F-1_M-1 = {f4}
week 1
week 2
week 4
week 8
F only up
Both up
M only up
No change
F only down
Both down
M only down
F up, M down
F down, M up
B
TRNSCRPT
PROT
PHOSPHO
METHYL
ATAC
Corr
1.0
0.5
0.0
-0.5
-1.0
C
within seed genes
with single ome genes
-log10 p-value
0
10
20
30
40
50
Pathways
Biogrid
STRING
D
Proportion of features
0.0
0.2
0.4
0.6
0.8
1.0
Cluster_1
Cluster_2
Cluster_3
ATAC
METAB
METHYL
PHOSPHO
PROT
TRNSCRPT

### Supplementary Figure 5. Graphical representation of differential results and network analysis.

**A)** A schematic example of the graphical representation of the differential analysis results. Top left: the z-scores of four features. A positive score corresponds to up-regulation (red), and a negative score corresponds to down regulation (blue). Bottom left: the assignment of features to node sets and full path sets (edge sets are not shown for conciseness but can be easily inferred from the full paths). Node labels follow the [time]_F[x]_M[y] format where [time] shows the animal sacrifice week and can take one of (1w, 2w, 4w, or 8w), and [x] and [y] are one of (-1,0,1), corresponding to down-regulation, no effect, and up-regulation, respectively. Right: the graphical representation of the feature sets. Columns are training time points, and rows are the differential abundance states. Node and edge sizes are proportional to the number of features that are assigned to each set. **B)** Comparison of gene sets identified by different omes in 8-week up-regulated features from the gastrocnemius. The matrix shows the Jaccard coefficient between the gene sets. Cells in which the overlap was not significant are marked by an “X”. **C)** Results of the network connectivity analysis. For each gene in each network (BioGRID, STRING, pathways), we tested its connectivity to the “multi-omic” genes (i.e., genes identified by at least two omes) using a hypergeometric test. This analysis resulted in a set of highly connected genes in each network. The plot shows the $-\log_{10}$ p-values for the overlap between the highly connected genes and: (1) the multi-omic genes; (2) genes identified by a single ome. **D)** The proportion of features from each ome represented in the gastrocnemius response clusters, identified by the network analysis.

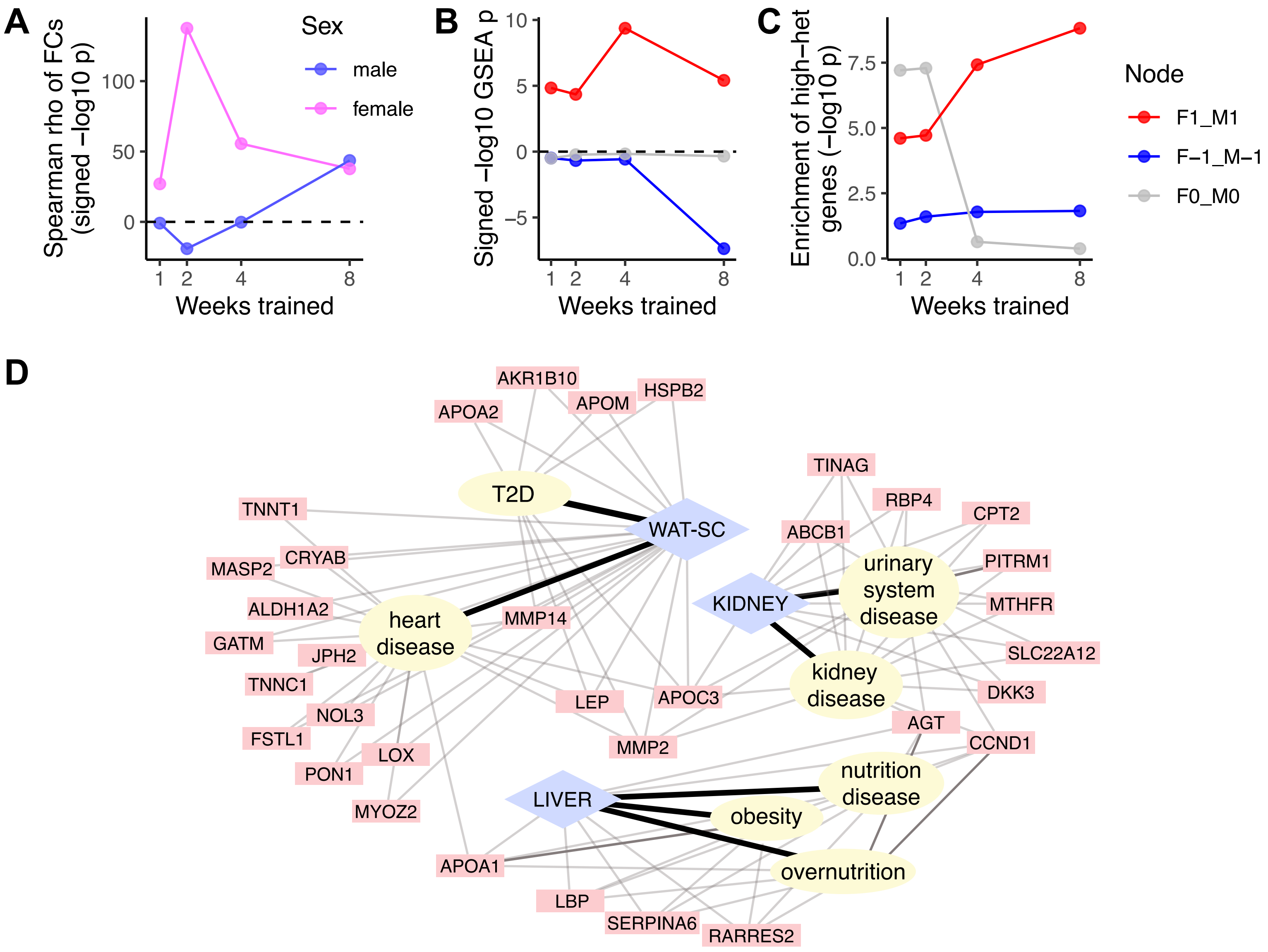

A
Spearman rho of FCs (signed −log10 p)
Weeks trained
Sex
male
female
B
Signed −log10 GSEA p
Weeks trained
C
Enrichment of high−het genes (−log10 p)
Weeks trained
Node
F1_M1
F−1_M−1
F0_M0
D
AKR1B10
HSPB2
APOM
APOA2
TINAG
T2D
RBP4
TNNT1
CPT2
WAT-SC
ABCB1
PITRM1
CRYAB
MASP2
urinary system disease
KIDNEY
MTHFR
ALDH1A2
heart disease
MMP14
GATM
SLC22A12
JPH2
kidney disease
TNNC1
LEP
APOC3
DKK3
NOL3
AGT
FSTL1
MMP2
CCND1
LOX
PON1
nutrition disease
LIVER
MYOZ2
obesity
overnutrition
APOA1
LBP
SERPINA6
RARRES2

### Supplementary Figure 6. Associations with signatures of human health and complex traits.

**A-C)** Overlap between the rat vastus lateralis differential expression results and the meta-analysis of human long-term exercise studies by Amar et al. (Amar et al., 2021b). **A)** Significance of the Spearman correlation between the fold-change estimates of vastus lateralis and the human skeletal muscle meta-analysis by (Amar et al., 2021b). Each data point is the $-\log_{10}$ p-value of the correlation between the meta-analysis fold-change and our $\log_2$ fold-changes in a specific sex and time point. **B)** GSEA results. Genes were ranked by (meta-analysis $-\log_{10}$ p-value)*(meta-analysis $\log_2$ fold-change), and rat training-differential, sex-consistent gene sets were tested for enrichment at the bottom of the ranking (negative scores) or the top (positive scores). **C)** Overlap between rat training-differential, sex-consistent gene sets and the high-heterogeneity human meta-analysis gene sets ($I^2$ > 75%). **D)** Disease ontology enrichment results of the white adipose, kidney, and liver gene sets. Enrichment analysis using DOSE was performed for each feature set, limiting the analysis for the relevant disease terms in each tissue. The network shows the results for the sex-consistent down-regulated features at week 8 (i.e., the 8w_F-1_M-1 graphical node). The omes that correspond to the feature sets in this panel are available in Table S12.

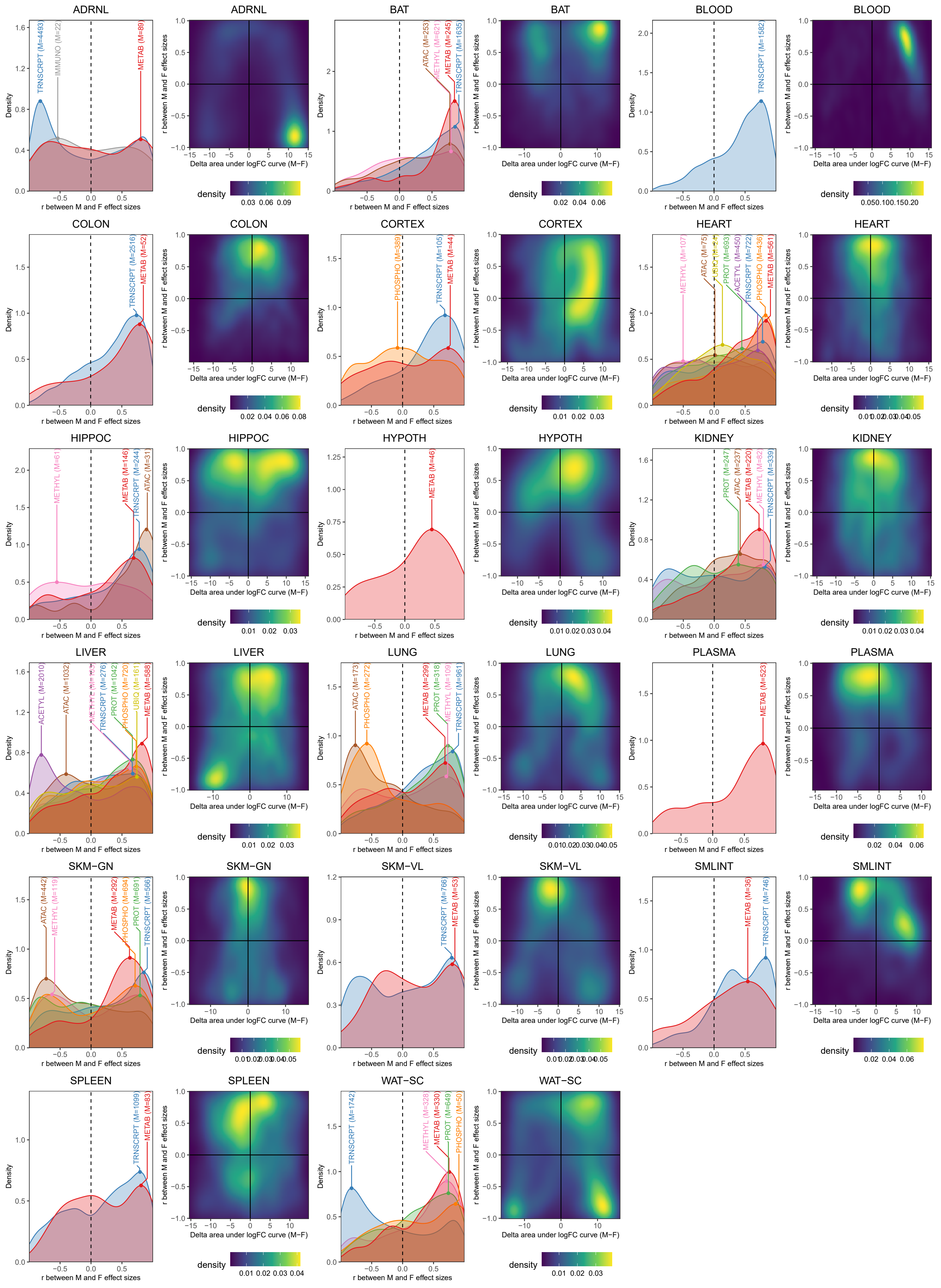

ADRNL
TRNSCRPT (M=4493)
IMMUNO (M=22)
METAB (M=89)
BAT
ATAC (M=253)
METHYL (M=621)
METAB (M=245)
TRNSCRPT (M=1635)
BLOOD
TRNSCRPT (M=1582)
COLON
TRNSCRPT (M=2516)
METAB (M=52)
CORTEX
PHOSPHO (M=389)
TRNSCRPT (M=105)
METAB (M=44)
HEART
METHYL (M=107)
ATAC (M=75)
PROT (M=693)
ACETYL (M=450)
TRNSCRPT (M=722)
PHOSPHO (M=436)
METAB (M=561)
HIPPOC
METHYL (M=61)
METAB (M=146)
TRNSCRPT (M=244)
ATAC (M=31)
HYPOTH
METAB (M=46)
KIDNEY
PROT (M=247)
ATAC (M=237)
METAB (M=220)
METHYL (M=82)
TRNSCRPT (M=339)
LIVER
ACETYL (M=2010)
ATAC (M=1032)
TRNSCRPT (M=276)
PROT (M=1042)
PHOSPHO (M=720)
UBIQ (M=161)
METAB (M=588)
LUNG
ATAC (M=173)
PHOSPHO (M=272)
METAB (M=299)
PROT (M=318)
METHYL (M=109)
TRNSCRPT (M=961)
PLASMA
METAB (M=523)
SKM−GN
ATAC (M=442)
METHYL (M=119)
METAB (M=292)
PHOSPHO (M=694)
PROT (M=691)
TRNSCRPT (M=566)
SKM−VL
TRNSCRPT (M=766)
METAB (M=53)
SMLINT
METAB (M=36)
TRNSCRPT (M=746)
SPLEEN
TRNSCRPT (M=1099)
METAB (M=83)
WAT−SC
TRNSCRPT (M=1742)
METHYL (M=328)
METAB (M=330)
PROT (M=649)
PHOSPHO (M=50)
Density
r between M and F effect sizes
Delta area under logFC curve (M−F)
density

### Supplementary figure 7. Characterization of the extent of sex difference in the endurance training response.

The extent of sex differences in the training response were characterized in two ways: first, by correlating $\log_2$ fold-changes between males and females for each training-differential feature; second, by calculating the difference between the area under the $\log_2$ fold-change curve for each training-differential feature, including a (0,0) point ($\Delta_{AUC}$, males - females). The first approach characterizes differences in direction of effect while the second approach characterizes differences in magnitude. We plotted density line plots of correlations from the first approach (left plot for each tissue). Densities or correlations corresponding to features in each ome are plotted separately, with a label that provides the ome and the number of differential features represented. We also plotted $\Delta_{AUC}$ against the correlation between the male and female $\log_2$ fold-changes for each training-differential feature (2D density plots, right plot for each tissue) to simultaneously evaluate sex differences in the direction and magnitude of the training response. Points at the top-center of these 2D density plots represent features with high similarity between males and females in terms of both direction and magnitude; features on the right and left sides of the plots represent features with greater magnitudes of response in males and females, respectively.

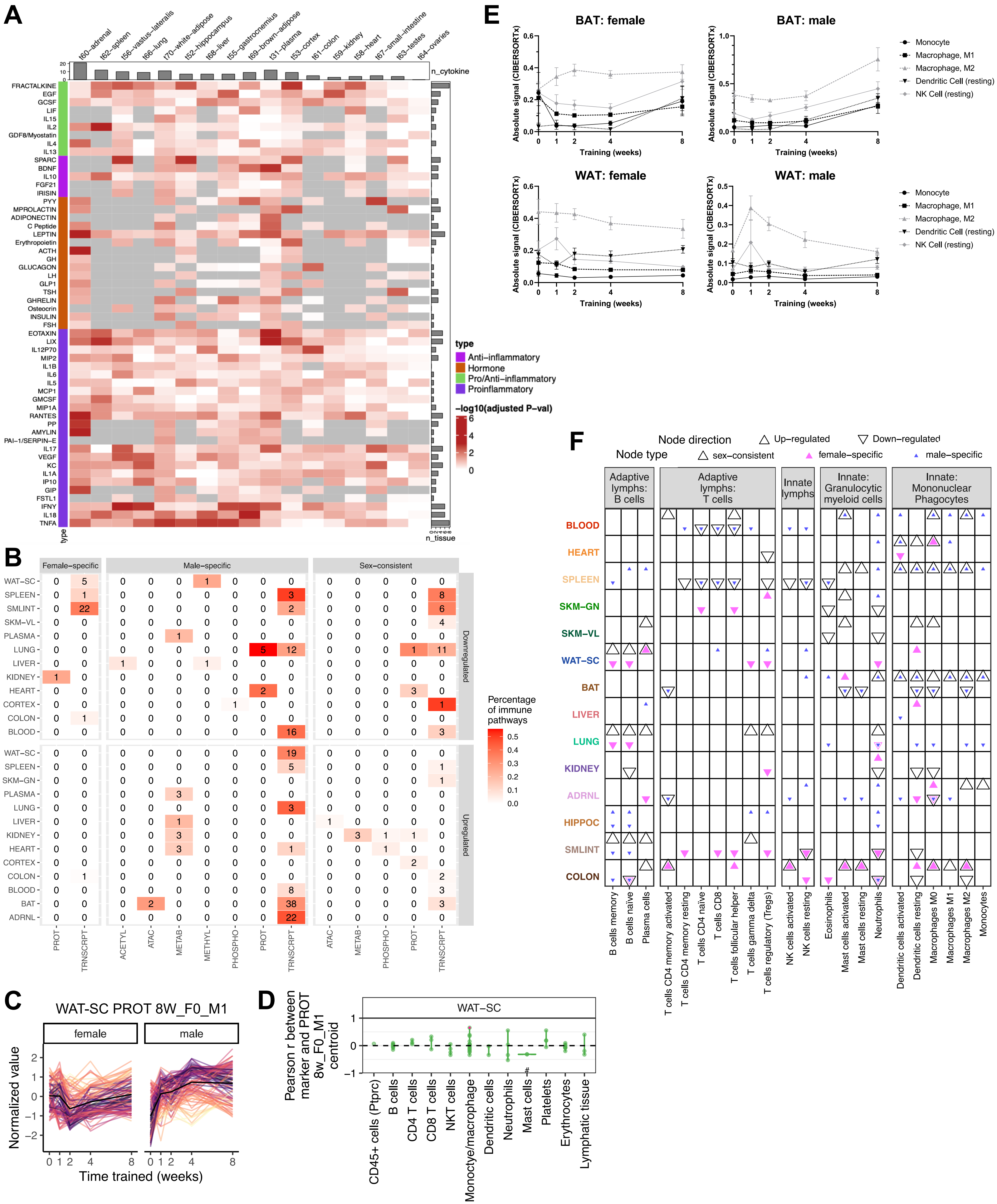

A
n_cytokine
n_tissue
type
t60–adrenal
t62–spleen
t56–vastus–lateralis
t66–lung
t70–white–adipose
t52–hippocampus
t68–liver
t55–gastrocnemius
t69–brown–adipose
t31–plasma
t53–cortex
t61–colon
t59–kidney
t58–heart
t67–small–intestine
t63–testes
t64–ovaries
FRACTALKINE
EGF
GCSF
LIF
IL15
IL2
GDF8/Myostatin
IL4
IL13
SPARC
BDNF
IL10
FGF21
IRISIN
PYY
MPROLACTIN
ADIPONECTIN
C Peptide
LEPTIN
Erythropoietin
ACTH
GH
GLUCAGON
LH
GLP1
TSH
GHRELIN
Osteocrin
INSULIN
FSH
EOTAXIN
LIX
IL12P70
MIP2
IL1B
IL6
IL5
MCP1
GMCSF
MIP1A
RANTES
PP
AMYLIN
PAI–1/SERPIN–E
IL17
VEGF
KC
IL1A
IP10
GIP
FSTL1
IFNY
IL18
TNFA
type
Anti–inflammatory
Hormone
Pro/Anti–inflammatory
Proinflammatory
–log10(adjusted P–val)
B
Female–specific
Male–specific
Sex–consistent
Downregulated
Upregulated
Percentage of immune pathways
WAT–SC
SPLEEN
SMLINT
SKM–VL
PLASMA
LUNG
LIVER
KIDNEY
HEART
CORTEX
COLON
BLOOD
SKM–GN
BAT
ADRNL
PROT
TRNSCRPT
ACETYL
ATAC
METAB
METHYL
PHOSPHO
C
WAT-SC PROT 8W_F0_M1
female
male
Normalized value
Time trained (weeks)
D
WAT–SC
Pearson r between marker and PROT 8w_F0_M1 centroid
CD45+ cells (Ptprc)
B cells
CD4 T cells
CD8 T cells
NKT cells
Monoctye/macrophage
Dendritic cells
Neutrophils
Mast cells
Platelets
Erythrocytes
Lymphatic tissue
E
BAT: female
BAT: male
WAT: female
WAT: male
Absolute signal (CIBERSORTx)
Training (weeks)
Monocyte
Macrophage, M1
Macrophage, M2
Dendritic Cell (resting)
NK Cell (resting)
F
Node direction
Up–regulated
Down–regulated
Node type
sex–consistent
female–specific
male–specific
Adaptive lymphs: B cells
Adaptive lymphs: T cells
Innate lymphs
Innate: Granulocytic myeloid cells
Innate: Mononuclear Phagocytes
BLOOD
HEART
SPLEEN
SKM–GN
SKM–VL
WAT–SC
BAT
LIVER
LUNG
KIDNEY
ADRNL
HIPPOC
SMLINT
COLON
B cells memory
B cells naive
Plasma cells
T cells CD4 memory activated
T cells CD4 memory resting
T cells CD4 naive
T cells CD8
T cells follicular helper
T cells gamma delta
T cells regulatory (Tregs)
NK cells activated
NK cells resting
Eosinophils
Mast cells activated
Mast cells resting
Neutrophils
Dendritic cells activated
Dendritic cells resting
Macrophages M0
Macrophages M1
Macrophages M2
Monocytes

### Supplementary Figure 8. Assessment of immune responses via immunoassays, pathways, and cell types.

**A)** Heatmap of the training response of analytes measured in the immunoassays in each tissue. Color (white to red) indicates the $-\log_{10}$ adjusted p-value. Gray indicates the tissue was not assayed for the specific analyte. Top bars indicate the number of differential analytes in each tissue; right bars indicate the number of tissues in which the analyte is training-regulated (5% FDR). **B)** Heatmap of the percentage and number of immune-related pathways significantly enriched at 8 weeks in each tissue and ome. Pathways belonging to the immune system categories in the KEGG and Reactome databases were labeled as immune-related. The number of immune-related pathways out of the total number of significantly enriched pathways is represented for nodes with down-regulated or up-regulated features at 8 weeks of training. Results are separated by female-specific, male-specific, and sex-consistent nodes. **C)** Line plots of standardized abundances of training-differential proteins in subcutaneous white adipose tissue (WAT-SC) that are up-regulated only in males at the 8-week time point (8w_F0_M1). The black line in the center represents the average value across all features. **D)** Violin plots of the sample-level Pearson correlation between markers of immune cell types, lymphatic tissue, or cell proliferation and the average value of features in **(C)** at the protein level. A red point indicates that the marker is also one of the differential features plotted in **(C)**. # indicates when the distribution of Pearson correlations for a set of at least two markers is significantly different from 0 (one-sample t-test, 5% BY FDR). When only one marker is used to define a category on the y-axis, the gene name is provided in parentheses. **E)** Trajectories of various immune cell type signals in BAT or WAT following deconvolution of bulk RNA-Seq with CIBERSORTx. **F)** Immune cell type enrichment analysis results of training-differentially expressed transcripts using cell type specificity scores generated from the CIBERSORT LM22 gene signature matrix. Each point represents a significant enrichment (5% FDR, one-sided Mann-Whitney U test) of cell-type-specific genes in the set of transcripts contained in a particular non-null 8-week node of the graphical clustering results, e.g., a blue triangle pointing up indicates an enrichment in the 8w_F0_M1 node, which represents features up-regulated only in males at 8 weeks.

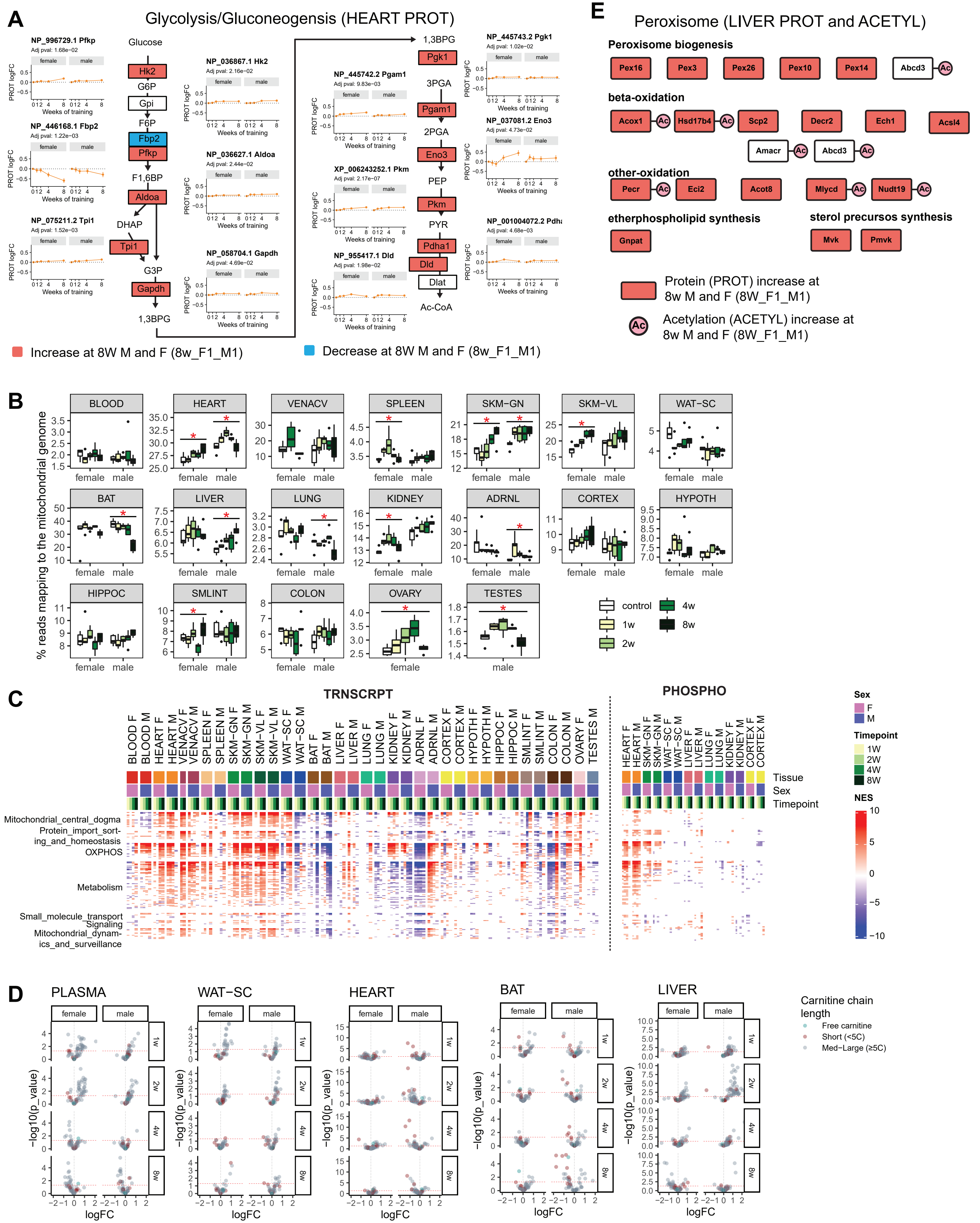
A
Glycolysis/Gluconeogensis (HEART PROT)
Glucose
Hk2
G6P
Gpi
F6P
Fbp2
Pfkp
F1,6BP
Aldoa
DHAP
Tpi1
G3P
Gapdh
1,3BPG
Pgk1
3PGA
Pgam1
2PGA
Eno3
PEP
Pkm
PYR
Pdha1
Dld
Dlat
Ac-CoA
NP_996729.1 Pfkp
NP_446168.1 Fbp2
NP_075211.2 Tpi1
NP_036867.1 Hk2
NP_036627.1 Aldoa
NP_058704.1 Gapdh
NP_445742.2 Pgam1
XP_006243252.1 Pkm
NP_955417.1 Dld
NP_445743.2 Pgk1
NP_037081.2 Eno3
NP_001004072.2 Pdha
Weeks of training
PROT logFC
Increase at 8W M and F (8w_F1_M1)
Decrease at 8W M and F (8w_F1_M1)
E
Peroxisome (LIVER PROT and ACETYL)
Peroxisome biogenesis
Pex16
Pex3
Pex26
Pex10
Pex14
Abcd3
beta-oxidation
Acox1
Hsd17b4
Scp2
Decr2
Ech1
Acsl4
Amacr
Abcd3
other-oxidation
Pecr
Eci2
Acot8
Mlycd
Nudt19
etherphospholipid synthesis
Gnpat
sterol precursos synthesis
Mvk
Pmvk
Protein (PROT) increase at 8w M and F (8W_F1_M1)
Acetylation (ACETYL) increase at 8w M and F (8W_F1_M1)
B
% reads mapping to the mitochondrial genome
BLOOD
HEART
VENACV
SPLEEN
SKM-GN
SKM-VL
WAT-SC
BAT
LIVER
LUNG
KIDNEY
ADRNL
CORTEX
HYPOTH
HIPPOC
SMLINT
COLON
OVARY
TESTES
female
male
control
1w
2w
4w
8w
C
TRNSCRPT
PHOSPHO
Tissue
Sex
Timepoint
Mitochondrial_central_dogma
Protein_import_sorting_and_homeostasis
OXPHOS
Metabolism
Small_molecule_transport
Signaling
Mitochondrial_dynamics_and_surveillance
Sex
F
M
Timepoint
1W
2W
4W
8W
NES
D
PLASMA
WAT-SC
HEART
BAT
LIVER
-log10(p_value)
logFC
Carnitine chain length
Free carnitine
Short (<5C)
Med-Large (≥5C)

### Supplementary Figure 9. Metabolic and mitochondrial effects of endurance training.

**A)** Diagram indicating protein abundance changes in the glycolysis and gluconeogenesis pathway in the heart tissue after 8 weeks of training. Line plots show the $\log_2$ fold-changes over the training time course (error bars indicate standard errors). Boxes colored in red indicate a statistically significant (5% FDR) increase in abundance for both males and females at 8 weeks, while blue indicates decreased abundance. **B)** Boxplots showing the percent of mitochondrial genome reads across tissue, sex, and time points. Red asterisks indicate a significant change throughout the training time course (F-test, 5% FDR). **C)** GSEA using the MitoCarta MitoPathways gene set database and transcriptome (TRNSCRPT) or phosphoproteome (PHOSPHO) differential analysis results. NES are shown for significant pathways (10% FDR) for all tissues, sexes, and time points within the heatmap. Mitochondria pathways are shown as rows and grouped using the parental group in the MitoPathways hierarchy. **D)** Volcano plots showing abundance changes ($\log_2$ fold-changes; logFC) and significance ($-\log_{10}$ nominal p-values) for acyl-carnitines. Features are colored based on the carnitine chain length as indicated in the legend. **E)** Diagram indicating protein abundance and protein acetylation level changes in the peroxisome KEGG pathway in the liver tissue after 8 weeks of training. Red boxes indicate an increase in abundance for both males and females, while red circles indicate an increase in at least one acetylsite within the protein (8w_F1_M1 cluster).
